# The Genetic Architecture of Dietary Iron Overload and Associated Pathology in Mice

**DOI:** 10.1101/2023.06.05.543764

**Authors:** Brie K. Fuqua, Lambda Moses, Stela McLachlan, Calvin Pan, Richard C. Davis, Simon T. Hui, Nam Che, Zhiqiang Zhou, Carmen Ng, Sarada Charugundla, Montgomery Blencowe, Zara Saleem, Aika Miikeda, Beyza Ozdemir, Chester Hui, Thy Li, Clara L. Stolin, Marianne Kozuch, Jie Zhou, Kathryn Page, Hiro Irimagawa, Nam Ku, Kodi Taraszka, Nathan LaPierre, David W. Killilea, David M. Frazer, Xia Yang, Eleazar Eskin, Chris D. Vulpe, Aldons J. Lusis

## Abstract

Tissue iron overload is a frequent pathologic finding in multiple disease states including non-alcoholic fatty liver disease (NAFLD), neurodegenerative disorders, cardiomyopathy, diabetes, and some forms of cancer. The role of iron, as a cause or consequence of disease progression and observed phenotypic manifestations, remains controversial. In addition, the impact of genetic variation on iron overload related phenotypes is unclear, and the identification of genetic modifiers is incomplete. Here, we used the Hybrid Mouse Diversity Panel (HMDP), consisting of over 100 genetically distinct mouse strains optimized for genome-wide association studies and systems genetics, to characterize the genetic architecture of dietary iron overload and pathology. Dietary iron overload was induced by feeding male mice (114 strains, 6-7 mice per strain on average) a high iron diet for six weeks, and then tissues were collected at 10-11 weeks of age. Liver metal levels and gene expression were measured by ICP-MS/ICP-AES and RNASeq, and lipids were measured by colorimetric assays. FaST-LMM was used for genetic mapping, and Metascape, WGCNA, and Mergeomics were used for pathway, module, and key driver bioinformatics analyses. Mice on the high iron diet accumulated iron in the liver, with a 6.5 fold difference across strain means. The iron loaded diet also led to a spectrum of copper deficiency and anemia, with liver copper levels highly positively correlated with red blood cell count, hemoglobin, and hematocrit. Hepatic steatosis of various severity was observed histologically, with 52.5 fold variation in triglyceride levels across the strains. Liver triglyceride and iron mapped most significantly to an overlapping locus on chromosome 7 that has not been previously associated with either trait. Based on network modeling, significant key drivers for both iron and triglyceride accumulation are involved in cholesterol biosynthesis and oxidative stress management. To make the full data set accessible and useable by others, we have made our data and analyses available on a resource website.

**Author summary:** The response to a high iron diet is determined in part by genetic factors. We now report the responses to such a diet in a diverse set of inbred strains of mice, known as the Hybrid Mouse Diversity Panel, that enables high resolution genetic mapping and systems genetics analyses. The levels of iron in the liver varied about >5 fold across the strains, with genetic variation explaining up to 74% of the variation in liver iron. Pathologies included copper deficiency, anemia, and fatty liver, with liver triglycerides varying over 50 fold among the strains. Genetic mapping and network modeling identified significant genetic loci and pathways underlying the response to diet.

## Introduction

Iron, with its powerful ability to gain and lose electrons, plays an essential role in many biological processes, including oxygen transport and storage (via hemoglobin and myoglobin), DNA synthesis (ribonucleotide reductase), energy production (electron transport chain heme and iron-sulfur containing proteins), and detoxification (e.g. by the cytochrome p450 enzymes) (Collins and Anderson 2012). In people, as well as most organisms, iron homeostasis is tightly managed to ensure adequate iron for biological function while preventing excess. However, when iron levels exceeds an organism’s regulatory capacity, cellular and tissue pathology can result. Oxidative stress, elicited by the rapid reaction of iron with cellular O_2*_ ^-^ and H_2_O_2_ to generate extremely reactive HO·, which can oxidize nucleic acids, lipids, sugars, and proteins and trigger oxidative chain reactions (Galaris, Barbouti et al. 2019), likely contributes to the observed pathophysiology.

Iron overload disorders, including primary iron-overload diseases such as hereditary hemochromatosis, and secondary iron-overload diseases including those resulting from transfusion, such as seen in individuals treated for β-thalassemia and sickle cell disease, are common genetic disorders worldwide (Taher, Weatherall et al. 2018; Xiao and Lauschke 2021). There is no regulated mechanism for the body to excrete iron, so excess iron acquired in these diseases accumulates, particularly in the liver, heart, pancreas, and endocrine glands (Kremastinos and Farmakis 2011; Anderson and Frazer 2017). Pathology associated with iron overload disorders includes chronic liver disease and cirrhosis, cardiomyopathy, dyslipidemia, arthritis, skin hyperpigmentation, and diabetes (van Bokhoven, van Deursen et al. 2011; Chacon, Morrison et al. 2013; Sengsuk, Tangvarasittichai et al. 2014; Anderson and Frazer 2017; Seeßle, Gan-Schreier et al. 2020; Kane, Roberts et al. 2021). Despite advances in detection and treatment, iron overload-related liver disease and iron overload cardiomyopathy remain major causes of morbidity and mortality in patients with primary and secondary iron overload (Gujja, Rosing et al. 2010; Kowdley, Brown et al. 2019; Hsu, Senussi et al. 2022). Abnormal tissue iron accumulation is also associated with other common diseases including fatty liver disease, cancer, and neurodegenerative diseases including Alzheimer’s, Huntington’s and Parkinson’s disease, but the etiology and implications of these associations are not well understood (Belaidi and Bush 2016; Britton, Subramaniam et al. 2016; Torti, Manz et al. 2018; Crawford, Ross et al. 2021; Fernandez, Lokan et al. 2022).

Key genes known to influence iron status in human populations have been identified and include those involved in iron absorption and trafficking (e.g. *TF*, *TFRC*, *SLC40A1*) and in the regulation of these processes (e.g. *HFE*, *HAMP*, *TFR2*, *TMPRSS6*, *HJV*). However, the penetrance of variants in these genes, most notably in *HFE*, one of the most common genetic contributors to iron overload, is incomplete and influenced by sex and other genetic and environmental factors (Anderson and Bardou-Jacquet 2021). Human genome-wide association studies (GWAS) have identified variants in known iron related genes, as well as novel loci, associated with markers of iron status (hematology, serum ferritin, serum iron, serum transferrin, and serum transferrin saturation) (McLaren, Garner et al. 2011; Benyamin, Esko et al. 2014; Liao, Shi et al. 2014; Li, Lange et al. 2015; Raffield, Louie et al. 2017; Bell, Rigas et al. 2021; Moksnes, Graham et al. 2022) and more recently, liver iron as measured by MRI in the UK Biobank Western European population (Wilman, Parisinos et al. 2019; Liu, Basty et al. 2021; O’Dushlaine, Germino et al. 2021). While some genes are shared between studies, new loci are still being identified as larger population sizes and a wider variety of ethnicities are examined.

In addition to the variation in iron status seen in populations, variation is also seen in the extent and type of pathology associated with iron overload with genetics playing an important role (El Beshlawy, El Tagui et al. 2014; Aydinok, Porter et al. 2015; Anderson and Bardou-Jacquet 2021; Buch, Sharma et al. 2021; Kang, Barad et al. 2021). Genes known to be involved in iron metabolism have also been identified in GWAS for traits linked to iron overload pathology, for example association of variants in *TFRC* with type 2 diabetes (Spracklen, Horikoshi et al. 2020; Vujkovic, Keaton et al. 2020), variants in *SLC39A8* and *TF* with HDL cholesterol (Richardson, Sanderson et al. 2020; Martin, Cule et al. 2021; Sakaue, Kanai et al. 2021; Sinnott-Armstrong, Tanigawa et al. 2021), and recent associations in the UK Biobank of *HFE*, *TMPRSS6*, and *SLC39A8* with MRI signals for liver inflammation and fibrosis (Parisinos, Wilman et al. 2020; O’Dushlaine, Germino et al. 2021).

In order to gain further insight into genetic contributors to iron overload and associated pathology, we have examined a genetically diverse cohort of mice known as the Hybrid Mouse Diversity Panel (HMDP). The panel has previously been used to examine numerous complex traits, including obesity, diabetes, atherosclerosis, heart failure, carbon tetrachloride induced liver fibrosis, and fatty liver disease (Lusis, Seldin et al. 2016; Seldin, Yang et al. 2019; Tuominen, Fuqua et al. 2021; Cao, Wang et al. 2022). Mice have important advantages for studies of the genetics of complex traits, including the ability to control the environment and access the relevant tissues. In the present study, 114 strains of mice were maintained on a 20k ppm (2% carbonyl iron) high iron diet for 6-7 weeks followed by analyses of pathology and molecular phenotypes.

## Results

### Pilot study liver metals

We first performed a pilot study with six HMDP strains (A/J, C3H/HeJ, DBA/2J, C57BL/6J, BALB/cJ, and AKR/J), the first five of which contribute much of the mapping power to the HMDP panel, to confirm that differences between strains in liver iron levels would be suitable for genetic mapping. Male mice were put on a 50 ppm iron sufficient control diet or a 20k ppm (20,000 ppm iron = 2% carbonyl iron) high iron diet for six weeks starting at four weeks of age (Figure 1A). Liver metals were measured by inductively coupled plasma atomic emission spectroscopy (ICP-AES). On the 50 ppm iron sufficient control and 20k ppm high iron diets, respectively, we observed a 3.3 and 1.9 fold difference between the strains with the highest and lowest liver iron levels (Figure 1B, 1C). On the 50 ppm iron sufficient control diet, mean strain liver iron levels ranged from approximately 270 ppm (C57BL/6J) to 900 ppm (AKR/J), and on the high iron diet, from 4550 ppm (A/J) to 8820 ppm (BALB/cJ). The 20k ppm high iron diet led to mean strain liver iron levels 6 to 30 times that of the 50 ppm iron sufficient control diet, with the greatest increase in the C57BL/6J strain (Figure 1D). For reference, normal human liver iron levels have been reported to range from 200-2000 ppm, with concentrations roughly above 5,000 ppm associated with adverse effects and above 15,000 ppm with cirrhosis (Fiel 2022). Human patients with hemochromatosis have up to 10 times more liver iron than the normal range, while patients with transfusion-dependent anemias can exceed 20 times the upper range of normal levels (Sirlin and Reeder 2010).

**Figure 1.**
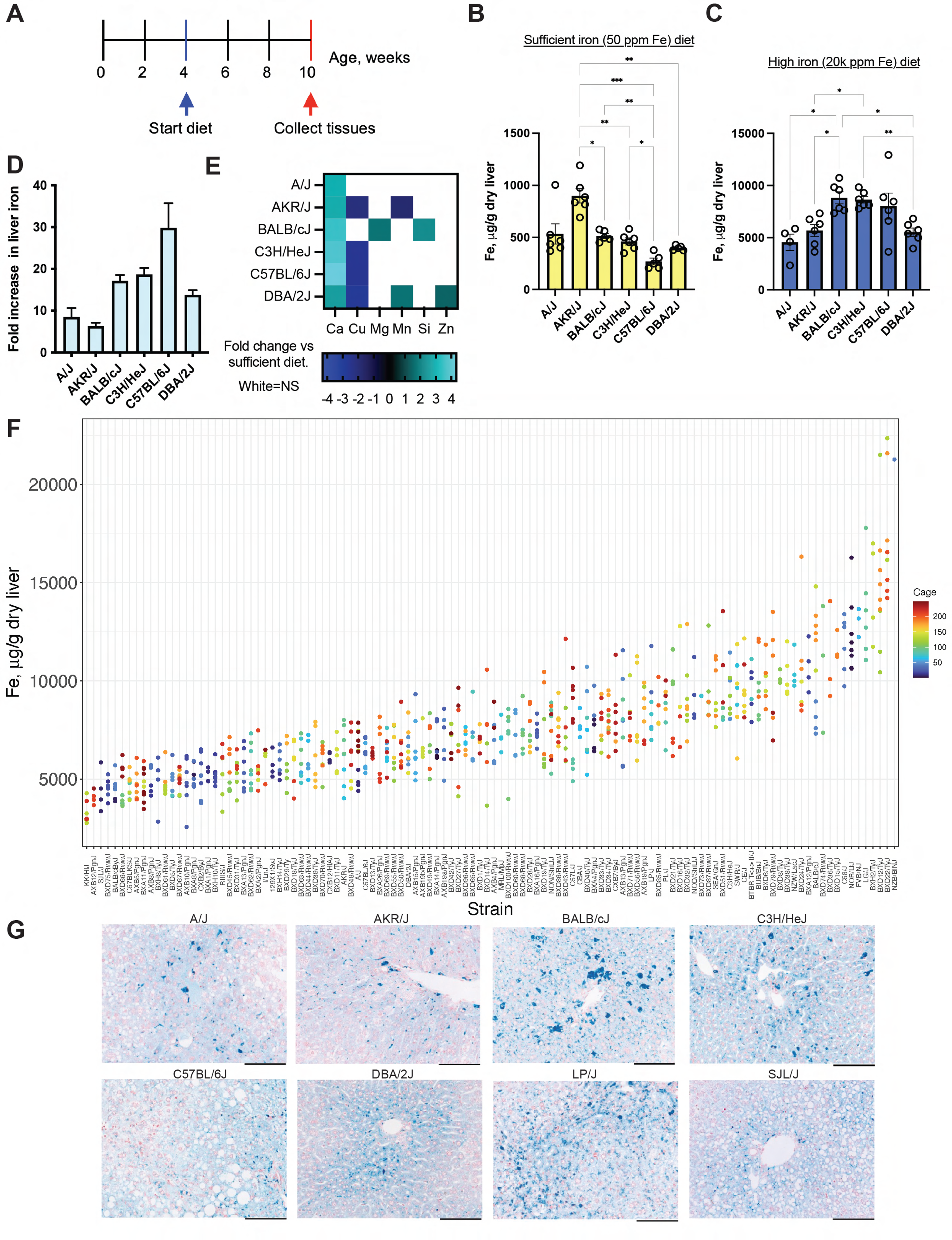
Six weeks of a high iron diet leads to variable iron loading across mouse strains. **(*A*)** Study schematic. For both the pilot study and the Control study, mice were switched from chow to a 50 ppm iron sufficient control diet or a 20k ppm (20,000 ppm iron = 2% carbonyl iron) high iron diet at 4 weeks of age. In the HMDP study, mice were switched from chow to a 20k ppm (20,000 ppm iron = 2% carbonyl iron) high iron diet at 4 weeks of age. Tissues were collected at 10-11 weeks of age following exsanguination and cardiac perfusion to remove blood. **(*B*)** Liver iron levels (µg iron per g dry liver weight) for the six pilot study strains fed a 50 ppm iron sufficient control diet. Mean and SEM. N=5-6 mice per strain. **(*C*)** Liver iron levels (µg iron per g dry liver weight) for six pilot study strains fed the 20k ppm high iron diet. Mean and SEM. N=4-6 mice per strain. For B and C, differences between strains that were significant (by one-way Brown-Forsythe and Welch’s ANOVA test with Dunnett’s T3 multiple comparisons test in GraphPad Prism v9) are denoted with \**P* ≤ 0.05; \*\**P* ≤ 0.01; \*\*\**P* ≤ 0.001. **(*D*)** Fold increase in liver iron in the six pilot study strains, calculated as the strain mean on the high iron diet minus the strain mean on the sufficient diet; error bars show SEM. **(*E*)** Significant fold changes in other liver metals, calculated as the strain mean on the high iron diet minus the strain mean on the sufficient diet. Changes that were <u>not</u> significant (*P* > 0.05 using unpaired t tests with Welch’s correction and the Holm-Šídák multiple comparisons test for each metal in GraphPad Prism v9) are in white. N=4-6 mice per strain per diet. **(*F*)** Individual value plot of liver iron (N=1-11 mice per strain) for the 114 HMDP strains fed the 20k ppm high iron diet, ordered by increasing strain mean. The color of each dot represents the cage number for a given mouse. Cages were sequentially ordered, so dots with similar colors indicate mice housed in the vivarium around the same dates. **(*G*)** Perls’ stained liver sections from eight strains, counterstained with Neutral Red. Iron stains blue. The snapshots shown are from whole-slide images scanned with the Aperio ScanScope AT at 200x magnification, and scale bars are 100μm. Source data for panels B-E are in Figure 1-Source Data 1. This file indicates and describes any data excluded from the analysis. Data for all the pilot study metals are visualized in Figure 1-figure supplement 1. Source data for panel F is Figure 1-Source Data 2.

Other metals were also examined in the liver. In addition to inter-strain variation in metal concentrations, we also observed several metals that were impacted by the high iron diet (Figure 1E, Figure 1-figure supplement 1, Figure 1-Source Data 1). Notably, mean liver calcium levels were 2.4 to 4.3 times higher in strains fed the 20k ppm high iron diet, and mean liver copper levels were reduced in 3 strains (C3H/HeJ, C57BL/6J, and DBA/2J) to less than half of levels on the 50 ppm iron sufficient control diet. Liver magnesium, silicon, and zinc were higher in some strains, while liver manganese was lower in AKR/J and higher in DBA/2J, in the iron loaded mice compared to those fed the 50 ppm iron sufficient control diet. The inter-strain variation in liver metal levels suggests that genetic differences contribute to the variation in these traits, so we proceeded with a systems genetics analysis of the full HMDP.

### HMDP study liver metals

A total of 746 male mice from 114 HMDP strains were fed a 20k ppm (20,000 ppm iron = 2% carbonyl iron) high iron diet for 6-7 weeks, starting at four weeks of age, to induce iron overload (Figure 1A, Supplementary File 1). Twenty-one mice (from 12 strains) not included in this number and excluded from further study died prematurely or were euthanized early due to fight wounds, hydrocephalus, sickly appearance, or rectal prolapse. The 114 strains consisted of 31 Classical Inbred (CI) strains and 83 Recombinant Inbred (RI) strains (10 AXB, 10 BXA, 56 BXD, 4 BXH, and 3 CXB). Liver metals were measured by inductively coupled plasma mass spectrometry (ICP-MS). There was a 6.5 fold variation in mean liver iron concentration across all 114 HMDP strains, ranging from less than 4,000 ppm to over 20,000 ppm (Figure 1F, Figure 1 - Source Data 2, Supplementary File 2). The estimated fraction of variance in liver iron levels that could be attributed to genetics (heritability) was high, with a broad sense heritability of 0.74 and a narrow sense heritability of 0.73 (Supplementary File 3). Heritability for non-heme liver iron was previously estimated to be 0.62 in a C57BL/6J HFE^-/-^ X DBA/2J HFE^-/-^ mixed sex F2 intercross (Bensaid, Fruchon et al. 2004). Variation was also seen in the concentration of other liver metals across the HMDP, with a notable 9.6 fold variation in both liver manganese and copper (Supplementary File 2 and web resource). Perls’ stain to visualize ferric iron deposits in the liver revealed diffuse and punctate staining in hepatocytes, with most staining near the portal vein regions (Figure 1G). Variation was seen visually between strains in the extent and pattern of iron loading.

### C57BL/6J comparative diet study liver metals

To gain insight into which phenotypes were influenced by the high iron diet, and to quantify the extent to which these phenotypes were influenced by the diet, we also did a study (“Control study”) in just one mouse strain, C57BL/6J, fed one of four diets (a chow diet, and three diets only differing in iron content: a 50 ppm FeSO_4_ iron sufficient control diet, a 500 ppm FeSO_4_ elevated iron diet, and a 20k ppm (20,000 ppm iron = 2% carbonyl iron) high iron diet for 6-7 weeks starting at 4 weeks of age. While the pilot study (performed before the Control study or full HMDP study) gave us early insight into phenotypes affected by a high iron diet, the pilot study diets were inadvertently not isocaloric, complicating fair comparisons for some traits of interest (e.g. lipid traits). The Control study also allowed us to collect some additional sample types for analysis that were not collected in the pilot study. In the Control study, as for the HMDP, liver metals were measured by ICP-MS. The chow, 50 ppm iron sufficient control diet, 500 ppm elevated iron diet, and 20k ppm high iron diet led to liver iron concentrations (mean and SEM) of 123 ± 4, 234 ± 9, 390 ± 14, and 5,860 ± 301 μg iron/g dry tissue, respectively, with a 25 fold increase in liver iron between the control and high iron diet groups. The concentration of other metals were also affected by the diets (Supplementary File 4, Supplementary File 5). While the 500 ppm elevated iron diet significantly but modestly increased liver iron levels compared to the 50 ppm iron sufficient control diet, a slight decrease in manganese was the only other difference between these two groups. The 20k ppm high iron diet, however, led to decreases in copper, zinc, magnesium, selenium, cobalt, rubidium, and strontium compared to the 50 ppm iron sufficient control diet. There is incomplete overlap in the types of metals that we could measure in the pilot study with ICP-AES and in the Control study with ICP-MS, but of the metals that overlapped, copper was significantly decreased in both studies in C57BL/6J in the high iron diet versus the 50 ppm iron sufficient control diet, while zinc and magnesium were significantly decreased in the Control study on the 20k ppm high iron diet but were not different in the pilot study in this strain compared to the 50 ppm iron sufficient control diet. The chow diet, which is a crude diet made from variable ingredients, led to higher levels of several metals than the other diets, most notably manganese, cobalt, rubidium, and cadmium.

### Additional phenotypes affected by the high iron diet

#### Liver and plasma lipids

We observed a spectrum of what appeared to be macrovesicular steatosis when examining the Perls’ stained and hematoxylin and eosin (H&E) stained sections across the HMDP (Figure 1G, Figure 2A). Staining of frozen sections of liver from the C57BL/6J Control study mice with Oil Red O confirmed that the droplets observed in the mice fed the 20k ppm high iron diet contained neutral lipids (Figure 2B). Large droplet macrovesicular steatosis was observed to varying extents in six out of the eight livers in the 20k ppm high iron diet group but not in the 50 ppm iron sufficient control diet or 500 ppm elevated iron diet groups. Liver triglyceride (TG) levels for the C57BL/6J mice fed the 20k ppm high iron diet, as measured by colorimetric assay, were not significantly different than those from the 50 ppm iron sufficient control diet (*P* = 0.0835), but they trended toward being higher (Figure 2C, Figure 2-Source Data 1). Liver total cholesterol (TC), esterified cholesterol (EC), and phospholipids-C (PL) were significantly altered by the 20k ppm high iron diet compared to control, with liver TC and EC increasing to 1.2 and 1.8 times that of control, and liver PL falling slightly to 91% that of control. Liver unesterified cholesterol (UC) was not affected by the diet, so the increase in liver TC is due to an increase in EC. Furthermore, the 20k ppm high iron diet led to alterations in plasma lipids in this strain, with a 3 fold increase in plasma TG and a 22-35% decrease in plasma TC, UC, EC, high-density lipoprotein (HDL) cholesterol, and low and very low density lipoprotein (LDL&VLDL) cholesterol (Figure 2D, Figure 2-Source Data 1). The percent of plasma TC in the esterified form decreased about 5% versus control. Plasma insulin levels and plasma glucose levels were not significantly altered. We did not detect evidence of liver fibrosis by picrosirius red staining of collagen fibers when we analyzed a subset of strains (A/J, AKR/J, BALB/cJ, C57BL/6J, DBA/2J, C3H/HeJ, BTBR T<+> tf/J, LP/J, and SJL/J) (data not shown).

**Figure 2.**
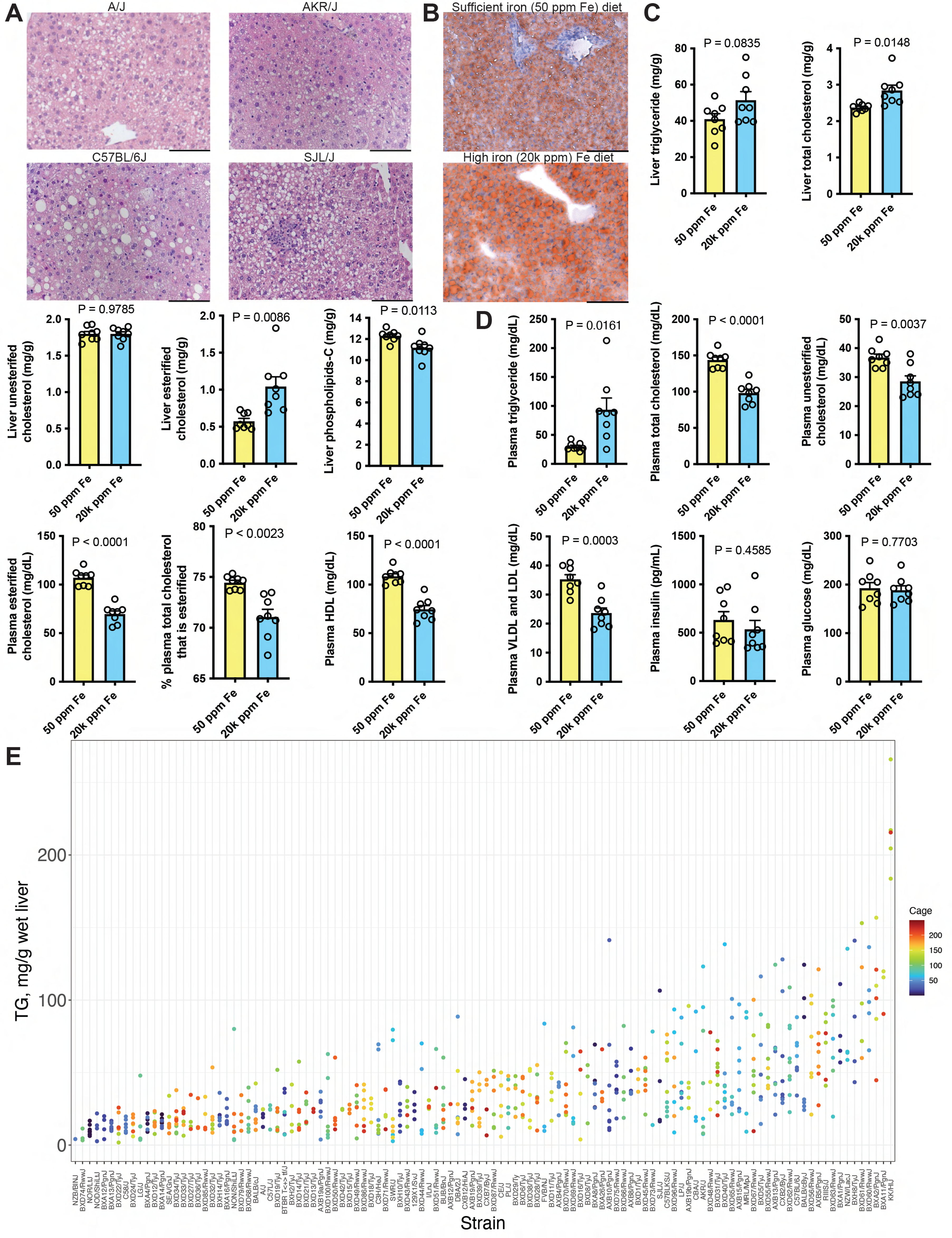
Response of liver and plasma lipids to the high iron diet. **(*A*)** H&E stained liver sections from four mouse strains from the HDMP study with regions that appear to contain small and/or large macrovesicular lipid droplets. **(*B*)** Oil Red O stained sections from frozen embedded liver from Control study C57BL/6J mice fed the 50 ppm iron sufficient control diet (top) or 20k ppm high iron diet (bottom). Neutral lipid stains red and nuclei stain pale blue. For A and B, images are at 200x magnification, and scale bars are 100μm. **(*C*)** Liver lipids as measured by colorimetric assay in the 50 ppm iron sufficient control diet fed and 20k ppm high iron fed Control study C57BL/6J mice. Mean and SEM, N=8 mice per diet. **(*D*)** Plasma lipids, glucose, and insulin in the 50 ppm iron sufficient control diet fed and 20k ppm high iron fed Control study C57BL/6J mice. Mean and SEM, N=8 mice per diet. For C and D, *P* values were calculated by t-test with Welch’s correction in Graphpad Prism v9. **(*E*)** Individual value plot of liver triglyceride (N=1-10 mice per strain) for the 114 HMDP strains fed the 20k ppm high iron diet, ordered by increasing strain mean. The color of each dot represents the cage number for a given mouse. Cages were sequentially ordered, so dots with similar colors indicate mice housed in the vivarium around the same dates.

In order to gain more insight into genetic factors that may influence lipid levels on the high iron diet, we extracted and quantified liver lipids (TG, TC, UC, EC, and PL) and plasma lipids (TG, TC, UC, EC, HDL cholesterol, and LDL&VLDL cholesterol) in the 114 iron loaded HMDP strains (Supplementary File 2 and web resource). We observed approximately 50 fold variation in mean liver TG levels across the strains, ranging from ∼4 to 217 mg/g wet liver (Figure 2E, Figure 2-Source Data 2). The variation in liver TC, PL, and UC was more modest, with 5.5, 2.4, and 2.6 fold variation across strains, respectively, while EC varied 20.1 fold (Supplementary File 6 and Supplementary File 7). For plasma lipids, we observed 28 fold variation in TG, 16-18 fold variation in EC and LDL&VLDL cholesterol, 11 fold variation in HDL cholesterol, and 6-7 fold variation in plasma TC and UC (Supplementary File 2). Broad sense heritability ranged from 0.38 to 0.53 for liver lipids and from 0.42 to 0.60 for plasma lipids, while narrow sense heritability ranged from 0.26 to 0.47 for liver lipids and from 0.22 to 0.50 for plasma lipids (Supplementary File 3).

#### Anemia

We also observed a spectrum of pallor and anemia in HMDP strains on the high iron diet, with many mice with red cell count and hemoglobin levels below normal reference ranges (7-13 x10^6^ red cells/µL and 10-17 g hemoglobin/dl) (McGarry, Protheroe et al. 2010; Fox 2014) (Supplementary File 8, Supplementary File 2, and web resource). This anemia has been previously reported by others, and while the molecular mechanism(s) leading to it are not known, it is likely due to copper deficiency (Wang, Xiang et al. 2018). This hypothesis is consistent with the lower liver copper levels observed in the pilot study for all strains on the high iron diet (Figure 1-figure supplement 1). In the pilot study, the red cell traits of some strains were much more affected by the high iron diet than other strains. Red blood cell count (RBC), hematocrit (HCT), and hemoglobin (HGB) were markedly lower in mice fed the 20k ppm high iron diet versus the 50 ppm iron sufficient control diet for the C3H/HeJ, C57BL/6J, and DBA/2J strains, and trended toward being lower for the AKR/J strain (Supplementary File 9, Supplementary File 10). These four strains had significantly decreased copper in response to the high iron diet in the pilot study. In contrast, A/J and BALB/cJ showed little change in these red cell parameters or in liver copper levels in the pilot study. In concordance with this, in the full HMDP study on the high iron diet, these two strains were also at the upper end of the spectrum for these traits. Mean cell volume (MCV) and mean cell hemoglobin (MCH) were also significantly higher in the C3H/HeJ, C57BL/6J, and DBA/2J strains on the high iron diet versus sufficient, while mean cell hemoglobin concentration (MCHC) was decreased in DBA/2J. Red cell distribution width (RDW) and RDW absolute (RDWa) were higher on the high iron diet for most strains. Platelet count trended toward being higher in C3H/HeJ, C57BL/6J, DBA/2J, and AKR/J. Total white blood cell count and the lymphocyte count subset were significantly lower in BALB/cJ on the high iron diet versus sufficient (Supplementary File 10 and Supplementary File 11). While some trends were apparent, other white cell parameters were not significantly different between the diets.

In the Control study with just the C57BL/6J strain, complete blood count results for the 50 ppm iron sufficient control diet and 20k ppm high iron diet were similar to those seen for this strain in the pilot study (Supplementary File 12, Supplementary File 5). Examination of H&E stained femurs of the Control study mice on the 20k ppm high iron diet revealed hypercellular marrow with increased erythroid production. Wright-Giemsa stained blood smears showed severe anemia with increased size variation (high RDW) and increased polychromasia (reticulocytes). Red blood cells appeared macrocytic but otherwise normal. There were few red cell fragments, not indicative of frank hemolysis. There was clear evidence of extramedullary erythropoiesis in all H&E stained spleens and most liver sections, with livers with more prominent steatosis exhibiting less obvious extramedullary erythropoiesis. Blood smears and histology can be viewed on the resource website. Overall, these findings suggest that, in some strains, the high iron diet leads to a normocytic to macrocytic anemia potentially due to ineffective erythropoiesis related to copper deficiency. Acute blood loss or hemolysis could also be contributors in some strains, as we noted blood in the stool of some mice.

#### Body weight and food consumption

The iron loaded diet disrupted weight gain in some HMDP strains, and thirty percent of the strains had decreased mean weights at the end of the study period (Supplementary File 13, Supplementary File 2 and web resource). Mice from some strains had a rapid decline in weight in the final days of the study period. Average daily food consumption (by cage) over the study period and average mouse body weight (by cage) at the end of the study were highly correlated (bicor = 0.80, *P* = 2.81e-55) (Supplementary File 13, Supplementary File 2 and web resource). In the Control study, C57BL/6J mice consumed more food on the chow diet than the isocaloric study diets, for an average of 3.19 ± 0.08 g/day. C57BL/6J mice on the isocaloric diets differing only in iron consumed similar amounts to each other, with a trend toward slightly less food consumption of the high iron diet : 2.73 ± 0.09 g/day on the 50 ppm iron sufficient control diet, 2.73 ± 0.07 g/day on the 500 ppm elevated iron diet, and 2.60 ± 0.08 g/day on the 20k ppm high iron diet. C57BL/6J mice on the 20k ppm high iron diet weighed on average 8 to 10% less than mice on the 50 ppm and 500 ppm iron diets by the end of the study period. While mice on all diets overall gained lean mass over the study period, mice on the 20k ppm high iron diet did not significantly accumulate fat mass (Supplementary File 13, Supplementary File 5). Liver and heart weights were significantly greater, and full cecum weights (tissue and contents) significantly lower, in mice fed the 20k ppm high iron diet compared to control (Supplementary File 14, Supplementary File 5). Tibia length, left kidney weight, and spleen weights were not significantly different between the 20k ppm high iron diet and control group. Blood glucose area under the curve (AUC) was not different between groups in a glucose tolerance test (GTT) performed at 3-4 weeks on the diet, but GTT baseline fasting blood glucose was elevated in the 20k ppm high iron group compared to control. This elevated fasting blood glucose was similar to what was observed at the final Control study endpoint prior to euthanasia when blood glucose was also measured by a glucometer (Supplementary File 5). We did not see differences, however, in plasma glucose at the study endpoint as measured by enzymatic assay (Figure 2D). This discrepancy may be explained by the anemia in the 20k ppm high iron group that can artifactually raise readings taken with a glucometer (Barreau and Buttery 1987).

In summary, across the HMDP, the high iron diet led a spectrum of liver iron loading, hepatic steatosis, anemia, and disrupted weight gain. Furthermore, in the Control study in the C57BL/6J strain, we confirmed that, compared to the 50 ppm iron sufficient control diet, the 20k ppm high iron diet induced these phenotypes as well as led to decreased liver copper, alterations in plasma and liver lipids, disruption in the accumulation of fat mass, and increased liver and heart weight. Data for all traits can be further explored and plotted on the resource webpage.

#### HMDP trait by trait correlations

To understand how these phenotypes might be related to each other, we performed two different analyses to identify relationships between measured traits: hierarchical clustering and principal component analysis (PCA). We used hierarchical clustering to group clinical traits by the correlation between their strain means (Figure 3A, Figure 3-Source Data 1). Biweight midcorrelation (bicor) was used since it is a non-parametric, median-based correlation method less sensitive to outliers. Liver iron was negatively correlated with many traits including plasma EC, TC, and HDL cholesterol, red cell traits HCT, RBC, and HGB, plasma glucose, insulin and homeostatic model assessment of insulin resistance (HOMA-IR), body weight, fat mass, liver TG, and tissue weights except for spleen. Liver copper was positively correlated with plasma HDL cholesterol and red cell traits HCT, RBC, and HGB (Figure 3B), and negatively correlated with liver TC, spleen weight, RDW, RDWa, mean platelet volume (MPV), and MCH. The negative correlation of HCT, RBC, and HGB with RDW, RDWa, MPV, spleen and heart weight, and MCH is consistent with the anemia observed. Some other clusters of positively correlated traits included liver PL, liver UC, and most liver metals with the notable exception of copper; liver weight, kidney weight, week 0 and final lean mass, week 0 and final total mass, week 0 fat mass, and average daily food consumption; and HOMA-IR, plasma glucose, insulin, HDL cholesterol, EC, TC, final fat mass and change in fat and total mass.

**Figure 3.**
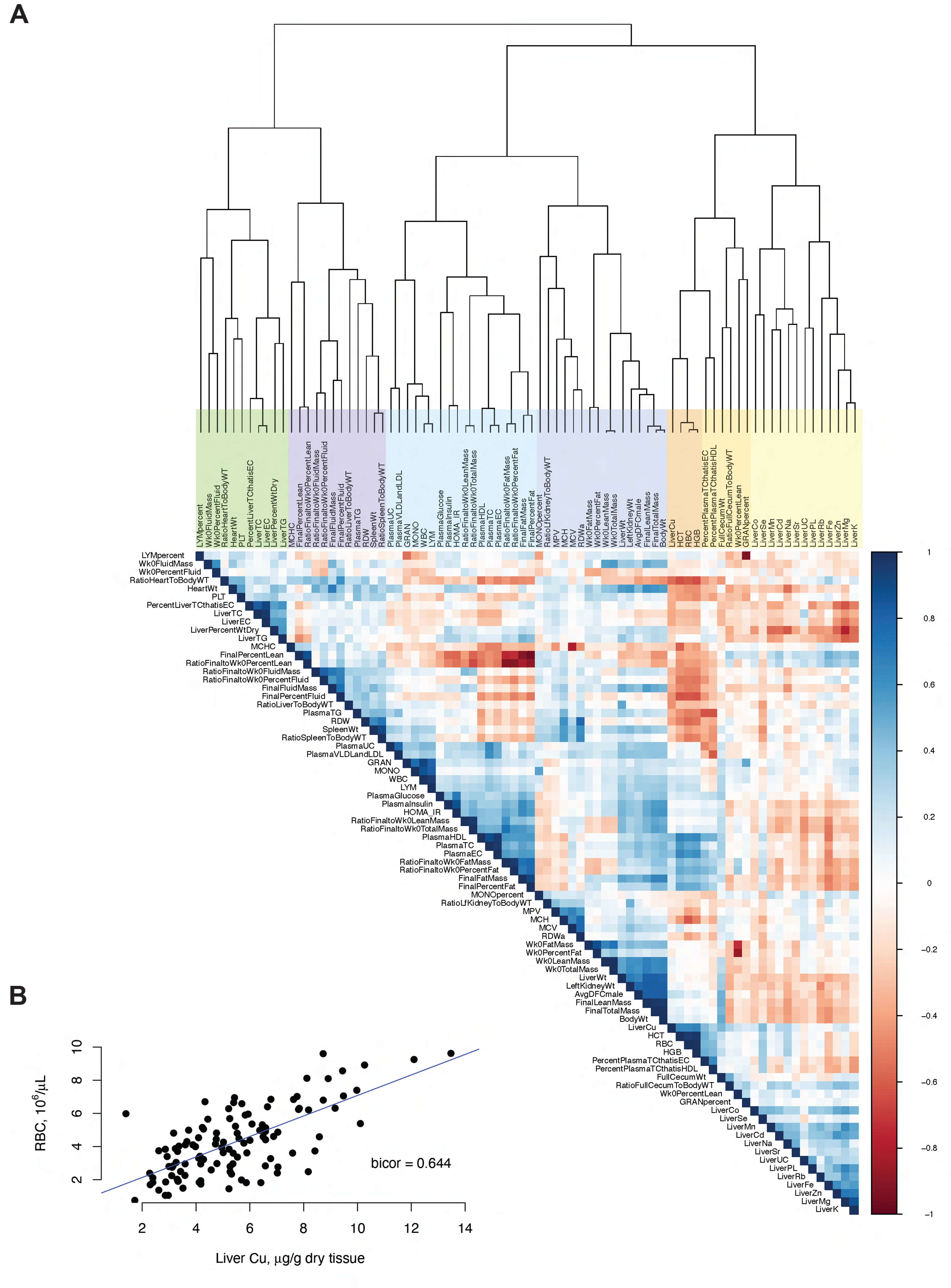
Pairwise correlation between traits across the HMDP. **(*A*)** Hierarchical clustered heatmap showing the correlation between eighty traits, with positive correlations in blue and negative correlations in red. Correlation was calculated across strain trait means using the bicor function in the WGCNA R package, using pairwise complete observations with a maxPOutliers setting of 0.05. The dendrogram on top shows how the traits clustered based on similarities in their correlations with other traits. Clusters of correlated traits are visible in the heatmap. A more detailed description of each of the traits is in Figure 3-Source Data 1. **(*B*)** Plot of red blood cell count versus liver copper concentration, with each dot representing the mean values for one mouse strain and the linear regression line shown.

We performed PCA using the strain means for 41 measured clinical traits that had data for all 114 strains. The first 5 principal components (PC) explain 68.93% of the variance. From the PCA loadings (Figure 4-Source Data 1) and seen visually on the loadings plot of PC1 and PC2 (Figure 4), PC1 mostly separated strains by body weight and composition, plasma cholesterol, and liver metals (especially iron), with liver iron negatively associated on this axis with final fat mass and plasma total and HDL cholesterol. PC2 was driven primarily by factors associated with the anemia seen in many strains (spleen and heart weight, fluid mass, red cell traits, liver copper). PC3 correlated with liver TG, white blood cell counts, and liver magnesium and potassium, while PC4 correlated with traits associated with red cell size (MCV, MCHC, RDWa, MCH). PC5 was most highly correlated with liver TC. The PCA plot of PC1 and PC2 (Figure 4-figure supplement 1), or any other plots of the first 5 PC combinations, do not reveal any tight distinct strain groupings. Figure 4-Source Data 2 shows the contribution of each strain to each PC.

**Figure 4.**
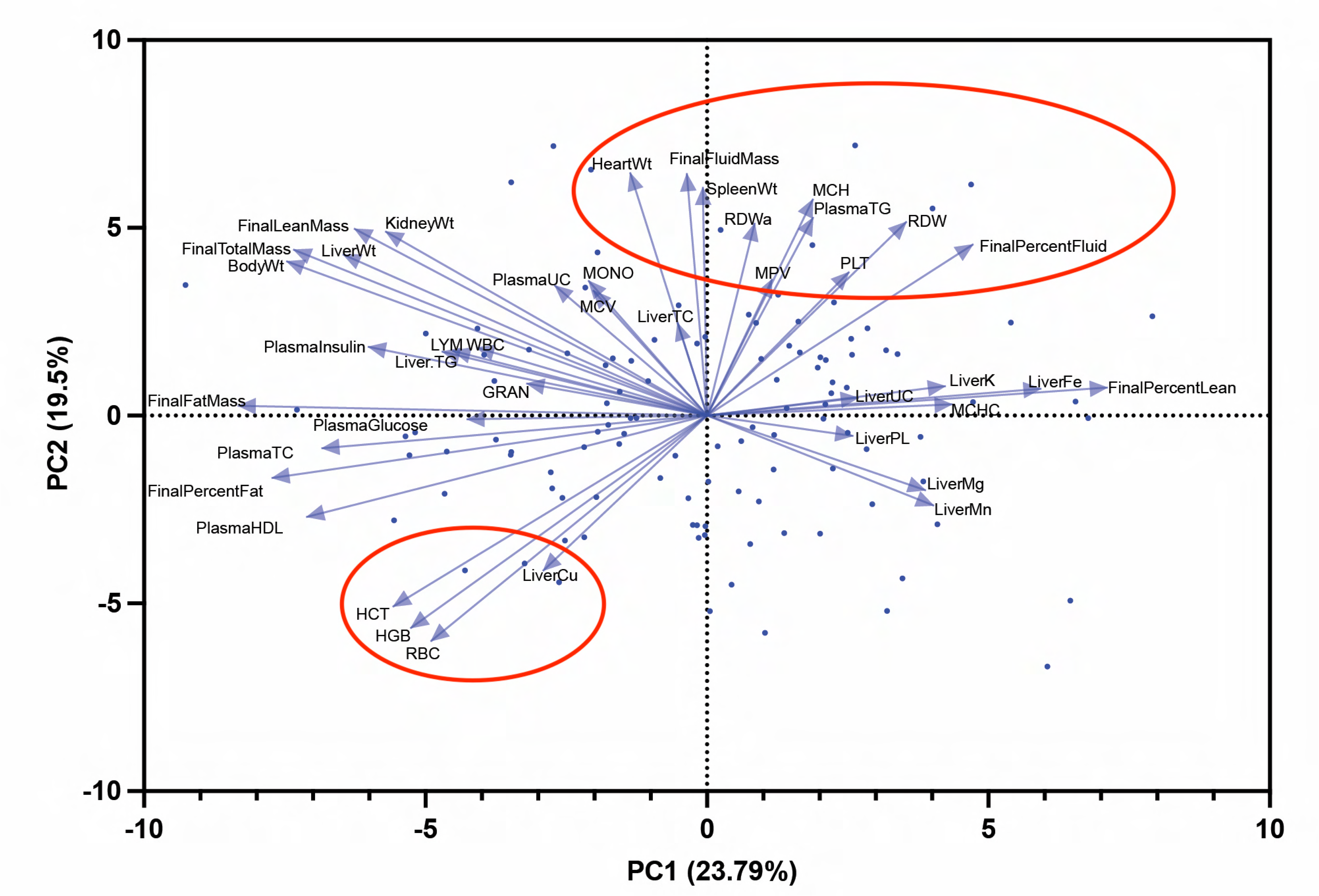
Principal component analysis (PCA) of 41 traits that had strain mean data for all 114 strains. PCA loadings plot showing the relationship between principal components PC1 and PC2 and the traits analyzed, with PC1 and PC2 explaining 23.79% and 19.5% of the variance, respectively. Traits with longer blue arrows and more parallel to a PC axis make a stronger contribution to that PC. A positive PC coordinate for a trait indicates that the trait is positively correlated with the PC, while a negative coordinate shows a negative correlation. Unlabeled blue dots represent the mouse strain PCA scores, which are labeled in Figure 4-figure supplement 1. The red circles highlight two clusters of traits associated with anemia. Trait means were z-score standardized as part of the analysis in GraphPad Prismv9. A more detailed description of each of the traits is in Figure 4-Source Data 1, and the contribution of each strain to each PC is in Figure 4-Source Data 2.

As expected, PCA trait vector groupings agreed well with those identified by hierarchical clustering of trait correlations. The PCA loadings plot (Figure 4) in particular emphasizes the strong correlation between liver copper and red cell traits, and their strong negative correlation with percent fluid mass (perhaps reflecting increased liquid blood volume in anemia), RDW (likely reflecting increased red cell synthesis in anemia), and platelet count (often elevated in anemia).

#### Genome-wide association analyses

To identify genetic loci that contribute to traits, we performed GWAS on the HMDP trait data. Significant quantitative trait loci (QTL) were identified for most traits examined (Supplementary Files 15-23). Loci for all GWAS can best be visualized, and top candidate genes explored, using the resource webpage. Below, we highlight some of the interesting QTL for liver iron, liver TG, red cell traits related to the anemia, and liver copper. Some additional QTL are discussed in Supplementary File 24.

#### Liver iron and TG QTL

A locus on chromosome 7 at 4.3-4.9 megabases (Mb) was strongly associated with liver iron levels (Figure 5A, Figure 5-Source Data 1) and overlapped with a slightly broader locus (4.3-5.1 Mb) for liver TG (Figure 5B, Figure 5-Source Data 2). This locus was the most significant locus for both of these traits, and the only significant locus for liver TG. This locus was also significantly associated with total liver weight and other liver metals including copper, potassium, magnesium, manganese, and rubidium, but not with zinc, cobalt, cadmium, sodium, selenium, or strontium (Supplementary Files 22-23, web resource). Linkage disequilibrium (LD) analysis shows that the significant SNPs in this locus for liver iron separate into two interspersed LD blocks (Figure 5A, top right plot). The most significant “top tier” SNPs for liver iron (6 in total, two visually overlapping, with -log_10_*P* values around 17) belong to one LD block (with the exception of the grey SNP, which has no LD information), while those in the “second tier” of significance (with -log_10_*P* values around 10) belong to another. The SNPs in the liver iron GWAS second tier (tagged in italics by SNP rs31801610) overlap with the most significant SNPs for liver TG (Figure 5B, bottom right plot), while the top tier SNPs for iron (tagged by SNP rs46388302) do not. Genotype analysis shows that the allele variation in the top tier significant SNPs for liver iron is only present in the CI strains, while the second tier significant SNPs have allele variation in both the CI and RI strains.

**Figure 5.**
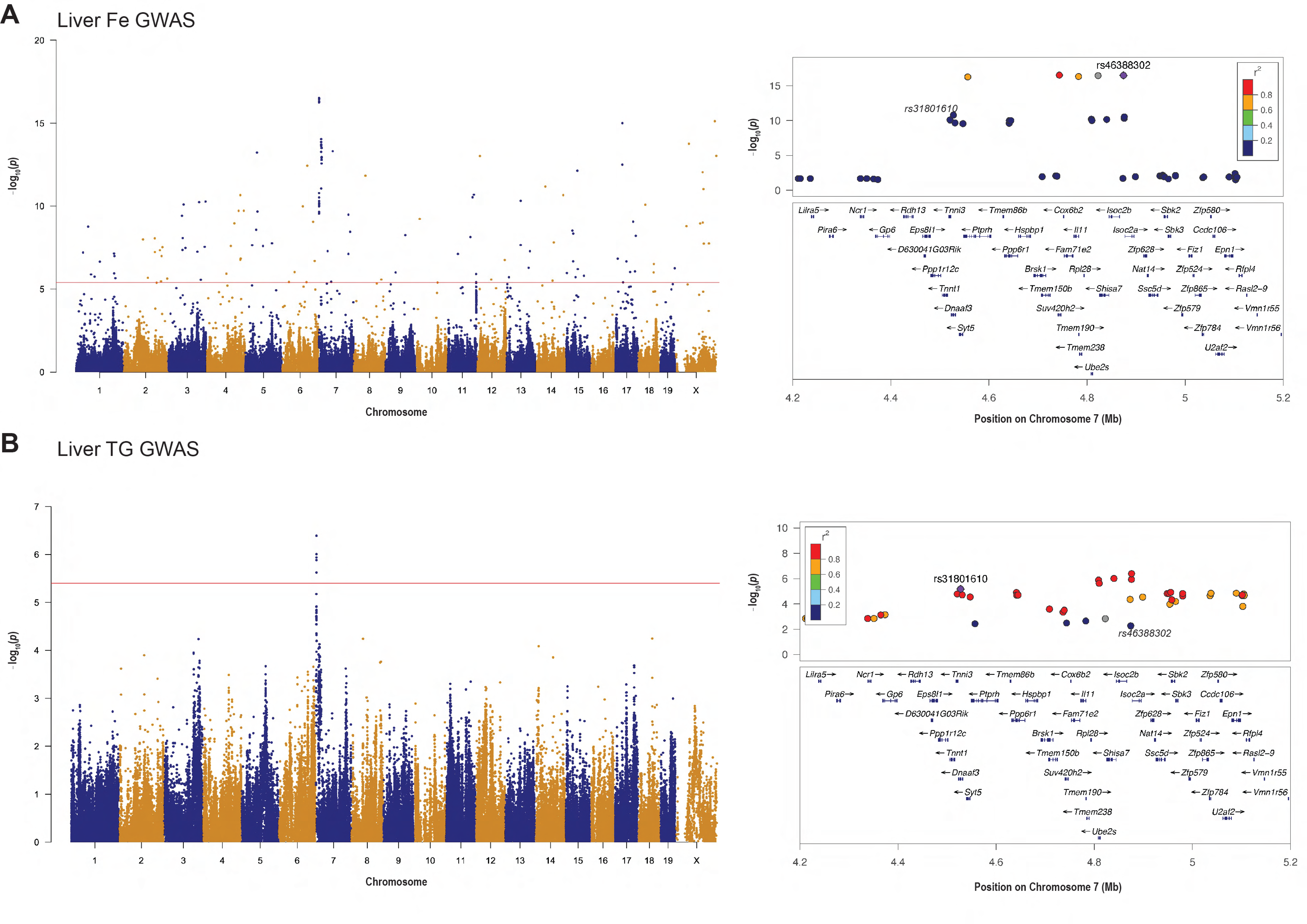
Overlapping loci associated with liver iron and liver triglyceride (TG) identified by genome-wide association studies (GWAS). **(*A*)** Manhattan plot (left) and LocusZoom plot of the overlapping chromosome 7 locus (right) from the liver iron GWAS. **(*B*)** Manhattan plot (left) and LocusZoom plot of the overlapping chromosome 7 locus (right) from the liver TG GWAS. For the Manhattan plots in A and B, loci with SNPs above the red bar (-log_10_*P* > 5.387) are statistically significant, while loci above -log_10_*P* > 4.387 are suggestive. In the LocusZoom plots, the r^2^ measure of linkage disequilibrium (LD) with the labeled SNP in purple (rs46388302 for the liver iron plot and rs31801610 for the liver TG plot) is indicated by the color of the SNP, as shown in the key. The liver iron labeled SNP is also labeled in italics on the TG locus zoom plot, and vice versa, for reference. Of note, the orange SNP on the top left of the LocusZoom plot for liver iron is actually made up of two side by side SNPs, so there are a total of six “top tier” SNPs, with five in strong LD (SNPs colored in grey, like one of these top tier SNPs, do not have LD data). Source data for the Manhattan plots are in Figure 5-Source Data 1 and Figure 5-Source Data 2.

Additional significant loci for liver iron include two significant loci between 13-15 Mb on chromosome 7 (Supplementary File 25). Significant (copper, liver TC, liver weight, magnesium, rubidium) and suggestive (potassium, manganese) loci in this region were also present for several traits. Conditioning on the top tier SNPs at the chromosome 7 4.3-4.9 Mb locus also removes these loci at 13-15 Mb in the GWAS for liver iron, suggesting potential long range linkage disequilibrium between these regions. Three other significant loci for liver iron are on distal chromosome 11 from 119-121 Mb with peak SNP rs27030299 at 119,387,437 bp (Supplementary File 25), on chromosome 6 from 29-30.5 Mb with peak SNP rs29657238 at 30,119,855 bp, and on chromosome 7 (peak SNP rs32227882 at 54,590,627 bp). The latter locus, near *Luzp2*, overlaps a previously described QTL for an eigentrait of transferrin saturation and plasma iron in a mouse BXD study (Jones, Beard et al. 2007). Some additional suggestive loci for liver iron include those on chromosome 1 (peak SNP rs49311170 at 153,571,983 bp), chromosome 3 (peak SNP rs47299487 at 122,856,962 bp), chromosome 4 (peak SNP rs28166315 at 105,919263 bp), chromosome 6 (peak SNP rs32013781 at 138,161,899 bp), chromosome 8 (peak SNP rs31663992 at 121,786,281 bp), chromosome 11 (peak SNP rs26946232 at 57,039,512 bp), chromosome 15 (peak SNP rs31773872 at 31,009,190 bp and peak SNP rs32357530 at 41,232,942 bp), and chromosome 19 (peak SNP rs30994468 at 47,275,146 bp). The loci on chromosome 1 and chromosome 3 may overlap with suggestive loci associated with increased non-heme liver iron in an HFE^-/-^ C57BL/6J and DBA/2J F2 intercross near microsatellite markers D1Mit206 and D3Mit32 (Bensaid, Fruchon et al. 2004). Syntenic genomic regions in humans near the suggestive chromosome 15 loci in mouse harbor suggestive loci for transferrin levels in a human meta-analysis GWAS (Benyamin, Esko et al. 2014) near the *CTNND2* gene (human chromosome 5 rs10055024) and the *ZFPM2*/*OXR1* genes (human chromosome 8 rs1354342). For reasons that are unclear, GWAS of liver iron and some other traits exhibited many significant lone SNPs, which are less likely to indicate a bonafide SNP association. Raising the minor allele frequency (MAF) cutoff from 5% to 20% removed some but not all of these SNPs (Supplementary File 26).

We also separately performed GWAS for liver iron (Supplementary File 27) and TG (Supplementary File 28) on subsets of the 114 HMDP strains (Supplementary File 1): the AXB/BXA strains plus their A/J and C57BL/6J founders (n=22 strains), RI strains plus their classical inbred A/J, BALBc/ByJ, C57BL/6J, DBA/2J, and C3H/HeJ founders (n=88 strains), and the BXD strains with their C57BL/6J and DBA/2J founders (n=58 strains). Subset GWAS can reveal loci that contribute to the variation in a subset of strains that may be diluted in the full HMDP GWAS by genetic background and effect size differences between populations. The AXB/BXA cohort had two suggestive loci for liver iron on chromosomes 9 (peak SNP rs36851549 at 105,358,144 bp) and 13 (peak SNP rs29530763 at 92,518,336 bp). A potential candidate gene in the chromosome 13 locus, *Zfyve16,* harbors the peak SNP and has been previously linked to iron. It encodes Endofin, a BMP-SMAD regulator of the iron-regulatory hormone hepcidin (Goh, Wallace et al. 2015). The RI and BXD subset liver iron mapping results were similar to each other and both retained a suggestive locus on chromosome 11 that overlapped with the significant chromosome 11 locus in the full HMDP mapping of liver iron. Notably, the chromosome 7 locus at 4.3-4.9 Mb observed in the liver iron full HMDP mapping was not present in any of the strain subset GWAS analyses for liver iron, and the inclusion of the classical inbred strains was needed for this locus to show an association with liver iron. For liver TG, no loci reaching significant or suggestive *P* values were observed for the AXB/BXA subset (Supplementary File 28). However, the chromosome 7 locus at 4.3-5.1 Mb that was significant in the full liver TG HMDP remained as suggestive in the BXD and RI strain mapping subsets. The locus in the BXD subset mapping was much broader than in the RI subset (which includes the BXD subset strains) and the full HMDP GWAS though, spanning ∼21 Mb, indicating that the additional strains add mapping resolution.

#### Anemia and liver copper QTL

We identified loci associated with RBC, HCT, and HGB that overlapped with loci for liver copper and other correlated traits. RBC, HCT, and HGB shared significant loci on chromosome 8 (peak SNP rs50858748 at 83,196,658 bp) and chromosome 9, and also shared many suggestive loci (Supplementary File 20, web resource). Notably, the chromosome 8 and 9 loci are not associated with RBC, HCT or HGB in other HMDP studies where these traits were measured (including studies with mice on chow, high fat/high sucrose, or high cholesterol diets, see the resource website), while some other loci for other red cell traits did overlap across studies, suggesting that the chromosome 8 and 9 associations are influenced by the high iron diet. The chromosome 8 locus is a suggestive locus for liver copper (Supplementary File 22, web resource). The locus on chromosome 9 is broad (∼30-60 Mb) and contains at least two peak regions (peak SNP rs29687983 at 34,844,320 bp and peak SNP rs29689367 at 52,592,601 bp). It is shared with the most significant locus in the GWAS for the percent of body weight attributed to fluid (Supplementary File 17, web resource), and it overlaps on the proximal end with loci for copper, plasma HDL cholesterol, and plasma total cholesterol (Supplementary File 16), traits that are well correlated with these red cell traits and in the same PCA loadings plot quadrant (Figure 4). A locus for brain hippocampal copper in the 31-51 Mb range identified in a study with 28 BXD strains by Jones et al. also overlaps with the chromosome 9 liver copper locus, suggesting that variation in this region may influence copper levels in multiple tissues (Jones, Beard et al. 2008). There are also many other significant and suggestive loci for copper (Supplementary File 22, web resource), although similarly to the GWAS for liver iron, the GWAS for liver copper has many lone SNPs that complicate interpretation.

### Genetics of hepatic gene expression at QTL for liver iron, TG, and copper

We next performed eQTL analyses to identify genes at trait QTL of interest that may be more likely than others to influence the trait. To do this, we ran a GWAS on the mRNA expression of each gene obtained from RNA-Seq data from 114 of the iron loaded livers (one mouse per strain). We identified QTL that are associated with the expression of each gene (eQTL, available on the web resource). The eQTL were classified as local (within 1 Mb of the gene) and likely *cis*-regulated (*cis*-eQTL), or distant (farther than 1 Mb of the gene, i.e. *trans*-regulated (*trans*-eQTL). Genes with a *cis*-eQTL at a trait QTL locus and whose expression correlates with the trait are potential good candidates that may influence the trait. Top candidate genes located at a trait locus of interest, however, are those whose local regulation of expression (i.e. variation in expression that can be explained by SNP variation near the gene) is associated with the trait. We thus examined genes to find those whose *cis* component of expression variance correlates significantly (bicor *P* < 0.05) with each trait, which we refer to here as transcriptome wide association study genes (“TWAS genes”, Supplementary File 29). These genes are strong causal candidates because a *cis-*acting genetic variant associated with gene expression is unlikely to be reactive (that is, downstream of the trait in regulatory hierarchy).

Supplementary File 30 lists genes located within 1 Mb of the overlapping chromosome 7 liver iron and TG GWAS locus, with significant *cis*-eQTL (*P* < 1e-4) noted. A total of 146 genes are present in the 2.5 Mb region (3,373,428 bp to 5,898,707 bp). In the iron loaded livers, there was no detectable expression of 50 of these genes (31 of these were miRNA, lncRNA, snRNA, and TEC Ensembl biotypes that we would not expect to pick up with our library prep). An additional 37 genes had low expression levels that did not meet our cutoffs. Of the 59 remaining genes, 15 had significant *cis*-eQTL, and the *cis* component of the variance in expression of 3 genes (*Isoc2b*, *Gm15922=Pira1*, and *Rdh13*) correlated with liver TG (none with liver iron) and were designated as TWAS genes for liver TG. These 3 genes and the peak *cis*-eQTL SNPs for these genes were all located within the chromosome 7 locus, and the peak *cis*-eQTL SNPs all were suggestive or significant GWAS SNPs for liver iron and TG. At the chromosome 7 13-15 Mb locus for liver iron, the only TWAS gene is *Selenow*. Supplementary File 31 lists genes that are located within 1 Mb of the distal chromosome 11 GWAS peak for liver iron (117,974,693-121,568,141 bp). TWAS genes at this locus in order of increasing TWAS *P* value are *Alyref*, *Gaa*, *Slc26a11*, *Usp36*, and *Pcyt2*. The chromosome 6 locus and the chromosome 7 locus near *Luzp2* for liver iron have no TWAS candidates for liver iron.

For liver copper, there are several TWAS genes at the chromosome 9 locus (Supplementary File 29). The most significant are *Sc5d, Pus3, Sorl1,and Slc37a2*. Another copper locus of interest is on chromosome 19 spanning 3.3 to 7 Mb with peak SNP rs13467290 at 6,542,381 Mb. It may encompass two loci based on its LD structure. Antioxidant enzyme *Prdx5* is the top TWAS gene for liver copper at the peak SNP. Two other genes at this locus are also TWAS genes for copper: *Ccs* (the copper chaperone for SOD1) and *Ap5bl*. Details of expressed genes, eQTLs, and their correlations with traits at significant and suggestive GWAS loci can be explored further on the resource webpage. We note that GWAS associations may result from gene coding variants that would not necessarily be reflected in expression variation. *Trans*-eQTL associated with the overlapping chromosome 7 locus for liver iron and TG are described in Supplementary File 24.

### Liver eQTL hotspots

Square plots were used to visualize liver eQTL across the genome. The position of each SNP associated with the expression of a gene was plotted versus the position of the gene, for eQTL in this study (Figure 6A, left bottom), and for comparison, for eQTL from RNASeq data from a different study where control vehicle-treated male mice were fed a chow diet (Figure 6A, right bottom) (Tuominen, Fuqua et al. 2021). *Cis*-regulated genes are visible as dots on the diagonal, while *trans* regulated genes are dots off the diagonal. Many of the most significant eQTL, visible in the square plots as larger colored circles per the legend, are conserved between the studies. SNPs that may influence the expression of many genes in *trans*, or eQTL regulatory hotspots, are apparent as linear veritical patterns of eQTL (Civelek and Lusis 2014). The tracks above the square plots quantify the linear vertical patterns and plot the number of genes associated with a SNP in *trans* (shown here for all genes with an eQTL at this SNP with *P* ≤ 1e-6 that are not on the same chromosome as the SNP). Overall, there is very little overlap in the top hotspot SNPs between these two studies (Figure 6A and Figure 6-Source Data 1). This likely reflects the effect of the high iron diet on transcriptional networks and cell composition but could also be influenced by the age of the mice (10-11 weeks vs 16-18 weeks) and study strain composition. The top 118 hotspot SNPs (in terms of number of *trans* associated genes) in the iron study, ranging from 30 to 169 *trans* associated genes, each associate with fewer than 5 genes in *trans* in the chow study. The top 10 SNPs in the chow study, each associated with 66 to 104 genes, are associated with 2 or fewer genes in the iron study.

**Figure 6.**
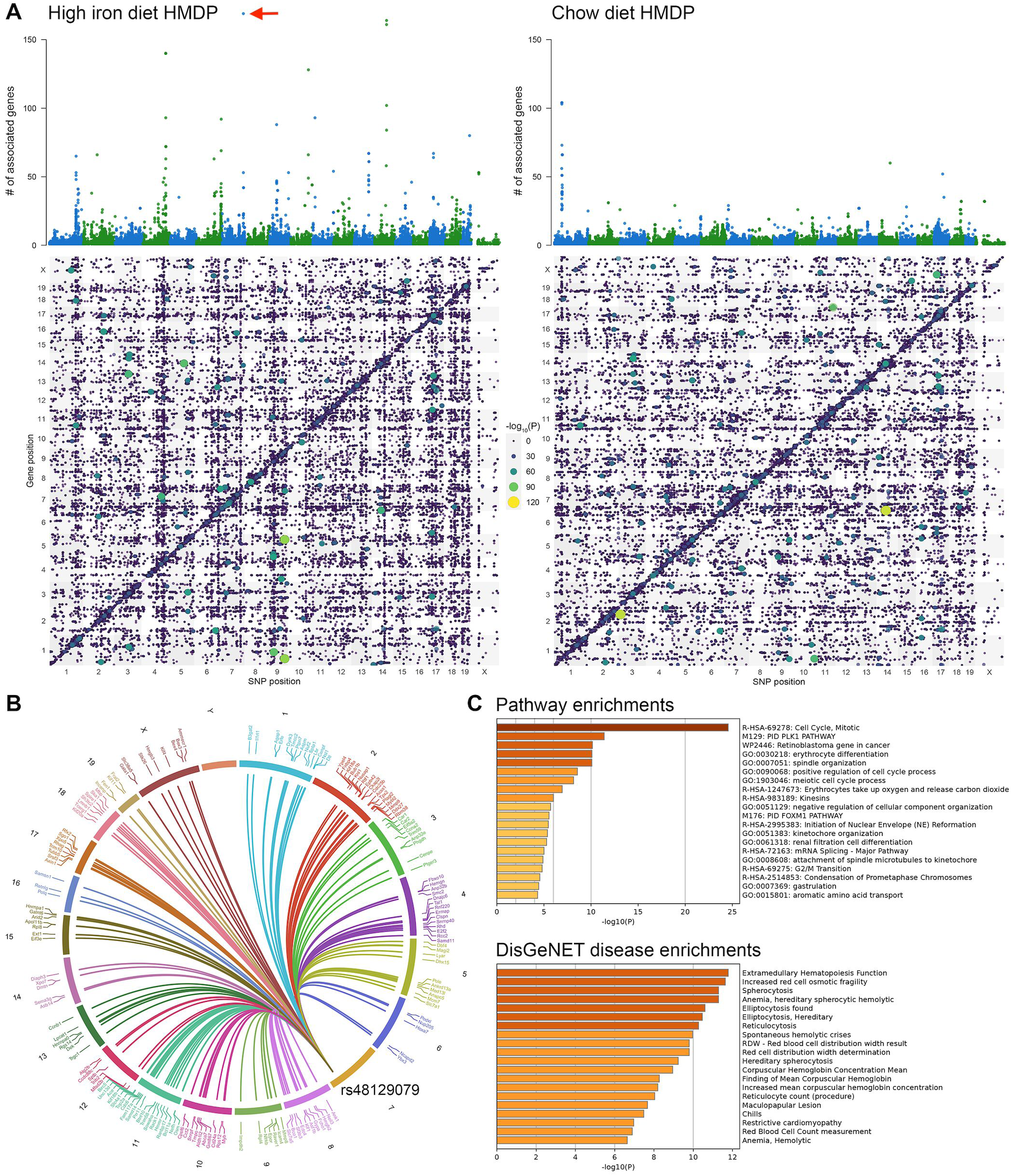
Regulatory liver eQTL hotspots. **(*A*)** In the top panels, each dot represents the number of genes with a significant liver *trans*-eQTL at a given SNP in the genome in the high iron diet HMDP (left panel) and a reference study where mice were fed a chow diet (right panel) (Tuominen, Fuqua et al. 2021). A significant *trans*-eQTL for a gene is defined here as an eQTL for that gene with *P* <1e-6 that is not located on the same chromosome as the gene. The top panels are aligned with square plots below for each study. The square plot on the right was generated with eQTL data from the chow control group in the reference study (Tuominen, Fuqua et al. 2021) and corresponds with Figure 12A in that study, although the appearance differs due to the different eQTL cutoffs used to generate each plot. In the square plots, gene *cis*-eQTL (located within 1 Mb of the gene) with *P* < 9.05e-4 for the high iron study and *P* < 2.1e-3 for the chow study, and gene *trans*-eQTL with *P* < 1e-6, are plotted by gene position (y-axis) versus SNP position in the genome (x-axis). The -log_10_*P* values of the eQTLs are sized and colored according to the legend at center. *Cis*-eQTL are on the diagonal. The visual appearance of vertical lines of dots represent eQTL regulatory hotspots. The top eQTL hotspot SNP in the high iron HMDP study, rs48129079, is located on chromosome 7 and is associated significantly (*P* < 1e-6) in *trans* with the liver mRNA expression of 169 genes (red arrow). **(*B*)** Circos plot of the mouse chromosomes showing the location and names of the 169 genes significantly associated in *trans* with the top eQTL hotspot rs48129079. **(*C*)** Metascape pathway enrichments (top) and DisGeNET (https://www.disgenet.org) disease enrichments for the 169 genes associated with the top hotspot rs48129079. Source data for panel A is in Figure 6-Source Data 1.

To investigate the types of genes regulated by top hotspots in the iron loaded livers, we performed pathway analysis of the genes associated in *trans* with each of the top 15 iron study hotspots (encompassing the top 71 SNPs, since several SNPs in LD often tag one hotspot). Several of these hotspots are linked to the extramedullary hepatic erythropoiesis that was observed histologically and are enriched in genes expressed in CD71+ (transferrin receptor high) erythroblast cells (Supplementary File 32). Polo-like kinase 1 (*Plk1*) is located ∼1 Mb from the top hotspot SNP rs48129079 and is a potential driver of this hotspot (Figure 6B). *Plk1* is the only gene with a significant liver *cis*-eQTL (*P* = 2.295e-6) at this SNP. By pathway enrichment analysis, the 169 genes associated in *trans* with this hotspot are enriched for genes in the PLK1 pathway (Figure 6C). PLK1 is noted to be involved in cell cycle and erythrocyte development (enucleation) (Huang, Hale et al. 2018) and is also upregulated by key erythropoietic genes GATA1 (Shen, Li et al. 2012) and HIF2A (Dufies, Verbiest et al. 2021). DisGeNET disease enrichments for the 169 genes for this hotspot predominantly include red cell related conditions (Figure 6C). In addition, at least two of the top 15 hotspots are associated with lupus erythematosus and immune function, which may be related to extramedullary erythropoiesis via the immunosuppressive or immunomodulatory properties of erythropoietin and erythroid precursor expansion (Elahi and Mashhouri 2020; Eswarappa, Cantarelli et al. 2021). *Rcbtb2*, the only gene with a significant *cis*-eQTL (*P* = 3.580e-5) for the top hotspot (peak SNP rs45680344) of those with enrichments related to lupus, has itself been previously linked to lupus (Santer, Wiedeman et al. 2012; Sandling, Pucholt et al. 2021). While two of the top 15 hotspots, hotspot #7 and hotspot #15 (Supplementary File 32), are located at GWAS loci significant for liver copper and liver iron, respectively, the hotspot SNPs are not among those significant in the metal GWAS. Hotspots and associated genes are listed in Supplementary File 33 and can be visualized on the resource website.

### Top genes and pathways associated with liver iron and TG

We next looked at the correlation between hepatic gene expression and traits across all 114 HMDP strains. The top five genes with hepatic expression positively correlated with liver iron (by *P* value) are *Rbm3* (induced by hypothermia and low oxygen), *Mt1* and *Mt2* (involved in metal homeostasis and the oxidative stress response), *Arg1* (urea cycle), and *Fkbp5* (induced by stress), and the top five negatively correlated genes are *Stk16* (a kinase), *Slc16a2* (thyroid hormone transport), *Cyp2j5* (arachidonic acid metabolism), *Pgpep1* (a pyroglutamyl peptidase), and *Nrp1* (a cell surface receptor; also impacts mitochondrial iron metabolism via ABCB8) (Issitt, Bosseboeuf et al. 2019). We performed pathway enrichment analysis using Metascape to identify biological pathways associated with the top 500 genes positively and negatively correlated with liver iron and TG by *P* value (Figures 7, 8, and Supplementary Files 34-36). Top pathways associated with genes positively correlated with liver iron included those related to ribosome biogenesis and translation, response to oxygen levels including the HIF1a and VEGF pathways, positive regulation of cell death and the intrinsic apoptosis pathway, RNA polymerase II transcription termination and regulation of RNA splicing, and the TNF-alpha/NF-κB signaling complex 6 (Figure 7A). Top diseases sharing these genes included those related to low oxygen levels, including anemias, myocardial ischemia, and pulmonary hypertension (Figure 7B). There was an enrichment for liver-specific genes, and top proteins that regulate these genes are PSMB5 (proteosome component), TAF9B (involved in transcription initiation), FOXE1 (hypothyroid transcription factor), HIF1A (hypoxia transcription factor), and NPAS2 and ARNTL (circadian rhythm transcription factors) (Figure 7C-E).

**Figure 7.**
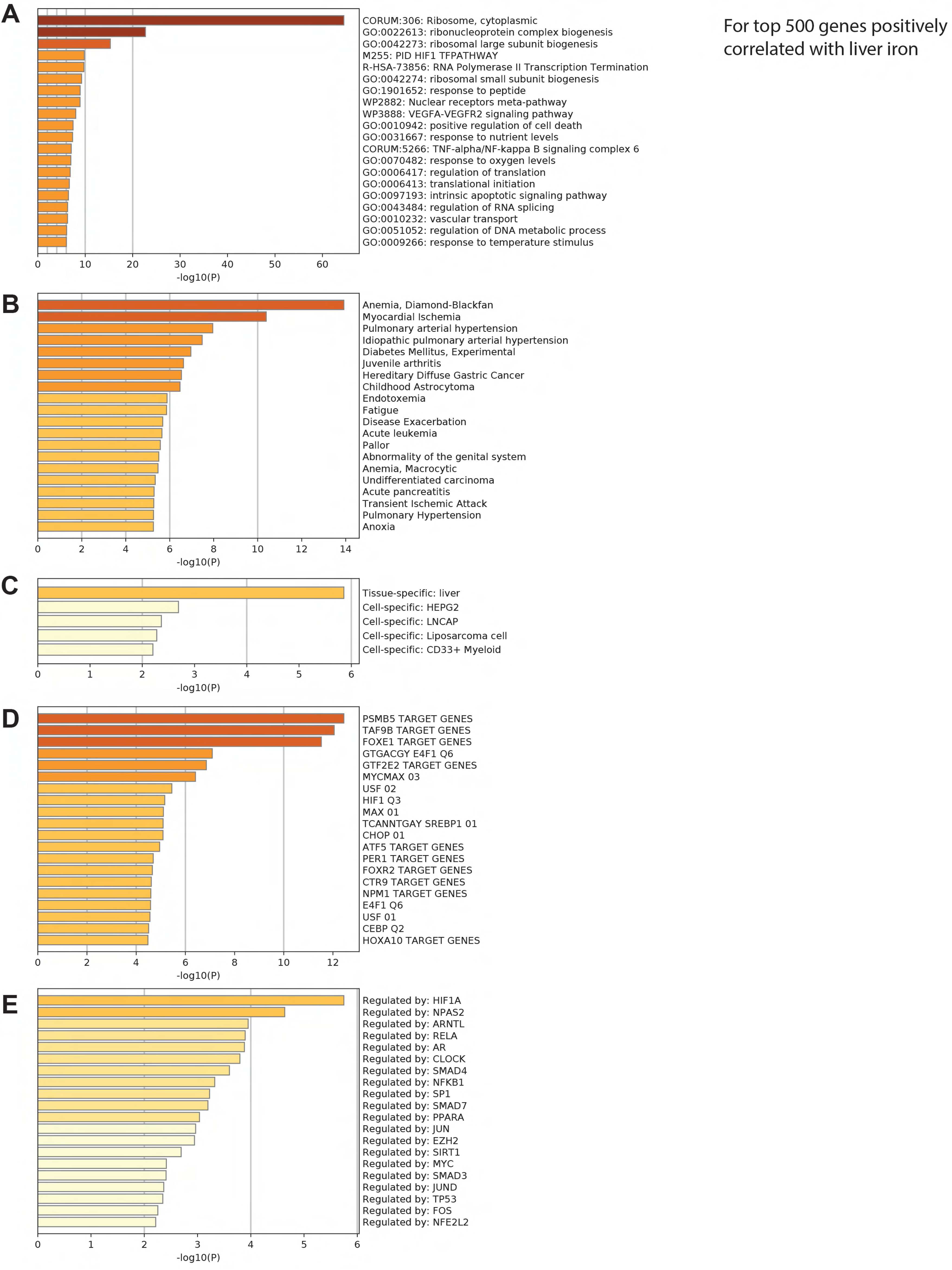
Top Metascape enrichment analysis annotations, ranked by their -log_10_*P* values, for the top ∼500 genes by *P* value <u>positively</u> and significantly correlated (bicor *P* < 0.05) with liver iron across all 114 strains of mice. (***A***) Metascape pathway enrichments. (***B***) Metascape disease enrichments from DisGeNET (https://www.disgenet.org). (***C***) Metascape cell type enrichments. (***D***) Metascape transcription factor target enrichments (***E***) Metascape transcriptional regulation enrichments from TRRUST (http://www.grnpedia.org/trrust). Gene expression correlation with liver iron is given in Supplementary File 36.

**Figure 8.**
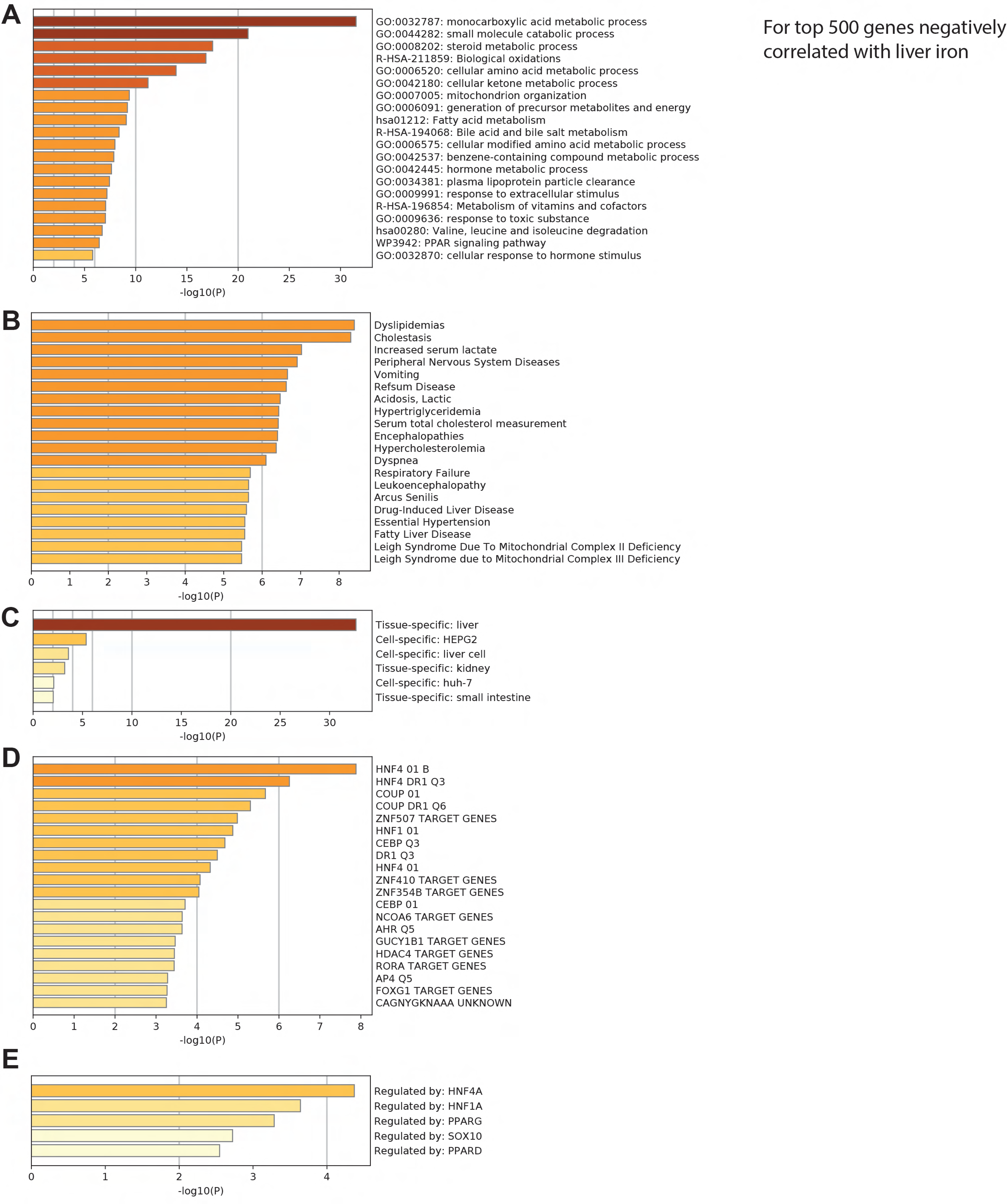
Top Metascape enrichment analysis annotations, ranked by their -log_10_*P* values, for the top ∼500 genes by *P* value <u>negatively</u> and significantly correlated (bicor *P* < 0.05) with liver iron across all 114 strains of mice. (***A***) Metascape pathway enrichments. (***B***) Metascape disease enrichments from DisGeNET (https://www.disgenet.org). (***C***) Metascape cell type enrichments. (***D***) Metascape transcription factor target enrichments (***E***) Metascape transcriptional regulation enrichments from TRRUST (http://www.grnpedia.org/trrust). Gene expression correlation with liver iron is given in Supplementary File 36.

Top pathways for genes negatively correlated with liver iron are broadly related to liver metabolic processes including the metabolism of small molecules, ketones, lipids, amino acids, fatty acids, bile acids, steroids, vitamins, and cofactors. Mitochondrion organization, the response to toxic substances, and the PPAR signaling pathway were also negatively correlated with iron levels (Figure 8A). Specifically, the mRNA expression of most genes involved in the KEGG fatty acid degradation and elongation pathways, as well as the valine, leucine, isoleucine, and tryptophan degradation pathways, were negatively correlated with liver iron across the HMDP. Diseases associated with these genes included those related to lipid dyshomeostasis, mitochondrial disease, liver damage, and lactic acidosis (Figure 8B). The top enriched regulators of this geneset negatively correlated with liver iron included the transcription factors HNF4A and HNF1A (hepatic gluconeogenesis, lipid metabolism, and liver tissue maintenance) (Sahoo, Singh et al. 2020), COUP (metabolic gene regulation) (Ashraf, Sanchez et al. 2019), and PPARG (lipid metabolism and steatosis regulator) (Lee, Park et al. 2018) (Figure 8D-E).

We also looked at hepatic gene expression correlations with liver TG. The top five genes positively correlated with liver TG across all strains by *P* value are *Raet1e* (immune response), *Serinc2* (phosphatidylserine metabolism), *Ngfr* (metabolic gene oscillation), *Zfp872*, and *Angptl6*, and the top five genes negatively correlated are *Klhl3* (facilitates ubiquitination), *Mat1a* (one carbon metabolism), *Safb* (transcriptosome complex assembly), *Fmo3* (xenobiotic metabolism), and *Amy1* (hepatic glycogen metabolism). Top pathways associated with liver TG were related to lipid biogenesis as well as to injury and repair, including response to drugs, Phase I and II metabolism, wounding, neutrophil degranulation, smooth muscle cell migration, and supramolecular fiber organization (Supplementary File 34). Sulfur compound metabolic processes were also enriched, which can be involved in managing oxidative stress (Miller and Schmidt 2020). Top diseases associated with these genes included those related to dyslipidemias, drug abuse, and toxicity, and top regulators of these genes included HNF1 (liver tissue maintenance), ZNF410, RELA and NFKB1 (NF-κB complex), and ESR1 (Supplementary File 34).

Top pathways negatively associated with liver TG were related to amino acid metabolism including the urea cycle (Supplementary File 35). Other top pathways included rhythmic and circadian activity, response to xenobiotics, RNA splicing and ribonucleoprotein complex biogenesis, and the breakdown of sulfur-containing cysteine and methionine. Top diseases associated with these genes were urea cycle and amino acid disorders, hematological conditions and cancers, and gallbladder cancers. Most of the genes were liver specific, but there was also an enrichment in early erythroid genes. Enriched top regulators of negatively correlated TG genes included SRY, HNF3, HES4, and CEBPB. All gene by gene and gene by clinical trait correlations can be further explored on the resource webpage, and gene sets of interest can be downloaded for pathway analysis.

### Differential gene expression in response to the high iron diet

To better understand what gene expression was reactive to the high iron diet, we looked at genes that were differentially expressed in the livers of the C57BL/6J mice in the Control study fed the 50 ppm iron sufficient control diet vs the 20k ppm high iron diet (Supplementary File 37 and Supplementary File 38). Top genes downregulated by the 20k ppm high iron diet (within the top 50 most differentially expressed genes by *P* value) included those in the Mup family, which are small lipocalin proteins regulated by testosterone, thyroxine, and growth hormone (Thoß, Luzynski et al. 2015). Also downregulated were several cytochrome P450s (*Cyp4a12a*, *Cyp2d9*, *Cyp2e1*) and lipid metabolism genes (*Acaa1b*, *Acat3*, *Apoa1*). Top upregulated genes included genes previously linked to iron overload (*Ftl1, Ftl1-ps1*, *Atoh8*, *Slc40a1*, *Id4*, *Bmp6*) (Kautz, Meynard et al. 2008; An, Wang et al. 2018), interferon-induced genes (*Oasl1*, *Ly6a*, *Rsad2*, *Oasl2*, *Siglec1*)(Jeidane, Scott-Boyer et al. 2016), hypoxia regulated genes (*Rgs4*, *Rgs5*, and *Pkm*) (Williams, Khadka et al. 2018), and Kupffer and infiltrating monocyte specific gene *Clec4f* (Jiang, Tang et al. 2021).

The top enriched KEGG and Reactome pathways from pathway enrichment analysis of the DE genes (*P*_adj_ < 0.05) with decreased expression in response to the 20k ppm high iron diet (1,062 mouse genes, Supplementary File 38) included those related to lipid metabolism (e.g. fatty acid degradation, bile acid and bile salt synthesis, beta-oxidation, steroid metabolism, arachidonic acid metabolism), amino acid metabolism (e.g. valine, leucine, and isoleucine degradation, tryptophan metabolism, and lysine degradation), biological oxidations (e.g. phase I functionalization of compounds, cytochrome P450s), the peroxisome (e.g. peroxisomal lipid metabolism and protein import), and the TCA cycle and respiratory electron transport. Top associated tissues and cells from PaGenBase were liver and HepG2, and top associated diseases from DisGeNet were metabolic acidosis, hyperammonemia, hypoglycemia, and dyslipidemias.

The top enriched KEGG and Reactome pathways from pathway enrichment analysis of the DE genes (*P*_adj_ < 0.05) with increased expression (1,069 mouse genes, Supplementary File 38) were immune related pathways (e.g. neutrophil degranulation, cytokine signaling, adaptive immune response), hemostasis (activation, signaling, aggregation, and degranulation of platelets), and interferon signaling. Top associated tissues from PaGenBase were blood, spleen, and bone marrow, and top associated diseases from DisGeNet included lupus, bacterial infections, and myocardial ischemia.

We looked in more detail at gene expression in several pathways related to the dysregulated liver and plasma lipids induced by the high iron diet. Almost all of the genes in the KEGG fatty acid degradation pathway were downregulated, while there was a slight (2.4 fold) increase in the expression of *de novo* lipogenesis enzymes *Fasn* and *Acaca*, although only the latter gene reached adjusted significance. Many of the genes in primary bile acid biosynthesis were down, including the rate limiting enzyme, *Cyp7a1* (Zhang, Huang et al. 2009). Expression of *Abcg5* and *Abcg8*, important in biliary cholesterol secretion, were decreased (Zein, Kaur et al. 2019), while *Npc2*, involved in cholesterol clearance from lysosomes, was increased (Meng, Heybrock et al. 2020). The expression of *Apob*, a component of LDL and VLDL, was down, in agreement with the decrease in plasma LDL and VLDL cholesterol levels. *Apoa1* and *Apoa2*, components of HDL, were also decreased, as was *Lipg*, which encodes endothelial lipase (EL). EL hydrolyzes phopholipids and TG from TG rich lipoproteins (Khetarpal, Vitali et al. 2021). Decreased EL expression is associated with increased plasma TG and HDL cholesterol, only the former of which was observed here. The expression of *Angptl3*, an inhibitor of lipoprotein lipase and EL, was increased, which is in accordance with the increase in plasma TG observed (Sylvers-Davie and Davies 2021). In the KEGG steroid biosynthesis pathway, only two genes, *Soat1* and *Tm7sf2*, were differentially expressed (*P*_adj_ < 0.05). *Soat1* expression increased 2.5 fold, while *Tm7sf2* expression decreased 1.8 fold, in response to the high iron diet. *Soat1* encodes the enzyme acyl-coenzyme A: cholesterol acyltransferase 1 (ACAT1) that is involved in making cholesterol esters, which are used to make lipid droplets to store TG. The increase in *Soat1* expression is in agreement with the observed increase in liver EC in the mice on the high iron diet, as well as the trend toward increased liver TG and the enlarged lipid droplets as visualized by Oil Red O. The protein encoded by *Tm7sf2* has been shown to increase liver LXR signaling and influence NF-κB activation, with knockout in mice leading to NF-κB activation and TNFα up-regulation (Bellezza, Roberti et al. 2013).

To gain insight into the top differentially expressed genes in response to the 20k ppm high iron diet in C57BL/6J that also had expression highly correlated with liver iron across all 114 HMDP strains, we ranked genes by increasing differential expression *P* value and compared these with genes ranked by correlation with liver iron concentration across the whole HMDP. The 96 genes with sum of ranks less than 1000 (Supplementary File 37) were mostly (80/96) downregulated in response to iron in the Control study and negatively correlated with liver iron in the HMDP (83/96), and they were correspondingly enriched in genes involved in lipid metabolism, peroxisomal protein import, biological oxidations, disorders of bile acid synthesis and biliary transport, and response to toxic substances.

### Network modeling of clinical traits

#### Weighted gene coexpression network analysis (WGCNA)

We next used WGCNA, an unbiased analysis that aggregates highly co-expressed genes into gene modules, to model mRNA co-expression networks in the liver and to then use these to understand the association of gene networks with clinical traits across the 114 mouse strains of the HMDP. We identified 13 co-expression modules (ranging in size from 54 to 2,927 genes) excluding the grey (unassigned genes that were not co-expressed) module (Figure 9A, Figure 9-Source Data 1), and using enrichment analysis of the member genes, identified pathways associated with each module. Several module eigengenes showed strong correlation with traits. Seven modules were significantly (*P* < 0.05) correlated with liver iron, four with liver TG, and nine with liver copper and red blood cell count (Figure 9B). Liver iron was most strongly positively correlated with the magenta module (bicor = 0.55), which is associated with FOXA2, lipid metabolism, FMOs, oxygen level response, and ferroptosis, and most negatively correlated with the blue module (bicor = -0.50), which is enriched in terms related to TCA cycle, ETC, mitochondria, lipid metabolism, and detoxification.

**Figure 9.**
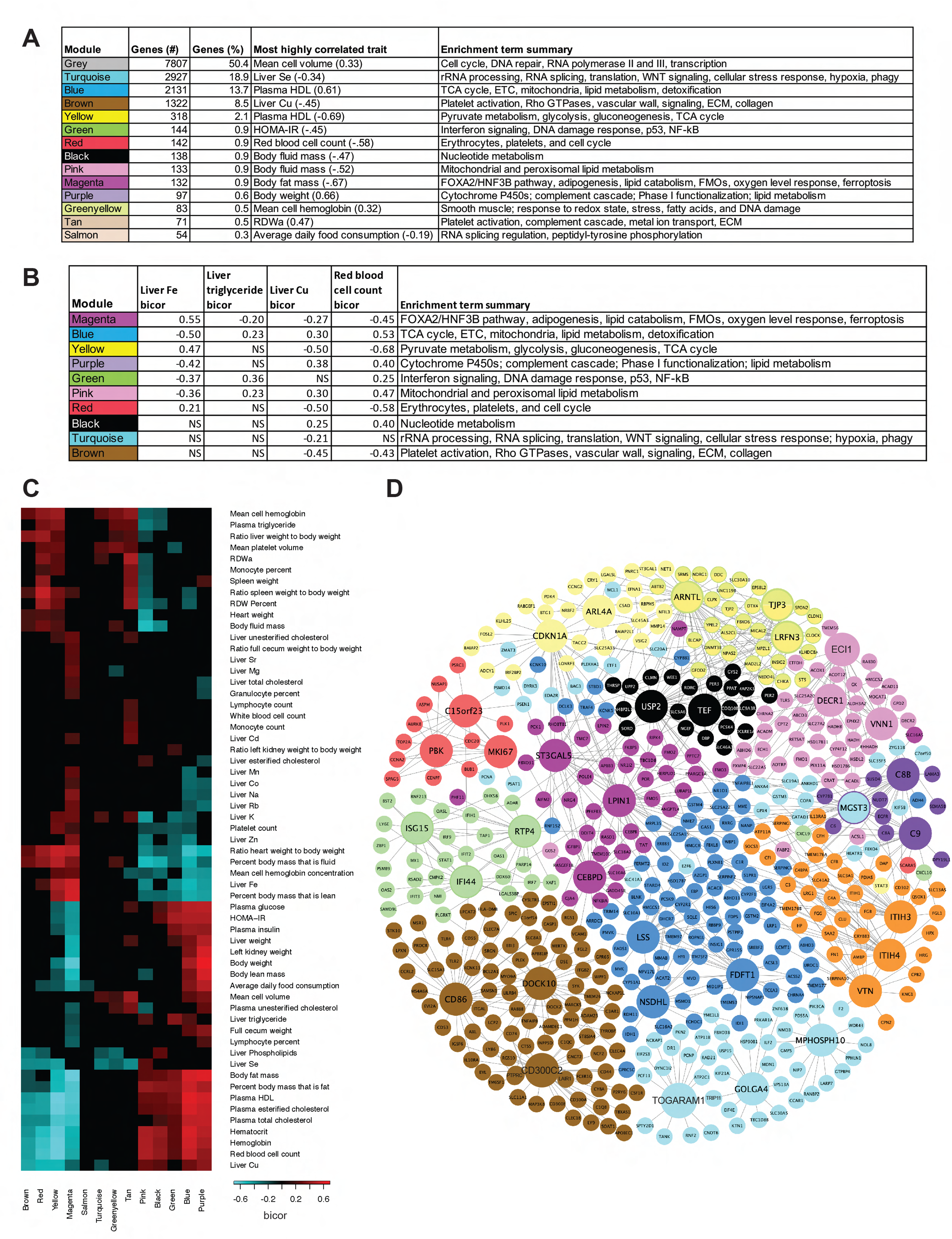
WGCNA co-expression modules, trait correlations, and key drivers. **(*A*)** The 13 modules (excluding the grey unassigned genes module) identified by WGCNA in the liver gene expression data across the HMDP, with the number of genes in each module, the percent of total genes in each module, the most highly correlated trait with the module’s eigenvector by *P* value (bicor shown in parentheses), and top pathway enrichment term summaries for the module gene set based on Reactome and Metascape analyses. **(*B*)** Modules significantly correlated (bicor *P* < 0.05) with liver iron, liver triglycerides, liver copper, or red blood cell count, ordered by correlation with liver iron, with the trait bicor and the module gene set summary enrichment terms. **(*C*)** Hierarchical clustered heatmap of the correlations between the WGCNA modules and 59 traits with at least one significant module correlation. Only body composition traits from the final NMR just before tissue collection were included in the analysis. Significant (bicor *P* < 0.05) trait to module correlations are shown in blue (negative correlations) and red (positive correlations), while correlations that were not significant are in solid black. Patterns in the heatmap highlight traits and modules that are similarly correlated. **(*D*)** Visualization of WGCNA module key drivers and top connected genes as identified using Mergeomics KDA analysis. The largest nodes are the key drivers. Genes are colored by their WGCNA module membership, and for key drivers, their border color denotes the module for which they are a key driver. There are some instances where genes are key drivers for modules that they themselves are not in, e.g. *MGST3* is a key driver for the purple module (and has a purple border to show this) but is itself in the WGCNA turquoise module. As an exception, the greenyellow module members (top right) have green borders but are all members of the greenyellow module. The module that appears orange in the figure represents the tan module. WGCNA data used to make this figure is in Figure 9-Source Data 1.

For broad visualization of how the gene expression modules were related to traits, we grouped clinical traits by hierarchical clustering according to their module bicor associations (Figure 9C). The traits clustered into two main visual blocks: Group 1 (traits above and inclusive of “Percent body mass that is lean” in the plot) including all tissue weights besides left kidney and liver, liver cholesterol traits, liver iron and most other metals, and Group 2 (traits below and exclusive of “Percent body mass that is lean” in the plot) including plasma glucose, insulin, and lipids (except for TG); body weight, liver TG and PL, red cell traits, and liver copper and selenium. Several traits exhibited similar overall module correlations. Most Group 1 traits were either negatively or not correlated with the pink, black, green, blue, and purple modules, and either positively or not correlated with the magenta, yellow, red, and brown modules. Group 1 traits including MCH, MPV, RDWa and RDW percent, plasma TG, spleen weight, heart weight, absolute fluid mass, and the ratio of liver weight to body weight showed overall positive correlations with the yellow and red modules, the two modules most correlated (negatively) with red blood cell count. Conversely, Group 2 traits showed overall positive or no correlation with the pink, black, green, blue, and purple modules, and overall negative or no correlations with the magenta, yellow, red, and brown modules. HCB, HCT, RBC, liver copper, plasma HDL cholesterol, and EC and TC were overall strongly negatively correlated with red, yellow, and magenta modules.

We looked at the WGCNA module membership of genes that were significantly differentially expressed (DE) (*P*_adj_ ≤ 0.05) between the 50 ppm iron sufficient control diet and 20k ppm high iron diet in the Control C57BL/6J study (Supplementary File 39). Of the 15,083 genes that we had both WGCNA and DE data for, 2115 (14%) were DE and 1641 (78%) of these DE genes were WGCNA co-expression module members (i.e. in modules excluding the grey module which contains all unassigned genes that were not co-expressed). For comparison, of the 12,968 genes that were not DE, 5,976 (46%) were WGCNA co-expression module members, indicating an overall enrichment of DE genes in WGCNA co-expression modules (two-tailed Chi-square test *P* < 0.0001). Over 50% of the genes in the purple, green, and pink modules were DE genes (Supplementary File 40). Of all 2115 DE genes, 948 (45%) were also correlated with liver iron (bicor *P* ≤ 0.05) across the HMDP. Of these DE and liver iron correlated genes, 62.8% were negatively DE and negatively correlated with liver iron, 2.8% were negatively DE and positively correlated with liver iron, 26.7% were positively DE and positively correlated with liver iron, and 7.7% were positively DE and negatively correlated with liver iron (Supplementary File 39). Notably, 96% of the DE genes in the magenta module were correlated with liver iron (mostly positively DE in response to the high iron diet and all positively liver iron correlated across the HMDP), followed by 84% of those in the blue module (mostly negatively DE and all negatively liver iron correlated) and 78% of those in the yellow module (mostly positively DE and all positively liver iron correlated) (Supplementary File 40). There were 73 genes with increased expression on the high iron diet that were negatively correlated with liver iron across the HMDP, of which 42 (58%) were in the green module (associated with interferon signaling and the DNA damage response).

Finally, QTL mapping was performed on the first principal components of the modules (e.g. their eigengenes, Figure 9-Source Data 1) to identify genetic loci that may influence module gene expression. Significant loci (5% false discovery rate (FDR), *P* < 4.1e-6) were identified for five modules (brown, green, pink, yellow, and greenyellow). Examples of two loci for the green module, and the top candidate genes based on *cis*-eQTL at these loci, are discussed in Supplementary File 24.

#### Mergeomics

We next used Mergeomics weighted key driver analysis (KDA) to identify key drivers associated with the WGCNA modules (Shu, Zhao et al. 2016; Ding, Blencowe et al. 2021). To identify key drivers, genes belonging to each WGCNA module were mapped onto a Bayesian regulatory network whose connections (or edges) were generated using directional data from multiple studies (see Methods). Key drivers for a given WGCNA module are those genes in the Bayesian regulatory network that are most significantly enriched in network neighbors from that WGCNA module. Key drivers are represented in Figure 9D by the largest nodes, with top connected genes from the network shown. Key drivers of a WGCNA module are top genes predicted to, if perturbed in mechanistic studies, influence the expression of genes in that module.

We looked at the top key drivers for WGCNA modules whose eigengenes were most correlated with liver iron and TG. Of note, the modules could be related to these traits in a causal and/or reactive manner. The top key drivers for the module most highly correlated with liver iron across the HMDP, the magenta module, are *St3gal5, Cebpd*, and *Lpin1*, of which only *St3gal5* was significantly differentially expressed at the mRNA level in the C57BL/6J Control study in response to the high iron diet. *St3gal5* converts lactosylceramide into the sialyl-lactosylceramide (GM3), a precursor to many other gangliosides. GM3 has been shown to impact EGF, bFGF, TNFα, PDGF, VEGF, and insulin signaling, potentially via its effects on glycolipid enriched membrane microdomains (Boccuto, Aoki et al. 2014), but a direct link to iron is not clear. *Cebpd* encodes the immune and inflammation regulator transcription factor C/EBP-δ. C/EBP-δ has been shown to activate the transcription of the systemic iron regulator hepcidin in response to the pro-inflammatory cytokine IL-1β (Kanamori, Murakami et al. 2019). *Cebpd* has also been shown to be regulated by iron via SMAD3 (Li, Song et al. 2015). *Lpin1* is involved in many aspects of lipid metabolism and inflammation (You, Jogasuria et al. 2017; Zhang and Reue 2017). Notably, induction of *Lpin1* via Hif1a by hypoxia caused TG and lipid droplet accumulation (Mylonis, Sembongi et al. 2012), and adipose-specific overexpression of *Lpin1* in a mouse model of alcoholic steatosis led to increased hepatic ferroptosis, steatosis, and liver damage (Zhou, Ye et al. 2019).

The top key drivers for the module most associated with liver TG, the green module, are *Isg15*, *Ifi44*, and *Rtp4*, all genes whose mRNA expression was significantly increased in the Control study in response to the high iron diet and that are known to influence and be influenced by the interferon response (Perng and Lenschow 2018; DeDiego, Martinez-Sobrido et al. 2019; He, Ashbrook et al. 2020). Interferon is induced as part of the DNA damage response via NF-κB (Brzostek-Racine, Gordon et al. 2011) and has been linked to hepatic steatosis and fibrosis in humans and mice (Hart, Fabre et al. 2017; Møhlenberg, Terczynska-Dyla et al. 2019).

We next used Mergeomics to identify additional candidate genes that may influence liver iron levels and associated pathology, but this time by a directional approach using GWAS loci to identify modules associated with each trait. Trait-associated gene sets were generated by mapping trait GWAS data to genes based on the liver eQTLs generated from the RNASeq data in this study. The full summary statistics of the GWAS data were used as input, and Marker Set Enrichment Analysis (MSEA) highlighted pathways and modules enriched in trait-associated genes. To identify key drivers, genes belonging to each of these trait-associated pathways or modules were mapped onto the Bayesian regulatory network. Key drivers for a given trait-associated module or pathway are those genes in the Bayesian regulatory network that are most significantly enriched in network neighbors from that trait-associated module or pathway. Several modules were found to be enriched in genes associated with liver iron GWAS loci, and key drivers for these modules (Figure 10, represented by the largest circles in the network) are involved in 1. lipid metabolism (cholesterol biosynthesis and SREBP regulated: *SQLE*, *DHCR7*, and *FDFT1*; bile acid metabolism: *AKR1D1*; fatty acid biosynthesis: *FASN*); 2. nitrogen metabolism and the urea cycle: *ASL* and *CPS1*; 3. cell cycle (DNA synthesis: *CCNA2*, *CDCA8*, *MCM6*; kinetochore: *C15orf23*; cytokinesis: *RACGAP1*); 4. *NRF2* regulated phase II conjugation of glutathione S-transferases: *GSTM4* and *GSTA1* (Chanas, Jiang et al. 2002); 5. phase I functionalization of compounds: *ACSS2*; and 6. peroxisomal lipid metabolism and protein import: *EHHADH*. Although not key drivers themselves, ABC transporters and genes involved in pyrimidine metabolism were also members of pathways associated with liver iron.

**Figure 10.**
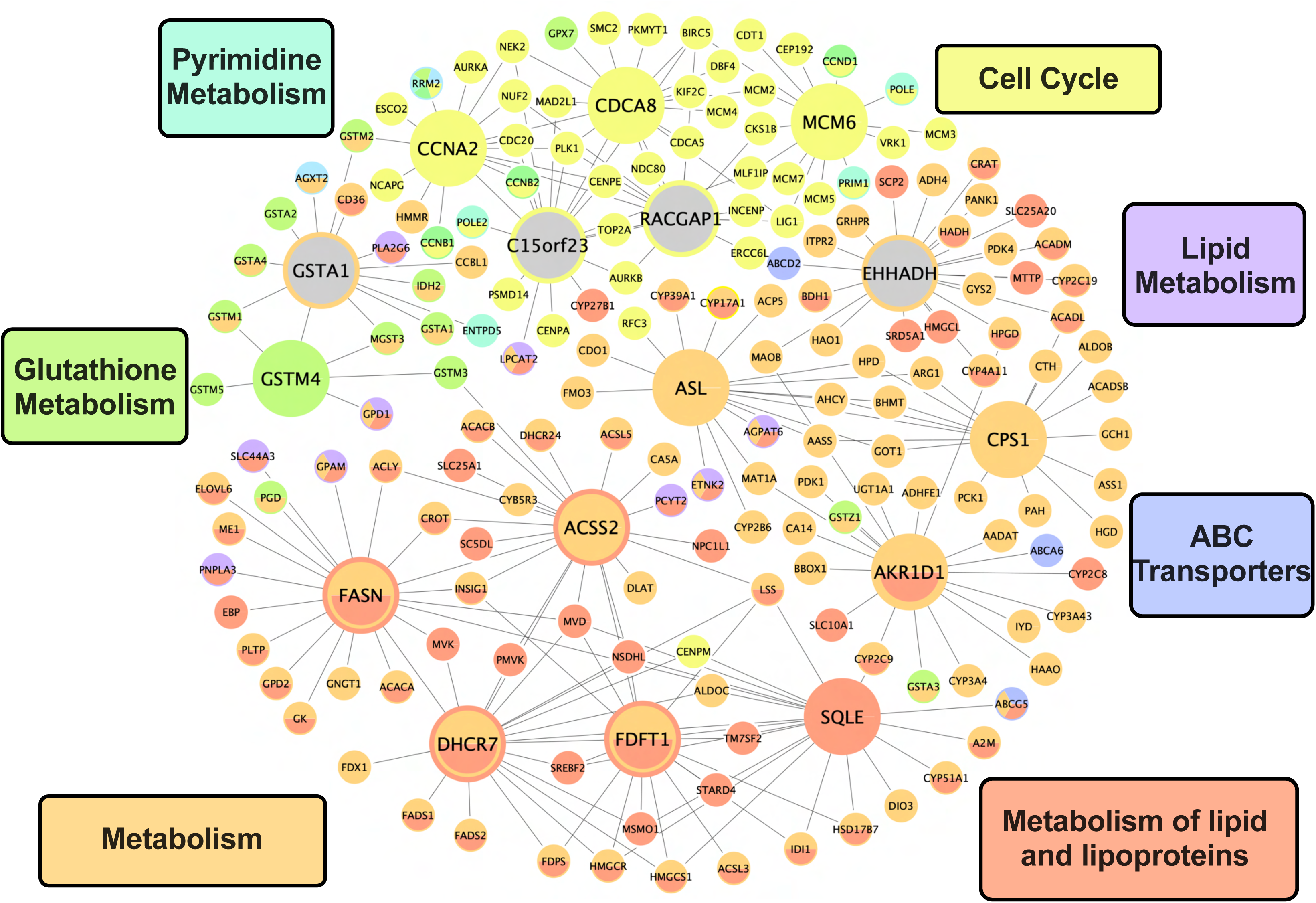
Key driver genes associated with liver iron concentration identified by Mergeomics MSEA and KDA. The key drivers (large nodes) and their top connected genes (small nodes) that belong to iron-associated modules are shown. The interior color(s) of the large nodes indicate the liver iron associated module(s) that the node gene is in, and the rim color(s) of the large nodes indicate the liver iron associated module(s) where that node gene is a key driver. If there is a double rim, the node gene is a key driver of two liver iron associated modules. The large nodes with grey interiors are key drivers but are not themselves module members of any liver iron associated modules. The small nodes are colored by the iron associated modules to which they belong. Some small nodes have the appearance of a colored rim, but this is an artifact that is not meaningful. The sky blue color associated with two small nodes, AGXT2 and RRM2 on the top left, represents a module associated with nucleotide metabolism.

For liver TG, the blue WGCNA coexpression module was found to be enriched in genes associated with liver TG GWAS loci. Key drivers for liver TG (Supplementary File 41) include two that overlap with key drivers for liver iron, *FDFT1* and *SQLE*. The other top key drivers for liver TG are *LSS* (which catalyzes the first step in the biosynthesis of cholesterol, steroid hormones, and vitamin D), *NSDHL* (downstream of LSS in cholesterol synthesis), and *QDPR* (involved in the recycling of tetrahydrobiopterin (BH4), a cofactor for nitric oxide production, monoamine synthesis, and phenylalanine metabolism). QDPR is also required for repairing oxidized tetrahydrofolate, and lack of QDPR under conditions of oxidative stress can disrupt folate pools (Zheng, Lin et al. 2018). Mergeomics key driver candidates linked to liver iron, TG, copper levels, and red cell traits converged on *FDFT1* and *SQLE*, which encode enzymes that are just up and downstream of squalene in the sterol synthesis pathway. In addition to its role in the synthesis of sterols, squalene affects endoplasmic reticulum membrane stability and lipid droplet size (Ben M’barek, Ajjaji et al. 2017), and in cancer cells, protects against cell death by ferroptosis under oxidative stress (Garcia-Bermudez, Baudrier et al. 2019).

## Discussion

To gain insight into genetic factors that contribute to the variation seen in iron overload and pathology, we measured clinical traits and hepatic mRNA expression in 114 strains of HMDP mice fed a 20k ppm (20,000 ppm iron = 2% carbonyl iron) high iron diet. Across the HMDP, we identified many traits that exhibited high inter-strain variability on the high iron diet, including red blood cell traits, fat mass, liver metals like copper and selenium, and liver and plasma TG and other lipids (Supplementary File 2). We found a substantial contribution of genetics to many traits including body weight and composition, red cell traits, liver iron and copper (Supplementary File 3). Our findings from the HDMP expanded and confirmed previous studies showing that mouse genetic background influences iron and metal parameters (McLachlan, Lee et al. 2011; Cavey, Ropert et al. 2015; McLachlan, Page et al. 2017). In our pilot study with six strains and in a separate Control study in C57BL/6J mice, we observed that the 20k ppm high iron diet induced significant changes in body weight and composition, plasma and liver lipid parameters, liver metals, and hematology (most notably to red cell phenotypes) compared to a 50 ppm iron sufficient control diet.

Using GWAS, we harnessed the variation between strains in the HMDP to identify genetic loci that influence these traits, finding that most clinical traits examined had at least one significant locus (Supplementary File 15). We also measured and genetically mapped liver mRNA expression, identifying *cis*- and *trans*-eQTL for thousands of genes in the context of the high iron diet. By integrated these findings, we prioritized candidate causal genes at loci of interest, identifying TWAS genes at many of these loci. Across all strains, we measured correlation between traits and between traits and mRNA expression in the liver, identifying relationships robust to differing genetic backgrounds. Notably, we identified a strong correlation between liver copper and red cell traits, and a correlation between liver iron and liver TG, that were reflected in shared GWAS loci. Using WGCNA and Mergeomics, we identified key driver genes that are hypothesized to have a particularly important impact on liver iron levels, liver TG, liver copper, and red cell traits. In a commitment to making this data publicly available and usable, we have created a web resource that can be used to explore the data and analyses.

Notably, our study identified top GWAS loci on chromosomes 7 and 11 associated with liver iron levels on the high iron diet. None of these loci harbor genes currently known to be linked to liver iron levels. The chromosome 7 locus at 4.3 to 4.9 Mb overlaps with the sole GWAS locus for liver TG, suggesting that these traits could be related via a common genetic driver. Other HMDP studies with mice fed different diets (mice on chow, high fat/high sucrose, or high cholesterol diets, see resource website), where liver TG levels were measured and mapped, did not have a liver TG GWAS locus in this region, suggesting a gene-environment (diet) interaction. Top candidates genes at this locus based on liver TG TWAS are *Isoc2b*, *Gm15922=Pira1*, and *Rdh13*. The cellular function of Isoc2b is not known, but its human homolog ISOC2 has been shown to functionally interact with and inhibit p16(INK4a), a tumor suppressor and senescence marker (Huang, Shi et al. 2007). *Pira1* is a poorly characterized gene that may function in the immune system. *Rdh13* encodes a mitochondrial protein that catalyzes the oxidation and reduction of retinoids and may protect mitochondria from oxidative stress (Belyaeva, Korkina et al. 2008; Wang, Cui et al. 2012). Ablation of *Rdh13* in mice reduced carbon tetrachloride induced hepatic injury (Cui, Ma et al. 2019). Based on reports in the literature, *Il11* is also a potential candidate gene at this locus. In mice fed diets that induce fibrosis and steatosis, disruption of Il11 signaling decreased these phenotypes (Widjaja, Singh et al. 2019). Furthermore, expression of Il11 in mouse colonic fibroblasts increased in response to oxidative stress, microbiome perturbations (Nishina, Deguchi et al. 2021), and potentially in response to a high iron diet (Chua, Klopcic et al. 2013). Expression of *Il11* at the mRNA level was low or not detected in the mouse livers in this study (Supplementary File 30), so we were not able to determine from our mRNA expression data a link between this locus and this gene. At the chromosome 7 13-15 Mb locus, the top and only candidate based on TWAS is *Selenow*. Selenow is a glutathione-dependent antioxidant enzyme that has been shown to protect cells from oxidative stress (Zhang and Song 2021). At the distal chromosome 11 GWAS peak for liver iron (117,974,693-121,568,141 bp, mm10), TWAS genes in order of increasing TWAS *P* value are *Alyref* (a nuclear molecular chaperone that regulates basic region-leucine zipper proteins and nuclear mRNA export) (Virbasius, Wagner et al. 1999; Okada and Ye 2009), *Gaa* (a lysosomal alpha-glucosidase), *Slc26a11* (a sulfate transporter), *Usp36* (a ubiquitin-specific protease), and *Pcyt2* (involved in phosphatidylethanolamine homeostasis) (Pavlovic and Bakovic 2013). Future studies are needed to functionally validate candidate genes at loci of interest. While we used bulk liver mRNA gene expression to prioritize gene candidates, we note that causal genetic factor(s) may only be evident at the protein level, in a specific cell type, or could be acting via another tissue, e.g. liver iron and TG levels could be influenced by a gene expressed in the intestine.

Previous studies in humans and mice have identified clear or suggestive genetic modifiers of the iron loading disease hereditary hemochromatosis, including genes involved in intestinal iron absorption (*Cybrd1*, *Tf*, *Slc40a1*) and regulation of intestinal iron absorption by hepcidin (*Hamp*, *Hjv*, *Bmp2*, *Bmp6*, *Tmprss6*, *Tfr2*), and novel genes (*Gnpat*, *Pcsk7*, *Arntl*, *Fads2*, *Nat2*). These have been linked to liver iron and other associated traits, including serum iron, transferrin, transferrin saturation and ferritin (Milet, Dehais et al. 2007; Oexle, Ried et al. 2011; Pelucchi, Mariani et al. 2012; Benyamin, Esko et al. 2014; Radio, Majore et al. 2015; Anderson and Bardou-Jacquet 2021). Some additional genes that when modified are known to cause iron overload disorders include *Cp*, *Fxn*, *Pklr*, *HBB*, and *Alas2* (Anderson and Bardou-Jacquet 2021). We did not identify any of these genes at significant or suggestive QTL for liver iron in our study. This indicates that in our study conditions and population, SNPs influencing the expression of these genes were not the most important contributors to liver iron levels.

Candidate modifiers of iron loading and pathology identified in our study may represent modifiers outside the classical genes and regulated pathways involved in intestinal iron absorption. The high iron diet may bypass normal regulatory mechanisms for intestinal iron absorption, entering via a paracellular route. Of note, there was little correlation between liver *Hamp* mRNA expression (the gene encoding hepcidin) and liver iron (bicor = 0.105, *P* = 0.265) across the HMDP. This finding is in accordance with the work of others and previously reported findings from our pilot study, where neither liver *Hamp* mRNA nor plasma hepcidin levels were predictive of tissue iron levels across multiple mouse genetic backgrounds (Cavey, Ropert et al. 2015; McLachlan, Page et al. 2017). Iron overload in the context of anemia likely confounds this correlation, as well as correlations of other genes with expression influenced jointly by iron levels, hypoxia, and anemia. The trait most correlated with *Hamp* mRNA expression in this study is liver copper (bicor = 0.318, *P* = 5.67e-4), followed by positive correlation with red cell traits (HGB, RBC, HCT), none of which have QTL near *Hamp*. Together, this suggests that *Hamp* levels here are more related to the observed anemia than to liver iron levels, and likely in a reactive manner.

Many genes known to influence iron metabolism had significant liver eQTL. These *cis*-eQTL, as well as strong *trans*-eQTL observed for many of these genes, could provide insight into the regulation of these genes at the transcript level in the context of this study. Some of these genes with notable strong *cis*- and or *trans*-eQTL include *Ftl1, Hamp, EpoR, Flvcr1, Atf4*, *Cybrd1*, *Epo*, *Erfe* (*Fam132b*), *Slc25a37*, *Steap2*, *Trf*, *Tfrc*, *B2m, Pcbp2, Hfe, Hjv, Hif1a,* and *Slc39a1*. *Cis*- and *trans*-eQTL for all genes expressed in our study can be explored on the resource website.

In order to identify biological pathways related to liver iron loading, we performed gene enrichment analysis of genes whose expression most highly correlated with liver iron levels across all 114 HMDP strains. Interestingly, enrichments for top genes that negatively correlated with liver iron across the 114 HMDP strains overlapped with pathways associated with differentially expressed genes that were downregulated in C57BL/6J in response to the 20k ppm high iron diet. These included pathways related to key metabolic processes in hepatocytes including lipid and amino acid metabolism, biological oxidations, and response to toxins. In accordance with this, the WGCNA gene expression modules that were most negatively correlated with liver iron across the HMDP, the blue and purple modules, have overlapping pathway enrichment terms with these genes. The downregulation of these genes may potentially be explained by a loss or dysfunction of hepatocytes in the context of iron overload. High iron can result in the death of primary hepatocytes, and iron overload in humans and rodents is linked to hepatocyte death and liver injury, with high iron also exacerbating liver injury induced by other toxins and disease (Chen, Sugiyama et al. 2020; Kouroumalis, Tsomidis et al. 2023). Death occurs by a variety of mechanisms including apoptosis, necrosis, and, as more recently reported, ferroptosis (Zhao, Laissue et al. 1997; Hassannia, Vandenabeele et al. 2019; Wu, Wang et al. 2021). Other studies have questioned the hepatotoxicity of liver iron by itself, based in part on the lack of fibrosis observed in many rodent studies of iron overload and the variability between studies in pathologic outcomes, and have suggested that other factors in addition to high iron may be required (Bloomer and Brown 2019).

Pathways that were associated with differentially expressed genes that were upregulated in C57BL/6J in response to the 20k ppm high iron diet differed from pathways associated with genes positively correlated with liver iron across the full HMDP. The upregulated differentially expressed genes were more related to the immune response, while genes most positively correlated with liver iron across the HMDP were linked to transcription, translation, hypoxia, inflammation, and, in further support of cell death, apoptosis. The WGCNA module most positively correlated with liver iron, the magenta module, shared some terms with this latter group of genes and was also enriched in genes linked to ferroptosis. Looking at individual genes, we saw little correlation between top ranked genes with differential expression in response to the 20k ppm high iron diet in C57BL/6J and genes with the highest correlations with liver iron levels across the HMDP (Supplementary File 37). Top differentially expressed genes in C57BL/6J were also not strong candidate genes associated with liver iron levels by GWAS. These findings suggest that genes with the greatest observed changes in response to high iron, at least in the C57BL/6J strain, are not necessarily those that correlate with and potentially contribute to the variation in iron levels across strains on the high iron diet.

In humans, high liver iron is often found in the context of hepatic steatosis, but the cause(s) of this relationship is not yet clear (Datz, Müller et al. 2017; Rametta, Fracanzani et al. 2020). We observed a spectrum of pathologic findings consistent with varying degrees of hepatic steatosis across the HMDP when examining liver histological sections. We verified these findings with Oil Red O staining of the liver of C57BL/6J mice fed the 20k ppm high iron diet. Compared to the 50 ppm iron sufficient control diet, C57BL/6J mice fed the 20k ppm high iron diet trended toward increased hepatic TG levels, which was also reported recently in other studies of mouse iron overload (Ding, Zhang et al. 2021; Protchenko, Baratz et al. 2021). Two key *de novo* lipogenesis genes were upregulated 2.4 fold, and most fatty acid degradation pathway genes were downregulated, by the 20k ppm high iron diet in the Control study. Lipid droplet formation has been observed in many contexts including mitochondrial dysfunction, liver injury, hypoxia, copper deficiency, and metabolic perturbations (Pressly, Gurumani et al. 2022), and the mechanisms driving this phenotype here are not known. In Drosophila, oxidative stress has been shown to stimulate lipid droplet synthesis, where PUFAs are then redistributed. Lipid droplet formation serves as part of the antioxidant response, as lipids in lipid droplets are better protected from peroxidation than when in cell membranes (Bailey, Koster et al. 2015). In support of oxidative stress driving lipid droplet formation, the antioxidant Vitamin E was shown to ameliorate hepatic steatosis in a *Pcbp1* hepatic knockout mouse model of unchaperoned iron (Protchenko, Baratz et al. 2021). Vitamin E has also been previously shown to decrease steatosis in NAFLD patients through an antioxidant mechanism that decreases hepatocyte *de novo* lipogenesis (Podszun, Alawad et al. 2020). Interestingly, in contrast to hepatocytes, adipose expansion has been reported to be blocked by oxidative stress by suppression of adipocyte *de novo* lipogenesis, leading to ectopic lipid accumulation in the liver (Okuno, Fukuhara et al. 2018). In our Control study in C57BL/6J, the 20k ppm high iron diet led to large hepatic lipid droplets but overall slowed normal accumulation of fat mass. This raises the possibility of a contributing role for adipocyte dysfunction in the hepatic steatosis observed in the context of iron overload. The overall halt in fat mass accumulation is likely also explained by the findings from a recent study in which C57BL/6 mice fed a 2% carbonyl iron diet developed negative energy balance and whole body wasting due at least in part to impaired intestinal lipid absorption (Romero, Mu et al. 2022). Several significant GWAS loci were identified for body composition traits that could be further explored to identify candidate genes that may influence these changes.

We also observed perturbations to other hepatic and plasma lipids in response to the high iron diet. In our Control study with mice fed the 20k ppm high iron diet for 6-7 weeks, there was an increase in liver total cholesterol that could be explained in our study by a similar increase in esterified cholesterol. In contrast, total liver phophatidylcholines were decreased. An increase in liver cholesterol in response to iron overload in AKR mice has been reported previously and potentially explained by an increase in cholesterol biosynthesis gene expression (Graham, Chua et al. 2010), and an increase in hepatic cholesterol was also observed in the *Pcbp1* hepatic knockout mouse model of unchaperoned iron (Protchenko, Baratz et al. 2021). Protchenko et al. also reported altered hepatic phospholipids and increased mRNA expression of genes involved in cholesterol biosynthesis, fatty acid metabolism, and TG synthesis in wild-type C57BL/6J mice fed a 20k ppm high iron diet for ∼3 weeks after weaning (Protchenko, Baratz et al. 2021). In our Control study with mice fed the 20k ppm high iron diet, we did not see significant differences in expression for most of these particular genes, however. Cholesterol biosynthesis gene expression trended toward being decreased compared to control, and there was a mix of up and down trending expression among the fatty acid and TG synthesis genes. Potential diet differences and the different time points (3 weeks vs 6 weeks of iron feeding) at which tissues were collected may explain these differences. We also observed an increase in plasma TG levels and a decrease in plasma cholesterol in all forms measured in C57BL/6J fed the high iron diet compared to control. Liver and plasma lipids have been reported previously to be altered by high iron (dietary, genetic, or injected), but the direction of change is not consistent across studies, and the mechanisms remain unclear (Graham, Chua et al. 2010; Ahmed, Latham et al. 2012; Prasnicka, Lastuvkova et al. 2019; Ding, Zhang et al. 2021). We identified GWAS loci associated with most lipid traits which could provide insight into the relationship between iron overload and lipid homeostasis in the liver.

The high iron diet also led to a wide spectrum of anemia across HMDP strains, ranging from severe to normal range. Based on the pilot and Control study, red blood cells were normal to macrocytic in size compared to those from mice on a control diet. Extramedullary erythropoiesis was also observed histologically in the liver of some strains, and genes regulated by the top eQTL hotspots in the liver were enriched in those involved in erythropoiesis. The severity of the anemia was highly correlated with several traits, including percent fluid mass, heart weight, plasma TG, plasma HDL cholesterol, and hepatic copper. In rats and C57BL/6J mice, a high iron diet can perturb copper absorption and distribution, leading to anemia and cardiac hypertrophy (Ha, Doguer et al. 2016; Ha, Doguer et al. 2017; Ha, Doguer et al. 2018). In those studies, supplementation of high iron diets with 10-25 times normal levels of copper prevented high iron diet induced cardiac hypertrophy and anemia and partially reduced serum nonheme iron levels and improved growth. Supplementation also increased levels of serum ceruloplasmin (Cp), the predominant copper protein in the blood and important for maintaining proper systemic iron distribution. Copper supplementation, however, did not prevent iron-overload induced hepatomegaly or alter TIBC or transferrin saturation. When copper was supplemented in the last 3 weeks of a 7 week high iron feeding regimen, anemia, serum copper levels, and serum Cp ferroxidase activity were normalized, but growth rates, cardiac copper concentrations, and heart size were only partially improved (Wang, Xiang et al. 2018). These findings in previous studies, coupled with the strong correlation between these traits in our HMDP study, strongly suggest that the spectrum of anemia and cardiac hypertrophy observed in the iron loaded HMDP mice were related to copper levels.

Currently, the mechanisms by which the high iron diet could decrease intestinal copper absorption and alter copper distribution (and potentially biliary excretion) are not known. It is currently unclear how copper is taken up into enterocytes (Pierson, Yang et al. 2019). Also unresolved is the mechanism by which copper deficiency can lead to a normo to macrocytic anemia in the context of high transferrin iron saturation. Of note, copper deficiency in people can manifest as megaloblastic or macrocytic anemia (Gregg, Reddy et al. 2002; Hariz and Bhattacharya Updated 2022 Sep 26). The overlapping GWAS loci that we have observed for liver copper and red cell traits RBC, HGB, and HCT may harbor genes that contribute mechanistically to these phenotypes. Although there are several candidate genes at the most prominent overlapping locus on chromosome 9, *Sc5d* and *Sorl1* are the most significant shared TWAS genes for these traits in the overlapping region. *Sc5d* is part of the cholesterol synthesis pathway, where it generates 7-DHC, a lipid highly susceptible to lipid peroxidation (Xu, Korade et al. 2010). 7-DHC was recently shown to protect cellular membranes from (phospho)-lipid peroxidation and associated ferroptotic cell death (Angeli, Freitas et al. 2021; Yamada, Karasawa et al. 2022). *Sc5d* is also one of the top genes downregulated in the liver of mice with defects in copper trafficking due to loss of copper transporter ATP7B (Muchenditsi, Talbot et al. 2021). The other top candidate, *Sorl1*, is a sorting receptor belonging to both the low density lipoprotein receptor (LDLR) family and the vacuolar protein sorting 10 (VPS10) domain-containing receptor family. *Sorl1* is genetically linked to Alzheimer’s disease and is involved in the trafficking of a variety of proteins including many involved in lipid metabolism. It has also been shown to functionally interact with retromer, a complex essential for the proper trafficking of the copper transporters ATP7A, ATP7B, and CTR1 (Fjorback and Andersen 2012; Phillips-Krawczak, Singla et al. 2015; Curnock and Cullen 2020; Das, Maji et al. 2020).

Copper deficiency can also lead to hepatic steatosis and alterations in lipid parameters. Dietary copper restriction in rats leads to hepatic steatosis and insulin resistance (Aigner, Strasser et al. 2010). Low copper and serum CP have also been associated with increased liver iron, steatosis, inflammation, and fasting glucose in NAFLD (Aigner, Theurl et al. 2008; Aigner, Strasser et al. 2010; Wang, Zhou et al. 2022). Copper has also been shown to play an important role in intestinal lipid absorption. Copper, iron, and lipid absorption all occur in the proximal regions of the intestine. Copper excess or deficiency in duodenal enterocyte organoids disrupts chylomicron assembly, and whole-body or intestine specific knockout of the copper transporter ATP7B in mice leads to the buildup of TG-rich vesicles and mislocalization of APOB in intestinal enterocytes (Pierson, Muchenditsi et al. 2018). We hypothesize that some of the differences observed between studies in iron-overload induced alterations in lipid parameters may be influenced by the level of copper in the study diets. While copper deficiency is not noted to be associated with iron loading diseases in humans, circulating CP, which is decreased in copper deficiency and measured more routinely than copper, has been reported to be low in patients with hereditary hemochromatosis (Cairo, Conte et al. 2001) (Lainé, Ropert et al. 2002). Retinopathy in a patient with hemochromatosis and unexplained CP and copper deficiency was hypothesized to be due to an increase in redox-active iron due to deficiency of CP (Bellsmith, Dunaief et al. 2020). *Cp* was also shown in a mouse recombinant congenic linkage study to be an HFE hereditary hemochromatosis modifier gene influencing hepatic iron loading (Gouya, Muzeau et al. 2007). Our findings are consistent with primary iron excess and secondary copper deficiency contributing to the observed liver steatosis in some strains.

In summary, our study provides a broad overview of the genetic architecture of dietary iron overload and pathology in a genetically diverse cohort of inbred mouse strains. We have integrated gene expression and clinical traits using association mapping, correlation structure, and statistical modeling, and made the data available as a publicly accessible resource.

### Limitations

While GWAS can identify novel genetic loci linked to traits and eQTL analyses can aid in prioritizing candidate genes, all candidates require further experimental validation. Gene expression (mRNA) used to prioritize candidate genes was measured in whole liver after 6-7 weeks on the high iron diet. It is likely that gene and protein expression in other tissues, at other time points, and at the single cell level could provide additional insight into genes that influence the phenotypes observed. Banked tissues are available, and we are happy to share and collaborate with others interested in expanding this work. Finally, the high iron diet led to several phenotypes that likely interact and that are not yet well understood in this context, including anemia, copper deficiency, and alterations in lipid metabolism, making it difficult to deconvolute associations in the data.

## Methods

### Mice

The HMDP strains of mice, described previously in detail (Bennett et al, 2010), were bred and maintained at UCLA from breeders purchased from the Jackson Laboratory (Bar Harbor, ME), from the colony of Rob Williams at the University of Tennessee Health Science Center, and from the Division of Laboratory Animal Medicine (DLAM) at UCLA. A limited number of mice used in the HMDP study were obtained directly from the Jackson Laboratory. All mice used in the studies were male. Mice were housed with woodchip bedding on a 12-hour light– dark cycle with ad libitum access to water. Mice were fed a standard rodent chow diet (LabDiet PicoLab Rodent Diet 20, cat #5R53*, St. Louis, MO) ad libitum until being placed on an AIN93M 20k ppm high iron diet (2% carbonyl iron, Dyets, cat#115122, Bethlehem, PA) at approximately 4 weeks of age. Body composition (fat mass, lean mass, and free fluid) was measured just before starting the 20k ppm high iron diet and then again the day before tissue collection by nuclear magnetic resonance (Brüker Biospin Corp., Billerica, MA). Six to seven weeks after being placed on the high iron diet, at approximately 10-11 weeks of age, mice were fasted for ∼4-6 hours and then tissues collected between 10 AM and 5 PM, with ∼90% collected between 10 AM and 2 PM. Mice were anesthetized in a chamber with isoflurane and then blood was collected from the retroorbital plexus or heart into lithium heparin tubes (BD Microtainer, cat#365971) and placed on ice. Once all samples were collected, the blood tubes were spun at 11,000 rpm for 5 minutes, and plasma was removed to new tube, spun again to pellet any remaining cells, and then plasma was aliquoted into microfuge tubes and snap frozen. A small amount (∼20 uL) of blood was also collected in an heparin-coated capillary tube and analyzed immediately for hematology. The body cavity was opened, and mice were perfused via the heart with cold phosphate buffered saline to flush out remaining blood from the tissues. Tissues (heart, liver, spleen, kidneys, duodenum, pancreas, muscle, gonadal fat, full cecum) were then excised and weighed (heart, liver, spleen, left kidney, full cecum). Depending on the tissue, pieces were either formalin-fixed, placed in RNAlater, and/or frozen immediately in liquid nitrogen and stored at −80°C until analysis. In particular for this study, the large lobe of the liver was divided into aliquots for RNA isolation, metals and lipid measurement (snap-frozen), and histology (placed in 10% formalin). On average, total processing time for each mouse (from anesthesia to complete collection) was 6 minutes. Mice that died prematurely or were euthanized early due to fight wounds, hydrocephalus, sickly appearance, or rectal prolapse were excluded from all analyses.

For the Pilot study, six male mice per study group from the A/J, AKR/J, BALB/cJ, C3H/HeJ, C57BL/6J, and DBA/2J strains were obtained from The Jackson Laboratory at 3 weeks of age. Mice were housed in woodchip bedding on a 12-hour light–dark cycle with ad libitum access to water and fed a standard rodent chow diet (LabDiet PicoLab Rodent Diet 20, cat #5R53*, St. Louis, MO) ad libitum upon arrival. At four weeks of age, mice were changed to a 20k ppm high iron diet (2% carbonyl iron, Dyets, cat #115122, Bethlehem, PA) or to a 50 ppm iron sufficient control diet (Dyets cat #515005). After 6 weeks on the defined diets, at 10 weeks of age, mice were fasted for four hours and tissues were collected as described for the HMDP study mice. Samples from these mice were also used in a previously published study (Study 2 in McLachlan et al.) (McLachlan, Page et al. 2017).

In the Control study, male C57BL/6J mice were obtained from The Jackson Laboratory at 3 weeks of age were kept in cages of four mice on the chow diet described above for one week. At that point, two cages were kept on the chow diet, and the other cages were placed on AIN93M defined diets with microcrystalline cellulose as the fiber source (Dyets, Bethlehem, PA, cat #115838 (50 ppm iron sufficient control diet), cat #115839 (500 ppm elevated iron diet), cat #115840 (20k ppm high iron diet = 2% carbonyl iron diet)). Body composition was measured weekly by NMR, with the exception of the week when a glucose tolerance was performed. Fecal samples and physical data from the chow, 50 ppm sufficient control, and 500 ppm elevated iron diets were also included in a previously published microbiome study, with methods described (Thingholm, Rühlemann et al. 2019).

### Food consumption measurement

Food consumption was calculated over the study period for each cage by weighing the remaining food since the last food change. These values were divided by the number of mice in the cage and then averaged to get the average daily food consumption per mouse.

### Tibia length measurement

Soft tissue was removed from tibia, and photographs were taken with a ruler included for scale. Measurements were made using ImageJ.

### Liver metals analysis

Approximately 100 mg of frozen liver from the large lobe of each mouse was excised on dry ice using a clean pre-chilled blade. The piece was transferred to a pre-weighed metal-free 15ml conical tube (Perfector Scientific #2625), weighed in triplicate on a microscale, and then dried at 60°C in an oven. Dry weight was then taken, and samples were sent to the Analytical Toxicology Core Laboratory (ATCL) at the University of Florida, Gainesville. Dried samples were transferred to perfluorooxyalkane (PFA) digestion vessels. Two mL 50% Optima HNO_3_ and 0.2 mL 30% ACS grade H_2_O_2_ (both from Fisher Chemical, Woodlawn, NJ) were added to the samples. Along with every batch of 18 samples, a method blank and a standard reference material (SRM) 1577c Bovine Liver (NIST, Gaithersburg, MD) were prepared. The PFA vessels were sealed and placed into a microwave digestion oven (CEM, Matthews, NC). The digestion program was as follows: a) 1600 W at 80% power, temperature ramp to 200°C within 15 minutes, b) hold at 200°C for 15 minutes, and c) cool down 15 minutes. After cooling, sample digests were quantitatively transferred to polypropylene tubes and diluted with ≥18.2 MΩ·cm water (Millipore Sigma, Burlington, MA) to 16 mL.

An Agilent 7900 inductively coupled plasma mass spectrometer (Santa Clara, CA) with in-line internal standard addition was used for ICP-MS metals analysis for the HMDP and Control study. ICP-AES was used for liver metals analysis in the pilot study as described previously for these samples (Study 2, McLachlan et al.) (McLachlan, Page et al. 2017). The liver iron concentrations from the pilot study were not previously published but were used in statistical analyses to examine the relationship between liver iron and sex, strain, liver hepcidin mRNA expression, and plasma hepcidin in the McLachlan et al. study (McLachlan, Page et al. 2017). For ICP-MS, the instrument was operated in helium gas mode for most metals of interest, which minimized polyatomic interferences; cadmium was the only metal quantitated without helium gas screening. Mixed-metals calibration dilutions ranged from 0.1 to 10,000 ng/mL; resulting calibration regressions were linear with r^2^ ≥ 0.999. Most SRM 1577c measured metals came to within 10% of published values, and 25% was determined as the cut-off limit for accuracy.

### Histology

Mouse livers and femurs were fixed in formalin for 24 hours, and then placed in 70% ethanol for paraffin blocking by the Translational Pathology Core Laboratory (TPCL) at UCLA. Femurs were decalcified with prior to dehydration and blocking. The blocks were cut into 4 μm sections and stained with hematoxylin and eosin (H&E) to visualize morphology and with potassium hexacyanoferrate(II) trihydrate (Sigma cat# P3289) to visualize ferric iron deposits and counterstained with Neutral Red to visualize cell morphology (Sigma-Aldrich cat#72210).

Liver sections from a smaller selection of strains were stained with picrosirius red to specifically examine fibrillar collagen as described previously (Tuominen, Fuqua et al. 2021). Deparaffinized sections were stained with Weigert’s haematoxylin (Sigma-Aldrich) for 8 minutes and washed for 10 minutes under running tap water. They were then stained with 0.1% picrosirius red for one hour (Direct Red 80 and 1.3% picric acid from Sigma-Aldrich) and washed in two changes of acidified water (10 seconds in the first and the remaining 10 minutes in the second, 5 ml of acetic acid per 1 liter of water). Slides were then dehydrated in three changes of absolute ethanol, cleared in two changes of xylene, and mounted with Permount (Fisher Scientific, Pittsburgh, PA). A Zeiss Axioimager and Aperio Scanscope were used for imaging.

To visualize lipids, a small piece of the large lobe of liver was placed in a tissue mold, covered with O.C.T. Compound (Tissue-Tek), and frozen. 10 μm sections were incubated in water for 2 minutes to clear OCT, followed by 7 minutes in Oil Red O (Sigma, cat# O0625, 7 minutes in water, 30 seconds in hemotoxylin, 5 minutes in water, 1 minute in bluing reagent (Fisher, cat# 22-050-115), and 10 minutes in water.

### Lipid quantification

Liver lipids were extracted using the Folch method (Folch, Lees et al. 1957). In brief, an approximately 60 mg liver piece was homogenized in methanol (Fisher Scientific, Waltham, MA); chloroform (Fisher Scientific, Waltham, MA) was then added to lyse cells and dissolve lipids. Proteins were precipitated and the lipids were collected into the lipid-soluble fraction, dried, and dissolved in 1.8% (wt/vol) Triton X-100 detergent (Fisher Scientific, Waltham, MA). Colorimetric assays were used according to the manufacturer’s instructions to quantify liver triglycerides (TG), total (TC) and unesterified cholesterol (UC) (Sigma, St. Louis, MO), and phospholipids-C (PL) (Wako, Richmond, VA). Phospholipids-C includes phosphatidylcholine and lysophosphatidylcholine. Plasma lipids were quantified as previously described (Warnick 1986; Hedrick, Castellani et al. 1993). Esterified cholesterol (EC) was determined by subtracting UC from TC, and plasma low density and very low density lipoprotein (LDL&VLDL) cholesterol was calculated by subtracting plasma HDL cholesterol from plasma TC.

### Glucose and insulin measurements

Plasma insulin was measured using an ELISA insulin kit (Alpco catalog # 80-INSMSU-E10). Plasma glucose was measured by an enzymatic assay (Stanbio, catalog # 1070-125). Homeostatic model assessment of insulin resistance (HOMA-IR) was calculated using the equation: (glucose × insulin)/(405) where glucose is in mg/dl and insulin is in mU/L (Bowe, Franklin et al. 2014). Insulin in pg/mL was multiplied by a conversion factor of 0.025 to get mU/L, using 1 mIU/L = 6.00 pmol/L insulin (Knopp, Holder-Pearson et al. 2019).

In the Control study, a glucose tolerance test (GTT) was performed after mice had been on study diets for 3-4 weeks as previously described (Thingholm, Rühlemann et al. 2019). The area under the glucose curve (AUC) for each mouse was calculated using the formula: AUC = 0.25 (fasting value) + 0.5 (30 minute value) + 0.75 (60 minute value) + 0.5 (120 minute value) per the Schonfeld1 project protocol (https://phenome.jax.org/projects/Schonfeld1/protocol). Blood glucose was also measured using the glucometer at 10:30am after a 5.5 hr fast prior on the day of tissue collection, and plasma glucose was measured as described above.

### Hematology

Hematological parameters were measured using a Heska Hematrue Veterinary Hematology Analyzer (Loveland, CO) following the manufacturer’s instructions. Anticoagulated blood was stained with Wright-Giemsa and smears were prepared by the UCLA DLAM Diagnostic Lab.

### Genome-wide association analysis and heritability estimation

Mice were genotyped using the Mouse Diversity Array (Rau, Parks et al. 2015), and informative SNP markers (obtained after filtering for 5% minor allele frequency and 10% missingness rate) were used in genetic mapping analyses. Traits were quantile transformed to normalize the distribution and then the FaST-LMM program (Lippert, Listgarten et al. 2011) was used to perform GWAS. To identify eQTL, liver mRNA expression was mapped from log_2_-transformed transcripts per million (TPM) values using FaST-LMM. Manhattan plots were generated with the R package qqman (Turner 2018) and loci were visualized with LocusZoom (Pruim, Welch et al. 2010) and the IGV app (https://igv.org/app/). The R package ‘heritability’ was used to calculate broad-sense heritability (H^2^), and the Genome-wide Complex Trait Analysis Tool (GCTA) (Yang, Lee et al. 2011) was used to calculate narrow-sense heritability (h^2^).

### Global gene expression and differential expression analyses

Total RNA from a 20 mg piece of the large lobe of the liver was purified from one mouse per strain of the HMDP, and from 3 mice per diet group in the C57BL6J Control study, using the Qiagen miRNeasy Mini kit (Qiagen cat#217004) per the manufacturer’s instructions. RNA preparation was as previously described (Ma, Fuqua et al. 2019) except that samples were homogenized in QIAzol using a TissueLyzer (Qiagen). All samples had RINe values greater than 8. RNA sequencing libraries were prepared as described previously (Ma, Fuqua et al. 2019) using the KAPA Stranded mRNA-Seq Kit (cat #KK8421, KAPA Biosystems, Wilmington, MA). The pooled libraries were sequenced in an Illumina HiSeq4000 instrument (Illumina, San Diego, CA). Reads were quantified against the GRCm38.p6 mouse reference transcriptome (Ensembl release 97) using kallisto version 0.46.0 with 100 bootstrap replicates. Differential expression analysis of RNA expression from 3 mice each in the 50 ppm iron and 20k ppm iron diet groups in the C57BL/6J Control study was performed using DESeq2 (Love, Huber et al. 2014). For all analyses except differential expression (where program default low abundance filtering parameters were used), only genes with expression values > 0.1 TPM in at least 20% of samples and ≥ 6 reads in at least 20% of samples were included.

### Pathway and network modeling

Correlations between traits and gene expression were analyzed using the bicor function in the weighted gene co-expression network analysis (WGCNA) R package (Langfelder and Horvath 2008). Metascape (https://metascape.org) was used to perform pathway analysis of top correlated genes using human pathway settings (Zhou, Zhou et al. 2019).

As previously described, to calculate the *cis*-component of expression correlated with traits, the expression values for each relevant gene with a significant cis-eQTL (*P* < 1e-4) were divided into groups based on the genotype of their most significant *cis*-eQTL (Tuominen, Fuqua et al. 2021). Median expression values were calculated for each group, and the medians (replicated once for each individual within each group) as a whole were correlated against the trait. Genes whose local, or *cis,* component of expression variance correlates with a trait with a bicor correlation *P* value < 0.05 are considered “TWAS genes” for that trait.

Correlated gene network analysis was performed similarly to as previously described (Tuominen, Fuqua et al. 2021) using the WGCNA R package (Langfelder and Horvath 2008), which groups highly co-expressed genes into modules. We first discarded genes with expression not meeting the expression cutoffs described above to get a subset of 15,499 genes. To generate a signed hybrid co-expression network (where only genes that are positively correlated with each other are grouped together into a module), an adjacency matrix was created by calculating pairwise gene-gene correlations using the log (TPM+1) for each gene. The resulting bicor correlations were raised to the 8th power, which was selected using the scale-free topology criterion. MaxPOutliers was set to 0.05. A topological overlap matrix-based dissimilarity measure was used for hierarchical clustering of the genes. Gene modules corresponded to the branches of the resulting dendrogram and were defined using the “Dynamic Hybrid” branch cutting algorithm. For module generation, “merge cut height” was set to 0.25 and “minimum module size” to 30. To identify gene clusters associated with traits, the first principal component of each module was analyzed for bicor correlation with traits. The first principal components were also mapped using FaST-LMM.

For Mergeomics, the full summary statistics of the trait-associated GWAS SNPs was used as the initial input. Next, we carried out marker dependency filtering (MDF) on the Mergeomics webserver (Arneson, et al., 2016) to select independent SNPs from LD blocks, with LD defined here as correlated SNPs with r^2^ > 0.7. Only the top 50% of SNPs in association with the trait were kept after MDF. Marker set enrichment analysis (MSEA) and key driver analysis (KDA) were done with the Mergeomics R package (Shu, Zhao et al. 2016; Ding, Blencowe et al. 2021) with default parameters except for trim, which was set to 0.01. The trim_start parameter was tuned by running MSEA with different values using a negative control (1000 gene sets randomly assigned to genes) until approximately 5% of the randomly assigned gene sets reached 5% FDR. The trim_end parameter for MSEA was simply 1 minus trim_start.

MSEA was used to find pathways enriched among genes associated with the trait. We assigned genes to trait GWAS SNPs using liver *cis*-eQTLs (*P* < 1e-5) obtained from gene expression in this HMDP study. Then we searched for enrichment in canonical pathways from KEGG (Ogata, et al., 1999), Reactome (Croft, et al., 2014), and Biocarta. KDA was used to identify candidate key driver genes using gene regulation Bayesian networks from a previous study (Zhao, et al., 2016) supplemented with a network generated from our gene expression data. Cytoscape (version 3.5.1) was used to visualize the key drivers and their neighborhoods.

### Statistical analyses

Simulation studies were previously used to test the statistical power of the HMDP using parameters including the variance explained by SNPs, genetic background, random errors, and the number of repeated measurements per strain (Bennett, Farber et al. 2010). Appropriate sample size to achieve adequate statistical power for all analyses was determined based on previous HMDP studies. Differences in sample sizes among the HMDP strains were due to differences in strain availability as determined by breeding success and losses. The initial number of mice per group in the pilot (N = 6 per group) and Control studies (N = 8 per group) were determined based on previous studies where similar phenotypes were measured. In the pilot study, one mouse died prematurely in the A/J high iron group, so the final N = 5 for that group.

Analyses were performed using GraphPad Prism (GraphPad Software, La Jolla, CA) and in R. *P* < 0.05 was considered significant for these tests and for bicor analyses. *P* < 1e-4 was considered significant for *cis*-eQTL, and P <1e-6 for *trans*-eQTL. For GWAS, thresholds for significant (*P* < 4.1e-6; -log_10_*P* > 5.387) and suggestive (*P* < 4.1e-5; -log_10_*P* > 4.387) loci were defined using simulation. All reported *P* values are based on a two-sided hypothesis. All N values represent the number of unique biological replicates or, as noted, the number of unique summary data replicates (e.g. trait means). All experiments were performed once. Missing values (NA) are present in the body and tissue weights when weight was inadvertently not recorded, in the hematological data when the data was not provided by the analysis instrument, in the ICP-MS metals measurements if the metal measurement could not be determined or failed the reference standard criteria for the run, and in the liver lipids and plasma glucose, insulin, and lipid data if the sample was not measured. In order to minimize missing values, in cases where sample results were extreme outliers or there was a technical issue with the data collection, analyses were repeated if possible and data was replaced. Unless noted, outliers were not removed in data analyses.

### Ethics statement

All animal work was approved by the University of California, Los Angeles, Animal Research Committee, the institutional animal care and use committee, in accordance with PHS guidelines and under ARC # 1992–169. In vivo experiments are reported in accordance with the ARRIVE guidelines. All authors had access to the study data and reviewed and approved the final manuscript.

## Abbreviations

AUC: Area under the curve
bicor: Biweight midcorrelation
CI: classical Inbred
DE: differentially expressed
EC: esterified cholesterol
eQTL: expression quantitative trait locus/loci
FDR: false discovery rate
GWAS: genome-wide association study
GTT: glucose tolerance test
HCT: hematocrit
H&E: hematoxylin and eosin
HGB: hemoglobin
HDL: high-density lipoprotein
HOMA-IR: homeostatic model assessment of insulin resistance
HMDP: Hybrid Mouse Diversity Panel
ICP-AES: inductively coupled plasma atomic emission spectroscopy
ICP-MS: inductively coupled plasma mass spectrometry
KDA: key driver analysis
LD: linkage disequilibrium
LDL: low density lipoprotein
MDF: marker dependency filtering
MSEA: marker set enrichment analysis
MCH: mean cell hemoglobin
MCHC: mean cell hemoglobin
MCV: mean cell volume
MPV: mean platelet volume
Mb: megabase
NAFLD: non-alcoholic fatty liver disease
PL: phospholipids-C
PCA: principal component analysis
QTL: quantitative trait locus/loci
RI: recombinant Inbred
RBC: red blood cell count
RDW: red cell distribution width
RDWa: red cell distribution width absolute
RNA-Seq: RNA sequencing
SNP: single nucleotide polymorphism
SD: standard deviation
SEM: standard error of the mean
SRM: standard reference material
TC: total cholesterol
TPM: transcripts per million
TG: triglyceride
UC: unesterified cholesterol
VLDL: very low density lipoprotein
WGCNA: weighted gene co-expression network analysis

## Newly created materials availability

Banked tissues are available for future collaborative research upon reasonable request.

## Data and code availability

The liver gene expression data generated in this study is available at www.ncbi.nlm.nih.gov/geo/, accession GSE230674. Mouse phenotype data, genetic mapping results, correlations, and interactive visualizations of this data are available at the resource website: https://systems.genetics.ucla.edu/HMDP/. All other data and code is available upon reasonable request.

## Author contributions

A.J.L., B.K.F., C.D.V., E.E., and S.M. conceived the study. A.M., B.K.F, B.O., C.H., C.N., C.P., C.L.S., D.W.K., H.I., J.Z., K.P, K.T, L.M, M.B., M.K., N.C., N.K., N.L., R.C.D., S.C., S.T.H., S.M., T.L., Z.S., and Z.Z. performed experiments and/or analyzed the data. X.Y. and D.M.F. provided consultation. B.K.F., L.M., A.J.L. and C.D.V. drafted the manuscript, and all authors read or revised the manuscript.

## Figure and supplementary file legends

**Figure 1-figure supplement 1.**
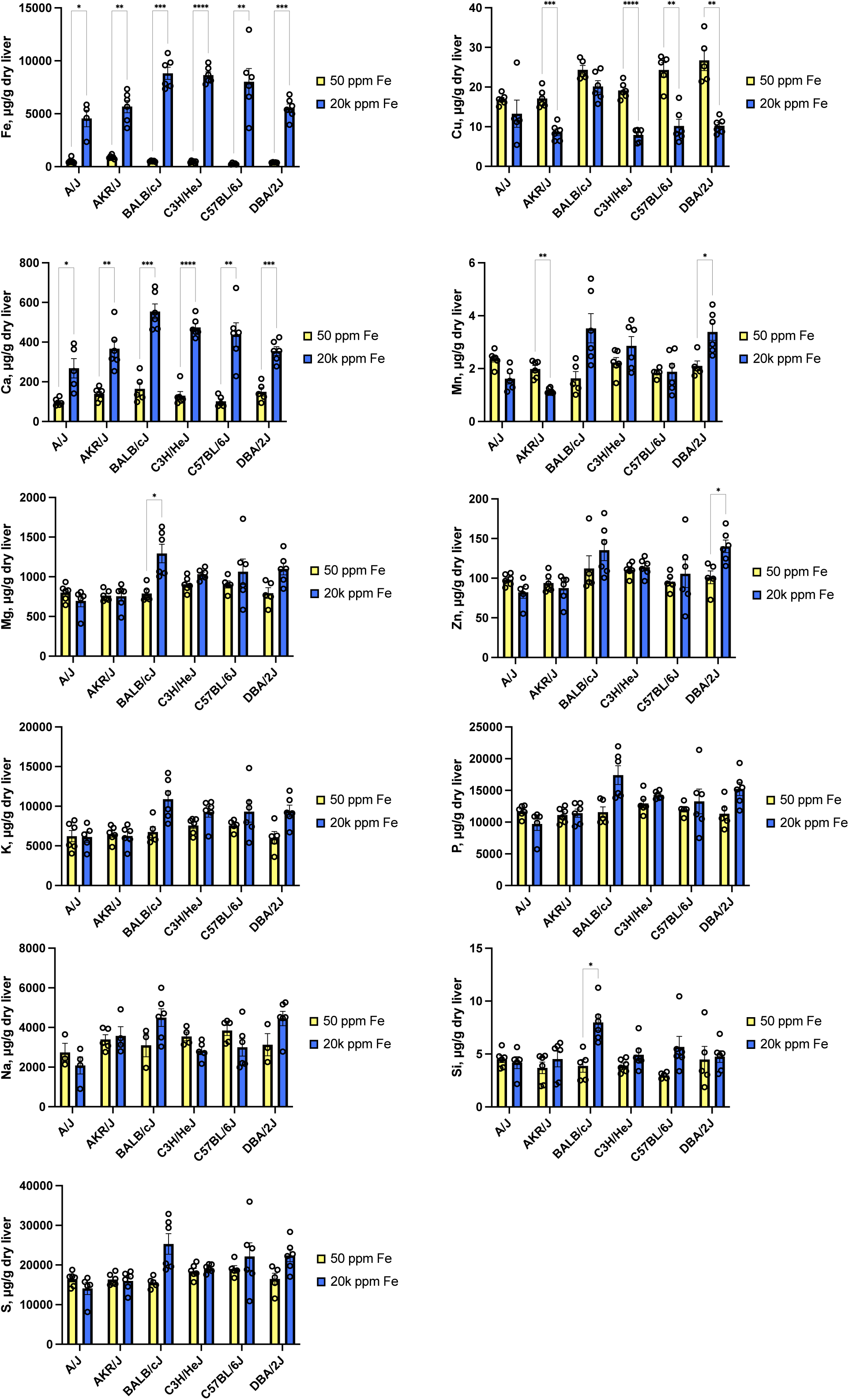
Liver metals in the pilot study as measured by ICP-AES. Barplots of all pilot study liver metals, expressed as μg metal per g dry tissue for mice fed the 50 ppm iron sufficient control diet and the 20k ppm high iron diet. Statistical comparisons are between mice of the same strain on different diets. Differences that were significant (*P* ≤ 0.05 using unpaired t tests with Welch’s correction and the Holm-Šídák multiple comparisons test for each metal in GraphPad Prism v9) are denoted with \**P* ≤ 0.05; \*\**P* ≤ 0.01 ; \*\*\**P* ≤ 0.001 ; \*\*\*\**P* ≤ 0.0001. N = 4-6 mice per group, mean and SEM. This data corresponds to data in Figure 1-Source Data 1.

**Figure 1-Source Data 1.** Liver metal measurements in the pilot study. Liver metal concentrations for mice fed a 50 ppm iron sufficient control diet and a 20k ppm high iron diet were measured by ICP-AES. Liver metals are in μg/g (ppm) dry liver tissue. The summary data reports the mean ± standard error of the mean (SEM) (N=number of mice in the group). Fold changes between the diets for each strain and trait are calculated as mean level in the high iron diet group divided by the mean level in the sufficient iron diet group. For fold change, an asterisk (*) notes *P* ≤ 0.05 using unpaired t tests with Welch’s correction and the Holm-Šídák multiple comparisons test for each trait in GraphPad Prism v9. This data is visualized in Figure 1 and the Figure 1-figure supplement 1.

**Supplementary File 1.** Mouse strains in the HMDP study, with the Jackson Laboratory (JAX) strain number, RRID, strain type and subtype, number of mice, and number of cages.

**Figure 1-Source Data 2.** Data file with liver iron levels (ppm, normalized to dry liver weight) for each HMDP mouse.

**Supplementary File 2.** HMDP study trait individual mouse and summary data, with separate tabs for strain mean, number of mice (N), standard deviation, and median, and for individual mouse data. The key tab gives further information about this table. A summary across all strains (or across all mice for the individual mouse data sheet) is in the last rows of each sheet.

**Supplementary File 3.** Estimated broad sense (H^2^) and narrow xsense (h^2^) heritability for traits in this study.

**Supplementary File 4.** Liver metals in the C57BL/6J Control study as measured by ICP-MS. Barplots of all liver metals for Control study C57BL/6J mice fed the chow diet, 50 ppm iron sufficient control diet, 500 ppm iron elevated diet, and the 20k ppm high iron diet. Statistical comparisons are between all groups except the chow diet group. Differences that were significant (by one-way Brown-Forsythe and Welch’s ANOVA test with Dunnett’s T3 multiple comparisons test in GraphPad Prism v9) are denoted with \**P* ≤ 0.05; \*\**P* ≤ 0.01 ; \*\*\**P* ≤ 0.001; \*\*\*\**P* ≤ 0.0001. N = 8 mice per group, mean and SEM. This data corresponds to data in Supplementary File 5.

**Supplementary File 5.** Individual mouse trait and summary data for the C57BL/6J Control study mice on the chow diet, 50 ppm iron sufficient control diet, 500 ppm elevated iron diet, and the 20k ppm high iron diet. Data includes individual mouse glucose tolerance test data, body weight and composition, tissue weights, complete blood count data, left tibia measurements, and liver metals. The glucose tolerance test, body composition, and cecum weight summary data for the 50 ppm Fe sufficient iron diet and the 500 ppm elevated iron diet fed mice were previously plotted and statistically compared as part of a microbiome study (Figure S3 and Table S6 in Thingholm, Rühlemann et al. 2019), with no significant differences found in these parameters between these two groups of mice.

**Figure 2-Source Data 1.** Individual mouse trait and summary data for the C57BL/6J Control study mice on the chow diet, 50 ppm iron sufficient control diet, 500 ppm elevated iron diet, and the 20k ppm high iron diet. Data includes liver lipids and plasma lipids, glucose, and insulin. Plasma glucose, insulin, and plasma total cholesterol and HDL cholesterol summary data for the 50 ppm Fe sufficient iron diet and the 500 ppm elevated iron diet fed mice were previously plotted and statistically compared as part of a microbiome study (Figure S3 in Thingholm, Rühlemann et al. 2019), with no significant differences found in these parameters between these two groups of mice.

**Figure 2-Source Data 2.** Data file with liver TG levels (mg/g wet liver weight) for each HMDP mouse. No data points were excluded; lipids from samples with no data (NA) were neither extracted nor analyzed due to time constraints.

**Supplementary File 6.** Liver total cholesterol and phospholipid-C levels across the HMDP. Individual value plots of liver total cholesterol (TC, panel A) and phospholipid-C (PL, panel B), N = 1-10 mice per strain, for the 114 HMDP strains fed the 20k ppm high iron diet, ordered by increasing strain mean. The color of each dot represents the cage number for a given mouse. Cages were sequentially ordered, so dots with similar colors indicate mice housed in the vivarium around the same dates. This data corresponds to data in Supplementary File 2.

**Supplementary File 7.** Liver unesterified and esterified cholesterol levels across the HMDP. Individual value plots of liver unesterified cholesterol (UC, panel C) and esterified cholesterol (EC, panel D), N = 1-10 mice per strain, for the 114 HMDP strains fed the 20k ppm high iron diet, ordered by increasing strain mean. The color of each dot represents the cage number for a given mouse. Cages were sequentially ordered, so dots with similar colors indicate mice housed in the vivarium around the same dates. This data corresponds to data in Supplementary File 2.

**Supplementary File 8.** Red blood cell count and hemoglobin levels across the HMDP. Individual value plot of red blood cell count (RBC) and hemoglobin (HGB), N=1-11 mice per strain, for the 114 HMDP strains fed the 20k ppm high iron diet, ordered by increasing strain mean. The color of each dot represents the cage number for a given mouse. Cages were sequentially ordered, so dots with similar colors indicate mice housed in the vivarium around the same dates. This data corresponds to data in Supplementary File 2.

**Supplementary File 9.** Effect of the high iron diet on red blood cells and platelets in the pilot study. Barplots of pilot study red blood cell and platelet trait data for the mice fed the 50 ppm iron sufficient control diet and the 20k ppm high iron diet. Statistical comparisons are between mice of the same strain on different diets. Differences that were significant (*P* ≤ 0.05 using unpaired t tests with Welch’s correction and the Holm-Šídák multiple comparisons test for each metal in GraphPad Prism v9) are denoted with \**P* ≤ 0.05; \*\**P* ≤ 0.01 ; \*\*\**P* ≤ 0.001 ; \*\*\*\**P* ≤ 0.0001. N = 5-6 mice per group, mean and SEM. This data corresponds to data in Supplementary File 10.

**Supplementary File 10.** Complete blood count measurements in the pilot study. Blood count data was measured using a veterinary hematology analyzer. The summary data reports the mean ± standard error of the mean (SEM) (N=number of mice in the group). Fold changes between the diets for each strain and trait are calculated as mean level in the high iron diet group divided by the mean level in the sufficient iron diet group. For fold change, an asterisk (*) notes *P* ≤ 0.05 using unpaired t tests with Welch’s correction and the Holm-Šídák multiple comparisons test for each trait in GraphPad Prism v9. This data is visualized in Supplementary File 9 and Supplementary File 11.

**Supplementary File 11.** Effects of the high iron diet on white blood cells in the pilot study. Barplots of pilot study white blood cell trait data for mice fed the 50 ppm iron sufficient control diet and the 20k ppm high iron diet, with statistical comparisons between mice of the same strain on different diets. Differences that were significant (*P* ≤ 0.05 using unpaired t tests with Welch’s correction and the Holm-Šídák multiple comparisons test for each metal in GraphPad Prism v9) are denoted with \**P* ≤ 0.05; \*\*\**P* ≤ 0.001. N = 5-6 mice per group, mean and SEM. This data corresponds to data in Supplementary File 10.

**Supplementary File 12**. Effect of the high iron diet on blood cells and platelets in the C57BL/6J Control study. Barplots of complete blood count trait data for mice fed the 50 ppm iron sufficient control diet and the 20k ppm high iron diet. Diet groups were statistically compared using the unpaired t test with Welch’s correction in Graphpad Prism v9. Mean and SEM, N=8 mice per group. This data corresponds to data in Supplementary File 5.

**Supplementary File 13.** Effect of the high iron diet on body mass and composition. (***A***) Individual value plot of the ratio of the last to first total body mass as measured by NMR (N = 1-11 mice per strain) for the 114 HMDP strains fed the 20k ppm high iron diet, ordered by increasing strain mean. The color of each dot represents the cage number for a given mouse. Cages were sequentially ordered, so dots with similar colors indicate mice housed in the vivarium around the same dates. (***B***) Individual value plot of the ratio of the average daily food consumption per mouse for the 114 HMDP strains while fed the 20k ppm high iron diet, ordered by increasing strain mean. Each dot represents the average food consumption per mouse in a given cage over the study period (N = 1-4 cages per mouse strain). The dots are colored by the average body weight of the mice in the cage at the end of the study period. (***C***) Time course of body weight, fat mass, lean mass, and free fluid in grams as measured by NMR for C57BL/6J mice fed the 50 ppm iron sufficient control diet (orange line) and 20k ppm high iron diet (blue line) in the Control study. Mean and SEM are plotted, N = 8 mice per strain. Timepoints with significant differences between groups (*P* ≤ 0.05 using unpaired t tests with Welch’s correction and the Holm-Šídák multiple comparisons test for each metal in GraphPad Prism v9) are denoted with an asterisk. The data presented in figure panels A and B corresponds to data in Supplementary File 2, and the data presented in figure panel C corresponds to data in Supplementary File 5. The body composition summary data for the 50 ppm Fe sufficient iron diet fed mice were previously plotted and statistically compared with data from another group not plotted here, mice fed the 500 ppm elevated iron diet, as part of a microbiome study (Figure S3 and Table S6 in Thingholm, Rühlemann et al. 2019), with no significant differences found in these parameters between those two groups of mice.

**Supplementary File 14.** Body and tissue weights, tibia length, and glucose measurements in the C57BL/6J Control study. Barplots for C57BL/6J mice fed the 50 ppm iron sufficient control diet and the 20k ppm high iron diet. Diet groups were statistically compared using the unpaired t test with Welch’s correction in Graphpad Prism v9. N = 8 mice per group, mean and SEM. Full cecum weight refers to the weight of the cecum including its contents, and GTT AUC is an abbreviation for “glucose tolerance test area under the curve”. Fasting glucose shown in the final panel was measured with a glucometer on the morning of tissue collection at the study endpoint, while GTT fasting glucose was measured prior to the GTT test, which was performed after 3-4 weeks on the study diet. This data corresponds to data in Supplementary File 5. The glucose tolerance test and cecum weight summary data for the 50 ppm Fe sufficient iron diet fed mice were previously plotted and statistically compared with data from another group not plotted here, mice fed the 500 ppm elevated iron diet, as part of a microbiome study (Figure S3 and Table S6 in Thingholm, Rühlemann et al. 2019).

**Figure 3-Source Data 1.** Trait bicor correlation matrix for 80 HMDP traits. Correlation was calculated across strain trait means using the bicor function in the WGCNA R package, using pairwise complete observations with a maxPOutliers setting of 0.05. This data is visualized in Figure 3.

**Figure 4-figure supplement 1.**
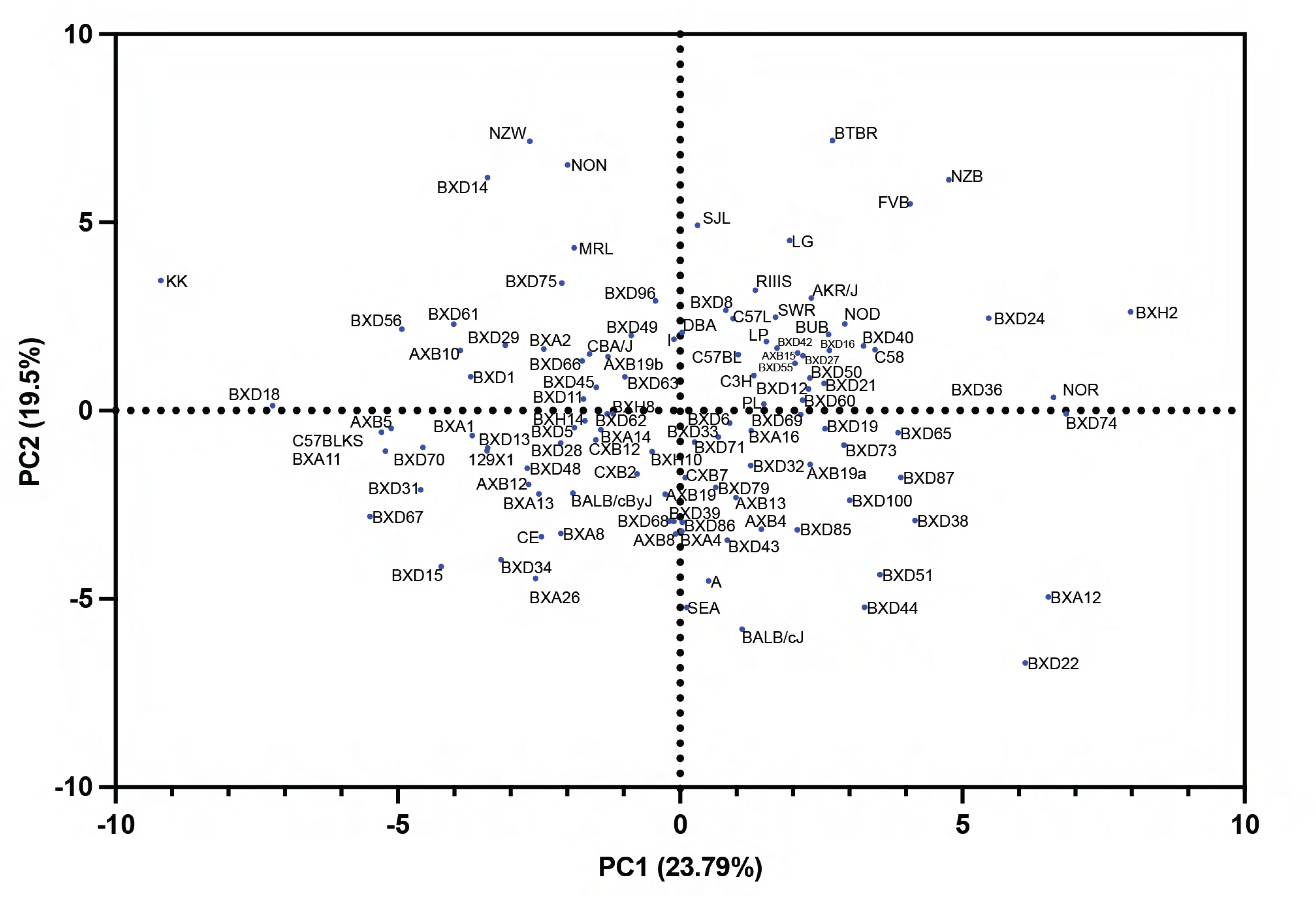
Principal component analysis (PCA) of 41 traits that had strain mean data for all 114 strains. PCA score plot of the 114 HMDP mouse strains across PC1 and PC2. Trait means were z-score standardized as part of the analysis in GraphPad Prismv9.

**Figure 4-Source Data 1.** Principal component analysis (PCA) loadings/correlation matrix for 41 traits and the first five principal components (PCs). For standardized data, the loadings matrix is identical to the correlation matrix between variables and PCs. Each row represents one trait, and each column represents one PC. Each value is the corresponding correlation coefficient between the trait and the PC.

**Figure 4-Source Data 2.** Contribution of strains to each of the first five principal components (PCs) from the PCA analysis. Each row represents one case (row in the original data table), and each column represents one PC. The values in a single column represent the fraction of total variance explained by that PC that each case (strain) contributes. As such, these values sum to 1.0 (100% of the variance explained by the PC). In the original data table used for the analysis in GraphPad Prism v9, each row included the mean trait value for a given mouse strain for 41 traits for which we had data for all 114 strains.

**Supplementary File 15.** Mapped traits, with the total number of strains and total number of mice available for the mapping of each trait. Whether the trait has GWAS suggestive loci and/or GWAS significant loci is noted, as well as notes on the mapping results.

**Supplementary File 16.** Plasma lipid Manhattan plots. Manhattan plots from GWAS of plasma lipid traits. Loci with SNPs above the red bar (-log_10_*P* > 5.387) are statistically significant, while loci above -log_10_*P* > 4.387 are suggestive. Plots are of GWAS results from plasma esterified cholesterol (EC) concentration, the percent of plasma total cholesterol (TC) that in high density lipoprotein cholesterol (HDL), plasma unesterified cholesterol (UC) concentration, the percent of plasma TC that is EC concentration, plasma triglyceride (TG) concentration, plasma HDL concentration, plasma very low density lipoprotein (VLDL) and low density lipoprotein (LDL) concentration, and plasma TC concentration.

**Supplementary File 17.** Body composition Manhattan plots. Manhattan plots from GWAS of body composition traits at the study endpoint. Loci with SNPs above the red bar (-log_10_*P* > 5.387) are statistically significant, while loci above -log_10_*P* > 4.387 are suggestive. Plots are of GWAS results of total body mass, fat mass, lean mass, free fluid mass, the percent of total body mass that is free fluid, the percent of total body mass that is fat, and the percent of total body mass that is lean.

**Supplementary File 18.** Liver lipid Manhattan plots. Manhattan plots from GWAS of liver lipid traits. Loci with SNPs above the red bar (-log_10_*P* > 5.387) are statistically significant, while loci above -log_10_*P* > 4.387 are suggestive. Plots are of GWAS results of liver esterified cholesterol (EC) concentration, liver total cholesterol (TC) concentration, liver unesterified cholesterol (UC) concentration, the percent of liver TC that is EC, liver triglyceride (TG) concentration, and liver phospholipid-C (PL) concentration.

**Supplementary File 19.** Body and tissue weight Manhattan plots. Manhattan plots from GWAS of body and tissue weights. Loci with SNPs above the red bar (-log_10_*P* > 5.387) are statistically significant, while loci above -log_10_*P* > 4.387 are suggestive. Plots are of GWAS results of body weight, liver weight, heart weight, spleen weight, left kidney weight, and full cecum weight (weight of excised cecum including contents).

**Supplementary File 20.** Red cell trait Manhattan plots. Manhattan plots from GWAS of red blood cell traits. Loci with SNPs above the red bar (-log_10_*P* > 5.387) are statistically significant, while loci above -log_10_*P* > 4.387 are suggestive. Plots are of GWAS results of red blood cell count (RBC), mean cell volume (MCV), hematocrit (HCT), mean cell hemoglobin concentration (MCHC), hemoglobin (HGB), mean cell hemoglobin (MCH), red cell distribution width percent (RDW), red cell distribution width absolute (RDWa).

**Supplementary File 21.** Platelet, WBC, and plasma glucose and insulin trait Manhattan plots. Manhattan plots from GWAS of platelet, white blood cell, plasma glucose and plasma insulin traits. Loci with SNPs above the red bar (-log_10_*P* > 5.387) are statistically significant, while loci above -log_10_*P* > 4.387 are suggestive. Plots are of GWAS results of platelet count, mean platelet volume, white blood cell count (WBC), lymphocyte count (LYM), monocyte count (MONO), granulocyte count (GRAN), plasma glucose, and plasma insulin.

**Supplementary File 22.** Liver metals 1 Manhattan plots. Manhattan plots from GWAS of liver metals. Loci with SNPs above the red bar (-log_10_*P* > 5.387) are statistically significant, while loci above -log_10_*P* > 4.387 are suggestive. Plots are of GWAS results of dried liver concentrations of iron, copper, magnesium, manganese, zinc, selenium, potassium, and rubidium.

**Supplementary File 23.** Liver metals 2 Manhattan plots. Manhattan plots from GWAS of liver metals. Loci with SNPs above the red bar (-log_10_*P* > 5.387) are statistically significant, while loci above -log_10_*P* > 4.387 are suggestive. Plots are of GWAS results of dried liver concentrations of sodium, cadmium, cobalt, and strontium.

**Supplementary File 24.** Supplemental text describing 1. *Trans*-eQTL at the overlapping chromosome 7 locus for liver iron and TG; 2. Additional trait QTL and eQTL of interest; and 3. Top WGCNA eigengene mapping results for the green module.

**Figure 5-Source Data 1.** Mapping results of quantile transformed liver iron concentrations. The .gwas file can be opened in a spreadsheet format such as Microsoft Excel and has columns listing each SNP ID, the chromosome that the SNP is located on (CHR), the base pair in mm10 for the SNP (BP), the P value (P), and the Beta. The file can also be interactively opened and viewed at https://igv.org/app/.

**Figure 5-Source Data 2.** Mapping results of quantile transformed liver TG concentrations. The .gwas file can be opened in a spreadsheet format such as Microsoft Excel and has columns listing each SNP ID, the chromosome that the SNP is located on (CHR), the base pair in mm10 for the SNP (BP), the P value (P), and the Beta. The file can also be interactively opened and viewed at https://igv.org/app/.

**Supplementary File 25.** LocusZoom plots of the (*A*) chromosome 7 13-15 Mb and (*B*) the chromosome 11 119-121 Mb liver iron GWAS significant loci. The r^2^ measure of linkage disequilibrium (LD) with the labeled SNP in purple is indicated by the color of the SNP, as shown in the key.

**Supplementary File 26.** The impact of different minor allele frequency (MAF) thresholds on GWAS for liver iron. Manhattan plots are shown for liver iron GWAS that were performed using the standard 5% MAF, as well as the more stringent 10% MAF and 20% MAF cutoffs. SNPs above the red bar (-log_10_*P* > 5.387) are statistically significant, while SNPs above the blue bar (-log_10_*P* > 4.387) are suggestive.

**Supplementary File 27.** Strain subset Manhattan plots for liver iron. Manhattan plots from the liver iron GWAS using all 114 strains (top), the AXB/BXA strains and their two founder strains (top middle), the BXD strains and their two founder strains (bottom middle), and all RI strains with their founders (bottom). The number of strains for each GWAS are noted in the plot titles. Loci with SNPs above the red bar (-log_10_*P* > 5.387) are statistically significant, while loci above the blue bar (-log_10_*P* > 4.387) are suggestive.

**Supplementary File 28.** Strain subset Manhattan plots for liver TG. Manhattan plots from the liver TG GWAS using all 114 strains (top), the AXB/BXA strains and their two founder strains (top middle), the BXD strains and their two founder strains (bottom middle), and all RI strains with their founders (bottom). The number of strains for each GWAS are noted in the plot titles. Loci with SNPs above the red bar (-log_10_*P* > 5.387) are statistically significant, while loci above the blue bar (-log_10_*P* > 4.387) are suggestive.

**Supplementary File 29.** Table of TWAS genes. Genes whose local, or *cis,* component of expression variance correlates with a trait with a bicor correlation *P* value < 0.05 are considered “TWAS genes” for that trait. Only genes with a cis-eQTL (*P* < 1e-4) are included in the analysis, and only the most significant cis-eQTL SNP for each gene is used in the analysis. For each TWAS gene for a trait, the Ensembl gene ID, gene symbol, gene description, gene chromosome, base pair (mm10) where the gene starts, and the base pair (mm10) where the gene ends are listed in columns A-F. The TWAS SNP rsID, SNP chromosome, and SNP base pair location (mm10) are noted in columns G-I. The trait that the gene is a TWAS gene for, followed by the bicor correlation value and *P* value for the correlation of the trait with the cis-component of the expression variance, are noted in columns J-L.

**Supplementary File 30.** List of genes in the chromosome 7 significant GWAS locus for liver iron and liver TG with data from some parameters that can be used to prioritize genes for further study. These parameters include significant: (A) *cis*-eQTL; (B) correlation between the *cis* component of expression and the trait (TWAS); and (C) correlation of overall expression with the trait. C*is*-eQTL with *P* < 1e-4 are considered significant, and correlations with bicor *P* < 0.05 are considered significant. If a gene has a significant *cis*-eQTL in this locus, the most significant *cis*-eQTL SNP rsID and its base pair location, *cis*-eQTL *P* value, SNP weight, and SNP odds ratio are noted in columns J-N. If the *cis* component of expression of a gene is significantly correlated with liver TG levels, and the gene is thus a “TWAS gene” for liver TG, the TWAS SNP rsID, TWAS SNP base pair, TWAS correlation value, and TWAS *P* value are noted in columns O-R. Note: no genes in this locus were significant TWAS genes for liver iron. If total liver mRNA expression of a gene is significantly correlated with liver iron levels or liver TG levels, then the correlation and *P* value are noted in columns S-V. Finally, the liver mRNA expression raw transcript count minimum, maximum, mean, and median across all 114 mouse strains is noted in columns W-Z. Genes that passed expression cutoffs (expression values >0.1 transcripts per million (TPM) in at least 20% of samples and ≥ 6 reads in at least 20% of samples) are noted in column AA. The top 3 candidate genes for liver TG based on this analysis, the liver TG TWAS genes, are highlighted in yellow.

**Supplementary File 31.** Table of genes located within 1 Mb of the chromosome 11 GWAS peak for liver iron with data from some parameters that can be to prioritize genes for further study. These parameters include significant: (A) *cis*-eQTL; (B) correlation between the *cis* component of expression and the trait (TWAS); and(C) correlation of overall expression with the trait. C*is*-eQTL with *P* < 1e-4 are considered significant, and correlations with bicor *P* < 0.05 are considered significant. If a gene has a significant *cis*-eQTL in this region, the most significant SNP rsID, base pair, eQTL *P* value, SNP weight, and odds ratio are noted in columns I-M. If the *cis* component of expression of a gene is significantly correlated with liver iron levels, the TWAS SNP rsID, SNP base pair, TWAS correlation value, and TWAS *P* value are noted in columns N-Q. If total liver mRNA expression of a gene is significantly correlated with liver iron levels, then the correlation and *P* value are noted in columns R and S. Finally, the liver mRNA expression raw transcript count minimum, maximum, and mean across all 114 mouse strains is noted in columns T-V. Genes that passed expression cutoffs (expression values >0.1 transcripts per million (TPM) in at least 20% of samples and ≥ 6 reads in at least 20% of samples) are noted in column W. The top candidates, which have significant liver iron TWAS results, are highlighted in yellow.

**Figure 6-Source Data 1.** Table of SNPs that are significantly associated in *trans* (*trans*-eQTL *P* value < 1e-6) with the liver mRNA expression of at least one gene. For the purposes of this table, *trans* associations are defined as associations between a SNP and a gene that are located on different chromosomes. The SNP rsID, chromosome, and base pair location are noted. The number of genes associated in *trans* with each SNP in the iron overload HMDP in column D. For comparison purposes, the number of genes associated in *trans* with each SNP from another HMDP study, where male mice were fed a control chow diet, are noted in column E (Tuominen, Fuqua et al. 2021).

**Supplementary File 32.** Table of the top 15 *trans*-eQTL hotspots as ranked by number of significant *trans* associated genes (*trans*-eQTL *P* < 1e-6) in the liver mRNA expression dataset. For the purposes of this table, *trans* associations are defined as associations between a SNP and a gene that are located on different chromosomes. Hotspot SNPs associated with the same hotspot region (based on close location in the genome and strong overlap in the list of genes associated in *trans* with the SNP) are grouped together under one hotspot. Only hotspot SNPs ranked in the top 100 by the number of genes associated in *trans* are included in this table. In column E, any genes significantly associated with these hotspot SNPs in *cis* (*cis*-eQTL *P* < 1e-4 and with a gene start or stop within 2 Mb of the SNP) are noted, as they are top candidate genes whose expression may influence the expression of the *trans*-associated genes of this hotspot. A summary of the top Metascape pathway enrichment analysis terms for the genes associated in *trans* with each of the top 15 hotspots in the iron study are listed in column F.

**Supplementary File 33.** List of all genes whose liver mRNA expression is significantly associated in *trans* with a SNP. For the purposes of this list, *trans* associations are defined as associations between a SNP and a gene that are located on different chromosomes, and where the P value for the association is less than 0.05. Column A is an ordering column used to maintain order if the list is sorted. In column B, the rsIDs of SNPs with *trans* associated genes are noted in green, with the list of genes below. In column C, the number of genes associated in *trans* with each SNP is noted.

**Supplementary File 34.** Top Metascape enrichment analysis annotations, ranked by their - log_10_*P* values, for the top ∼500 genes by *P* value <u>positively</u> and significantly correlated (bicor *P* < 0.05) with liver TG across all 114 strains of mice. (***A***) Metascape pathway enrichments. (***B***) Metascape disease enrichments from DisGeNET (https://www.disgenet.org). (***C***) Metascape cell type enrichments. (***D***) Metascape transcription factor target enrichments (***E***) Metascape transcriptional regulation enrichments from TRRUST (http://www.grnpedia.org/trrust). Gene expression correlation with liver TG is given in Supplementary File 36.

**Supplementary File 35.** Top Metascape enrichment analysis annotations, ranked by their - log_10_*P* values, for the top ∼500 genes by *P* value <u>negatively</u> and significantly correlated (bicor *P* < 0.05) with liver TG across all 114 strains of mice. (***A***) Metascape pathway enrichments. (***B***) Metascape disease enrichments from DisGeNET (https://www.disgenet.org). (***C***) Metascape cell type enrichments. (***D***) Metascape transcription factor target enrichments (***E***) Metascape transcriptional regulation enrichments from TRRUST (http://www.grnpedia.org/trrust). Gene expression correlation with liver TG is given in Supplementary File 36.

**Supplementary File 36.** Liver mRNA expression correlations with liver iron and liver TG across the HMDP. The first tab gives correlations with liver iron, and the second tab gives correlations for liver TG.

**Supplementary File 37.** Table showing the differential expression of genes in the livers of the C57BL/6J Control study mice fed the 50 ppm iron sufficient control diet versus the 20k ppm iron loaded diet. N = 3 mice per diet group. Fold change, *P* values and FDR adjusted *P* values (*P* _adj_) for differential expression are noted. In column J, genes are ranked by increasing *P* value for differential expression. In column K, the ranking of genes by increasing *P* value for correlation with liver iron in the HMDP is given. The sum of these two ranks is given in column L. Genes with a low sum of ranks are more likely to be more differentially expressed in response to iron in C57BL/6J and to have liver mRNA expression levels more strongly correlated with liver iron levels across all 114 HMDP mouse strains.

**Supplementary File 38.** Differential gene mRNA expression analysis between livers from mice fed the 50 ppm iron sufficient control diet and those fed the 20k ppm high iron diet in the C57BL/6J Control study. (***A***) Volcano plot of the -log_10_*P* for differential expression versus the log_2_(fold change in gene expression). Genes with increased expression in mice on the high iron diet have positive log_2_(fold change) values. (***B*** and ***C***). Metascape pathway enrichment analysis pathways and -log_10_(enrichment *P*) for genes with *P*_adj_ < 0.05 that have decreased (***B***) or increased expression (***C***) in the 20k ppm high iron diet livers compared to the 50 ppm iron sufficient control diet livers.

**Figure 9-Source Data 1.** WGCNA co-expression data. Module eigengene values (first tab), module eigengene to trait bicor and P value (second tab), and the list of gene membership in each module (third tab) are given.

**Supplementary File 39.** WGCNA module membership of genes that were significantly differentially expressed (DE) (*P*_adj_ ≤ 0.05) between the 50 ppm iron sufficient control diet and 20k ppm high iron diet in the Control C57BL/6J study. The first table lists the genes, their WGCNA module membership, their DE P adj values and log 2 fold change in the C57BL/6J Control study, and the correlation (and P value) of their liver mRNA expression with liver iron across the HMDP. The table at the right gives the gene counts and percentages of DE and Fe correlated genes in each module.

**Supplementary File 40**. WGCNA module membership of differentially expressed and iron correlated genes. (***A***) The percentage of genes in each WGCNA module that were or were not significantly differentially expressed (DE) (*P*_adj_ ≤ 0.05) between the 50 ppm iron sufficient control diet and 20k ppm high iron diet in the Control C57BL/6J study. Negative DE genes were downregulated in response to the high iron diet, while positive DE genes were upregulated in response to the high iron diet. (***B***) The percentage of DE genes in each WGCNA module that were positively DE and positively correlated with liver iron across the HMDP, positively DE and negatively correlated with liver iron, negatively DE and positively correlated with liver iron, negatively DE and negatively correlated with liver iron, or DE but not correlated with liver iron. The grey module contains all the genes that were not clustered into any module (i.e. not co-expressed). The data in Supplementary File 39 corresponds with this figure.

**Supplementary File 41.** Key driver genes associated with liver TG identified by Mergeomics KDA. The key drivers (large nodes) and their top connected genes (small nodes) that belong to the TG-associated WGCNA blue module are shown.

## References

Ahmed, U., P. S. Latham and P. S. Oates (2012). “Interactions between hepatic iron and lipid metabolism with possible relevance to steatohepatitis.” World J Gastroenterol 18(34): 4651–4658.

Aigner, E., M. Strasser, H. Haufe, T. Sonnweber, F. Hohla, A. Stadlmayr, M. Solioz, H. Tilg, W. Patsch, G. Weiss, F. Stickel and C. Datz (2010). “A role for low hepatic copper concentrations in nonalcoholic Fatty liver disease.” Am J Gastroenterol 105(9): 1978–1985.

Aigner, E., I. Theurl, H. Haufe, M. Seifert, F. Hohla, L. Scharinger, F. Stickel, F. Mourlane, G. Weiss and C. Datz (2008). “Copper availability contributes to iron perturbations in human nonalcoholic fatty liver disease.” Gastroenterology 135(2): 680–688.

An, P., H. Wang, Q. Wu, J. Wang, Z. Xia, X. He, X. Wang, Y. Chen, J. Min and F. Wang (2018). “Smad7 deficiency decreases iron and haemoglobin through hepcidin up-regulation by multilayer compensatory mechanisms.” J Cell Mol Med 22(6): 3035–3044.

Anderson, G. J. and E. Bardou-Jacquet (2021). “Revisiting hemochromatosis: genetic vs. phenotypic manifestations.” Ann Transl Med 9(8): 731.

Anderson, G. J. and D. M. Frazer (2017). “Current understanding of iron homeostasis.” Am J Clin Nutr 106(Suppl 6): 1559s–1566s.

Angeli, J. P. F., F. P. Freitas, P. Nepachalovich, L. Puentes, O. Zilka, A. Inague, S. Lorenz, V. Kunz, H. Nehring, T. N. X. d. Silva, Z. Chen, S. Doll, W. Schmitz, P. Imming, S. Miyamoto, J. Klein-Seetharaman, L. Kumar, T. C. Genaro-Mattos, K. Mirnics, S. Meierjohann, M. Kroiss, I. Weigand, K. Bommert, R. Bargou, A. Garcia-Saez, D. Pratt, M. Fedorova, A. Wehmann, A. Horling, G. Bornkamm and M. Conrad (2021). 7-Dehydrocholesterol is an endogenous suppressor of ferroptosis, Research Square.

Ashraf, U. M., E. R. Sanchez and S. Kumarasamy (2019). “COUP-TFII revisited: Its role in metabolic gene regulation.” Steroids 141: 63–69.

Aydinok, Y., J. B. Porter, A. Piga, M. Elalfy, A. El-Beshlawy, Y. Kilinç, V. Viprakasit, A. Yesilipek, D. Habr, E. Quebe-Fehling and D. J. Pennell (2015). “Prevalence and distribution of iron overload in patients with transfusion-dependent anemias differs across geographic regions: results from the CORDELIA study.” Eur J Haematol 95(3): 244–253.

Bailey, A. P., G. Koster, C. Guillermier, E. M. Hirst, J. I. MacRae, C. P. Lechene, A. D. Postle and A. P. Gould (2015). “Antioxidant Role for Lipid Droplets in a Stem Cell Niche of Drosophila.” Cell 163(2): 340–353.

Barreau, P. B. and J. E. Buttery (1987). “The effect of the haematocrit value on the determination of glucose levels by reagent-strip methods.” Med J Aust 147(6): 286–288.

Belaidi, A. A. and A. I. Bush (2016). “Iron neurochemistry in Alzheimer’s disease and Parkinson’s disease: targets for therapeutics.” J Neurochem 139 Suppl 1: 179–197.

Bell, S., A. S. Rigas, M. K. Magnusson, E. Ferkingstad, E. Allara, G. Bjornsdottir, A. Ramond, E. Sørensen, G. H. Halldorsson, D. S. Paul, K. S. Burgdorf, H. P. Eggertsson, J. M. M. Howson, L. W. Thørner, S. Kristmundsdottir, W. J. Astle, C. Erikstrup, J. K. Sigurdsson, D. Vuckovic, K. M. Dinh, V. Tragante, P. Surendran, O. B. Pedersen, B. Vidarsson, T. Jiang, H. M. Paarup, P. T. Onundarson, P. Akbari, K. R. Nielsen, S. H. Lund, K. Juliusson, M. I. Magnusson, M. L. Frigge, A. Oddsson, I. Olafsson, S. Kaptoge, H. Hjalgrim, G. Runarsson, A. M. Wood, I. Jonsdottir, T. F. Hansen, O. Sigurdardottir, H. Stefansson, D. Rye, J. E. Peters, D. Westergaard, H. Holm, N. Soranzo, K. Banasik, G. Thorleifsson, W. H. Ouwehand, U. Thorsteinsdottir, D. J. Roberts, P. Sulem, A. S. Butterworth, D. F. Gudbjartsson, J. Danesh, S. Brunak, E. Di Angelantonio, H. Ullum and K. Stefansson (2021). “A genome-wide meta-analysis yields 46 new loci associating with biomarkers of iron homeostasis.” Commun Biol 4(1): 156.

Bellezza, I., R. Roberti, L. Gatticchi, R. Del Sordo, M. G. Rambotti, M. C. Marchetti, A. Sidoni and A. Minelli (2013). “A novel role for Tm7sf2 gene in regulating TNFα expression.” PLoS One 8(7): e68017.

Bellsmith, K. N., J. L. Dunaief, P. Yang, M. E. Pennesi, E. Davis, H. Hofkamp and B. J. Lujan (2020). “Bull’s eye maculopathy associated with hereditary hemochromatosis.” Am J Ophthalmol Case Rep 18: 100674.

Belyaeva, O. V., O. V. Korkina, A. V. Stetsenko and N. Y. Kedishvili (2008). “Human retinol dehydrogenase 13 (RDH13) is a mitochondrial short-chain dehydrogenase/reductase with a retinaldehyde reductase activity.” Febs j 275(1): 138–147.

Ben M’barek, K., D. Ajjaji, A. Chorlay, S. Vanni, L. Forêt and A. R. Thiam (2017). “ER Membrane Phospholipids and Surface Tension Control Cellular Lipid Droplet Formation.” Dev Cell 41(6): 591–604.e597.

Bennett, B. J., C. R. Farber, L. Orozco, H. M. Kang, A. Ghazalpour, N. Siemers, M. Neubauer, I. Neuhaus, R. Yordanova, B. Guan, A. Truong, W. P. Yang, A. He, P. Kayne, P. Gargalovic, T. Kirchgessner, C. Pan, L. W. Castellani, E. Kostem, N. Furlotte, T. A. Drake, E. Eskin and A. J. Lusis (2010). “A high-resolution association mapping panel for the dissection of complex traits in mice.” Genome Res 20(2): 281–290.

Bensaid, M., S. Fruchon, C. Mazères, S. Bahram, M. P. Roth and H. Coppin (2004). “Multigenic control of hepatic iron loading in a murine model of hemochromatosis.” Gastroenterology 126(5): 1400–1408.

Benyamin, B., T. Esko, J. S. Ried, A. Radhakrishnan, S. H. Vermeulen, M. Traglia, M. Gögele, D. Anderson, L. Broer, C. Podmore, J. Luan, Z. Kutalik, S. Sanna, P. van der Meer, T. Tanaka, F. Wang, H. J. Westra, L. Franke, E. Mihailov, L. Milani, J. Hälldin, J. Winkelmann, T. Meitinger, J. Thiery, A. Peters, M. Waldenberger, A. Rendon, J. Jolley, J. Sambrook, L. A. Kiemeney, F. C. Sweep, C. F. Sala, C. Schwienbacher, I. Pichler, J. Hui, A. Demirkan, A. Isaacs, N. Amin, M. Steri, G. Waeber, N. Verweij, J. E. Powell, D. R. Nyholt, A. C. Heath, P. A. Madden, P. M. Visscher, M. J. Wright, G. W. Montgomery, N. G. Martin, D. Hernandez, S. Bandinelli, P. van der Harst, M. Uda, P. Vollenweider, R. A. Scott, C. Langenberg, N. J. Wareham, C. van Duijn, J. Beilby, P. P. Pramstaller, A. A. Hicks, W. H. Ouwehand, K. Oexle, C. Gieger, A. Metspalu, C. Camaschella, D. Toniolo, D. W. Swinkels and J. B. Whitfield (2014). “Novel loci affecting iron homeostasis and their effects in individuals at risk for hemochromatosis.” Nat Commun 5: 4926.

Bloomer, S. A. and K. E. Brown (2019). “Iron-Induced Liver Injury: A Critical Reappraisal.” Int J Mol Sci 20(9).

Boccuto, L., K. Aoki, H. Flanagan-Steet, C. F. Chen, X. Fan, F. Bartel, M. Petukh, A. Pittman, R. Saul, A. Chaubey, E. Alexov, M. Tiemeyer, R. Steet and C. E. Schwartz (2014). “A mutation in a ganglioside biosynthetic enzyme, ST3GAL5, results in salt & pepper syndrome, a neurocutaneous disorder with altered glycolipid and glycoprotein glycosylation.” Hum Mol Genet 23(2): 418–433.

Bowe, J. E., Z. J. Franklin, A. C. Hauge-Evans, A. J. King, S. J. Persaud and P. M. Jones (2014). “Metabolic phenotyping guidelines: assessing glucose homeostasis in rodent models.” J Endocrinol 222(3): G13–25.

Britton, L. J., V. N. Subramaniam and D. H. Crawford (2016). “Iron and non-alcoholic fatty liver disease.” World J Gastroenterol 22(36): 8112–8122.

Brzostek-Racine, S., C. Gordon, S. Van Scoy and N. C. Reich (2011). “The DNA damage response induces IFN.” J Immunol 187(10): 5336–5345.

Buch, S., A. Sharma, E. Ryan, C. Datz, W. J. H. Griffiths, M. Way, T. W. M. Buckley, J. D. Ryan, S. Stewart, C. Wright, P. Dongiovanni, A. Fracanzani, J. Zwerina, U. Merle, K. H. Weiss, E. Aigner, E. Krones, C. Dejaco, J. Fischer, T. Berg, L. Valenti, H. Zoller, A. McQuillin, J. Hampe, F. Stickel and M. Y. Morgan (2021). “Variants in PCSK7, PNPLA3 and TM6SF2 are risk factors for the development of cirrhosis in hereditary haemochromatosis.” Aliment Pharmacol Ther 53(7): 830–843.

Cairo, G., D. Conte, L. Bianchi, M. Fraquelli and S. Recalcati (2001). “Reduced serum ceruloplasmin levels in hereditary haemochromatosis.” Br J Haematol 114(1): 226–229.

Cao, Y., Y. Wang, Z. Zhou, C. Pan, L. Jiang, Z. Zhou, Y. Meng, S. Charugundla, T. Li, H. Allayee, M. M. Seldin and A. J. Lusis (2022). “Liver-heart cross-talk mediated by coagulation factor XI protects against heart failure.” Science 377(6613): 1399–1406.

Cavey, T., M. Ropert, M. de Tayrac, E. Bardou-Jacquet, M. L. Island, P. Leroyer, C. Bendavid, P. Brissot and O. Loréal (2015). “Mouse genetic background impacts both on iron and non-iron metals parameters and on their relationships.” Biometals 28(4): 733–743.

Chacon, A. H., B. Morrison and S. Hu (2013). “Acquired hemochromatosis with pronounced pigment deposition of the upper eyelids.” J Clin Aesthet Dermatol 6(10): 44–46.

Chanas, S. A., Q. Jiang, M. McMahon, G. K. McWalter, L. I. McLellan, C. R. Elcombe, C. J. Henderson, C. R. Wolf, G. J. Moffat, K. Itoh, M. Yamamoto and J. D. Hayes (2002). “Loss of the Nrf2 transcription factor causes a marked reduction in constitutive and inducible expression of the glutathione S-transferase Gsta1, Gsta2, Gstm1, Gstm2, Gstm3 and Gstm4 genes in the livers of male and female mice.” Biochem J 365(Pt 2): 405–416.

Chen, H. J., M. Sugiyama, F. Shimokawa, M. Murakami, O. Hashimoto, T. Matsui and M. Funaba (2020). “Response to iron overload in cultured hepatocytes.” Sci Rep 10(1): 21184.

Chua, A. C., B. R. Klopcic, D. S. Ho, S. K. Fu, C. H. Forrest, K. D. Croft, J. K. Olynyk, I. C. Lawrance and D. Trinder (2013). “Dietary iron enhances colonic inflammation and IL-6/IL-11-Stat3 signaling promoting colonic tumor development in mice.” PLoS One 8(11): e78850.

Civelek, M. and A. J. Lusis (2014). “Systems genetics approaches to understand complex traits.” Nat Rev Genet 15(1): 34–48.

Collins, J. F. and G. J. Anderson (2012). Chapter 71 - Molecular Mechanisms of Intestinal Iron Transport. Physiology of the Gastrointestinal Tract (Fifth Edition). L. R. Johnson, F. K. Ghishan, J. D. Kaunitz et al. Boston, Academic Press: 1921–1947.

Crawford, D. H. G., D. G. F. Ross, L. A. Jaskowski, L. J. Burke, L. J. Britton, N. Musgrave, D. Briskey, G. Rishi, K. R. Bridle and V. N. Subramaniam (2021). “Iron depletion attenuates steatosis in a mouse model of non-alcoholic fatty liver disease: Role of iron-dependent pathways.” Biochim Biophys Acta Mol Basis Dis 1867(7): 166142.

Cui, X., B. Ma, Y. Wang, Y. Chen, C. Shen, Y. Kuang, J. Fei, L. Lu and Z. Wang (2019). “Rdh13 deficiency weakens carbon tetrachloride-induced liver injury by regulating Spot14 and Cyp2e1 expression levels.” Front Med 13(1): 104–111.

Curnock, R. and P. J. Cullen (2020). “Mammalian copper homeostasis requires retromer-dependent recycling of the high-affinity copper transporter 1.” J Cell Sci 133(16).

Das, S., S. Maji, Ruturaj, I. Bhattacharya, T. Saha, N. Naskar and A. Gupta (2020). “Retromer retrieves the Wilson disease protein ATP7B from endolysosomes in a copper-dependent manner.” J Cell Sci 133(24).

Datz, C., E. Müller and E. Aigner (2017). “Iron overload and non-alcoholic fatty liver disease.” Minerva Endocrinol 42(2): 173–183.

DeDiego, M. L., L. Martinez-Sobrido and D. J. Topham (2019). “Novel Functions of IFI44L as a Feedback Regulator of Host Antiviral Responses.” J Virol 93(21).

Ding, H., Q. Zhang, X. Yu, L. Chen, Z. Wang and J. Feng (2021). “Lipidomics reveals perturbations in the liver lipid profile of iron-overloaded mice.” Metallomics 13(10): mfab057.

Ding, J., M. Blencowe, T. Nghiem, S. M. Ha, Y. W. Chen, G. Li and X. Yang (2021). “Mergeomics 2.0: a web server for multi-omics data integration to elucidate disease networks and predict therapeutics.” Nucleic Acids Res 49(W1): W375–w387.

Dufies, M., A. Verbiest, L. S. Cooley, P. D. Ndiaye, X. He, N. Nottet, W. Souleyreau, A. Hagege, S. Torrino, J. Parola, S. Giuliano, D. Borchiellini, R. Schiappa, B. Mograbi, J. Zucman-Rossi, K. Bensalah, A. Ravaud, P. Auberger, A. Bikfalvi, E. Chamorey, N. Rioux-Leclercq, N. M. Mazure, B. Beuselinck, Y. Cao, J. C. Bernhard, D. Ambrosetti and G. Pagès (2021). “Plk1, upregulated by HIF-2, mediates metastasis and drug resistance of clear cell renal cell carcinoma.” Commun Biol 4(1): 166.

El Beshlawy, A., M. El Tagui, M. Hamdy, M. El Ghamrawy, K. A. Azim, D. Salem, F. Said, A. Samir, T. St Pierre and D. J. Pennell (2014). “Low prevalence of cardiac siderosis in heavily iron loaded Egyptian thalassemia major patients.” Ann Hematol 93(3): 375–379.

Elahi, S. and S. Mashhouri (2020). “Immunological consequences of extramedullary erythropoiesis: immunoregulatory functions of CD71(+) erythroid cells.” Haematologica 105(6): 1478–1483.

Eswarappa, M., C. Cantarelli and P. Cravedi (2021). “Erythropoietin in Lupus: Unanticipated Immune Modulating Effects of a Kidney Hormone.” Front Immunol 12: 639370.

Fernandez, M., J. Lokan, C. Leung and A. Grigg (2022). “A critical evaluation of the role of iron overload in fatty liver disease.” J Gastroenterol Hepatol.

Fiel, M. I. (2022). “Methods to determine hepatic iron content.” UpToDate Retrieved July 16, 2022.

Fjorback, A. W. and O. M. Andersen (2012). “SorLA is a molecular link for retromer-dependent sorting of the Amyloid precursor protein.” Commun Integr Biol 5(6): 616–619.

Folch, J., M. Lees and G. H. Sloane Stanley (1957). “A simple method for the isolation and purification of total lipides from animal tissues.” J Biol Chem 226(1): 497–509.

Fox, J. G. (2014). The mouse in biomedical research: Normative biology, husbandry, and models, Academic Press.

Galaris, D., A. Barbouti and K. Pantopoulos (2019). “Iron homeostasis and oxidative stress: An intimate relationship.” Biochim Biophys Acta Mol Cell Res 1866(12): 118535.

Garcia-Bermudez, J., L. Baudrier, E. C. Bayraktar, Y. Shen, K. La, R. Guarecuco, B. Yucel, D. Fiore, B. Tavora, E. Freinkman, S. H. Chan, C. Lewis, W. Min, G. Inghirami, D. M. Sabatini and K. Birsoy (2019). “Squalene accumulation in cholesterol auxotrophic lymphomas prevents oxidative cell death.” Nature 567(7746): 118–122.

Goh, J. B., D. F. Wallace, W. Hong and V. N. Subramaniam (2015). “Endofin, a novel BMP-SMAD regulator of the iron-regulatory hormone, hepcidin.” Sci Rep 5: 13986.

Gouya, L., F. Muzeau, A. M. Robreau, P. Letteron, E. Couchi, S. Lyoumi, J. C. Deybach, H. Puy, R. Fleming, P. Demant, C. Beaumont and B. Grandchamp (2007). “Genetic study of variation in normal mouse iron homeostasis reveals ceruloplasmin as an HFE-hemochromatosis modifier gene.” Gastroenterology 132(2): 679–686.

Graham, R. M., A. C. Chua, K. W. Carter, R. D. Delima, D. Johnstone, C. E. Herbison, M. J. Firth, R. O’Leary, E. A. Milward, J. K. Olynyk and D. Trinder (2010). “Hepatic iron loading in mice increases cholesterol biosynthesis.” Hepatology 52(2): 462–471.

Gregg, X. T., V. Reddy and J. T. Prchal (2002). “Copper deficiency masquerading as myelodysplastic syndrome.” Blood 100(4): 1493–1495.

Gujja, P., D. R. Rosing, D. J. Tripodi and Y. Shizukuda (2010). “Iron overload cardiomyopathy: better understanding of an increasing disorder.” J Am Coll Cardiol 56(13): 1001–1012.

Ha, J. H., C. Doguer and J. F. Collins (2017). “Consumption of a High-Iron Diet Disrupts Homeostatic Regulation of Intestinal Copper Absorption in Adolescent Mice.” Am J Physiol Gastrointest Liver Physiol 313(4): G535–g360.

Ha, J. H., C. Doguer, S. R. L. Flores, T. Wang and J. F. Collins (2018). “Progressive Increases in Dietary Iron Are Associated with the Emergence of Pathologic Disturbances of Copper Homeostasis in Growing Rats.” J Nutr 148(3): 373–378.

Ha, J. H., C. Doguer, X. Wang, S. R. Flores and J. F. Collins (2016). “High-Iron Consumption Impairs Growth and Causes Copper-Deficiency Anemia in Weanling Sprague-Dawley Rats.” PLoS One 11(8): e0161033.

Hariz, A. and P. Bhattacharya (Updated 2022 Sep 26). Megaloblastic Anemia. StatPearls [Internet]. Treasure Island (FL), StatPearls Publishing.

Hart, K. M., T. Fabre, J. C. Sciurba, R. L. Gieseck, 3rd, L. A. Borthwick, K. M. Vannella, T. H. Acciani, R. de Queiroz Prado, R. W. Thompson, S. White, G. Soucy, M. Bilodeau, T. R. Ramalingam, J. R. Arron, N. H. Shoukry and T. A. Wynn (2017). “Type 2 immunity is protective in metabolic disease but exacerbates NAFLD collaboratively with TGF-β.” Sci Transl Med 9(396).

Hassannia, B., P. Vandenabeele and T. Vanden Berghe (2019). “Targeting Ferroptosis to Iron Out Cancer.” Cancer Cell 35(6): 830–849.

He, X., A. W. Ashbrook, Y. Du, J. Wu, H. H. Hoffmann, C. Zhang, L. Xia, Y. C. Peng, K. C. Tumas, B. K. Singh, C. F. Qi, T. G. Myers, C. A. Long, C. Liu, R. Wang, C. M. Rice and X. Z. Su (2020). “RTP4 inhibits IFN-I response and enhances experimental cerebral malaria and neuropathology.” Proc Natl Acad Sci U S A 117(32): 19465–19474.

Hedrick, C. C., L. W. Castellani, C. H. Warden, D. L. Puppione and A. J. Lusis (1993). “Influence of mouse apolipoprotein A-II on plasma lipoproteins in transgenic mice.” J Biol Chem 268(27): 20676–20682.

Hsu, C. C., N. H. Senussi, K. Y. Fertrin and K. V. Kowdley (2022). “Iron overload disorders.” Hepatology Communications 6(8): 1842–1854.

Huang, X., Z. Shi, W. Wang, J. Bai, Z. Chen, J. Xu, D. Zhang and S. Fu (2007). “Identification and characterization of a novel protein ISOC2 that interacts with p16INK4a.” Biochem Biophys Res Commun 361(2): 287–293.

Huang, Y., J. Hale, Y. Wang, W. Li, S. Zhang, J. Zhang, H. Zhao, X. Guo, J. Liu, H. Yan, K. Yazdanbakhsh, G. Huang, C. D. Hillyer, N. Mohandas, L. Chen, L. Sun and X. An (2018). “SF3B1 deficiency impairs human erythropoiesis via activation of p53 pathway: implications for understanding of ineffective erythropoiesis in MDS.” J Hematol Oncol 11(1): 19.

Issitt, T., E. Bosseboeuf, N. De Winter, N. Dufton, G. Gestri, V. Senatore, A. Chikh, A. M. Randi and C. Raimondi (2019). “Neuropilin-1 Controls Endothelial Homeostasis by Regulating Mitochondrial Function and Iron-Dependent Oxidative Stress.” iScience 11: 205–223.

Jeidane, S., M. P. Scott-Boyer, N. Tremblay, S. Cardin, S. Picard, M. Baril, D. Lamarre and C. F. Deschepper (2016). “Association of a Network of Interferon-Stimulated Genes with a Locus Encoding a Negative Regulator of Non-conventional IKK Kinases and IFNB1.” Cell Rep 17(2): 425–435.

Jiang, Y., Y. Tang, C. Hoover, Y. Kondo, D. Huang, D. Restagno, B. Shao, L. Gao, J. Michael McDaniel, M. Zhou, R. Silasi-Mansat, S. McGee, M. Jiang, X. Bai, F. Lupu, C. Ruan, J. D. Marth, D. Wu, Y. Han and L. Xia (2021). “Kupffer cell receptor CLEC4F is important for the destruction of desialylated platelets in mice.” Cell Death Differ 28(11): 3009–3021.

Jones, B. C., J. L. Beard, J. N. Gibson, E. L. Unger, R. P. Allen, K. A. McCarthy and C. J. Earley (2007). “Systems genetic analysis of peripheral iron parameters in the mouse.” Am J Physiol Regul Integr Comp Physiol 293(1): R116–124.

Jones, L. C., J. L. Beard and B. C. Jones (2008). “Genetic analysis reveals polygenic influences on iron, copper, and zinc in mouse hippocampus with neurobiological implications.” Hippocampus 18(4): 398–410.

Kanamori, Y., M. Murakami, M. Sugiyama, O. Hashimoto, T. Matsui and M. Funaba (2019). “Hepcidin and IL-1β.” Vitam Horm 110: 143–156.

Kane, S. F., C. Roberts and R. Paulus (2021). “Hereditary Hemochromatosis: Rapid Evidence Review.” Am Fam Physician 104(3): 263–270.

Kang, W., A. Barad, A. G. Clark, Y. Wang, X. Lin, Z. Gu and K. O. O’Brien (2021). “Ethnic Differences in Iron Status.” Adv Nutr 12(5): 1838–1853.

Kautz, L., D. Meynard, A. Monnier, V. Darnaud, R. Bouvet, R. H. Wang, C. Deng, S. Vaulont, J. Mosser, H. Coppin and M. P. Roth (2008). “Iron regulates phosphorylation of Smad1/5/8 and gene expression of Bmp6, Smad7, Id1, and Atoh8 in the mouse liver.” Blood 112(4): 1503–1509.

Khetarpal, S. A., C. Vitali, M. G. Levin, D. Klarin, J. Park, A. Pampana, J. S. Millar, T. Kuwano, D. Sugasini, P. V. Subbaiah, J. T. Billheimer, P. Natarajan and D. J. Rader (2021). “Endothelial lipase mediates efficient lipolysis of triglyceride-rich lipoproteins.” PLoS Genet 17(9): e1009802.

Knopp, J. L., L. Holder-Pearson and J. G. Chase (2019). “Insulin Units and Conversion Factors: A Story of Truth, Boots, and Faster Half-Truths.” J Diabetes Sci Technol 13(3): 597–600.

Kouroumalis, E., I. Tsomidis and A. Voumvouraki (2023). “Iron as a therapeutic target in chronic liver disease.” World J Gastroenterol 29(4): 616–655.

Kowdley, K. V., K. E. Brown, J. Ahn and V. Sundaram (2019). “ACG Clinical Guideline: Hereditary Hemochromatosis.” Am J Gastroenterol 114(8): 1202–1218.

Kremastinos, D. T. and D. Farmakis (2011). “Iron overload cardiomyopathy in clinical practice.” Circulation 124(20): 2253–2263.

Lainé, F., M. Ropert, C. L. Lan, O. Loréal, E. Bellissant, C. Jard, M. Pouchard, A. Le Treut and P. Brissot (2002). “Serum ceruloplasmin and ferroxidase activity are decreased in HFE C282Y homozygote male iron-overloaded patients.” J Hepatol 36(1): 60–65.

Langfelder, P. and S. Horvath (2008). “WGCNA: an R package for weighted correlation network analysis.” BMC Bioinformatics 9: 559.

Lee, Y. K., J. E. Park, M. Lee and J. P. Hardwick (2018). “Hepatic lipid homeostasis by peroxisome proliferator-activated receptor gamma 2.” Liver Res 2(4): 209–215.

Li, J., L. A. Lange, Q. Duan, Y. Lu, A. B. Singleton, A. B. Zonderman, M. K. Evans, Y. Li, H. A. Taylor, M. S. Willis, M. Nalls, J. G. Wilson and E. M. Lange (2015). “Genome-wide admixture and association study of serum iron, ferritin, transferrin saturation and total iron binding capacity in African Americans.” Hum Mol Genet 24(2): 572–581.

Li, Y., D. Song, Y. Song, L. Zhao, N. Wolkow, J. W. Tobias, W. Song and J. L. Dunaief (2015). “Iron-induced Local Complement Component 3 (C3) Up-regulation via Non-canonical Transforming Growth Factor (TGF)-β Signaling in the Retinal Pigment Epithelium.” J Biol Chem 290(19): 11918–11934.

Liao, M., J. Shi, L. Huang, Y. Gao, A. Tan, C. Wu, Z. Lu, X. Yang, S. Zhang, Y. Hu, X. Qin, J. Li, G. Chen, J. Xu, Z. Mo and H. Zhang (2014). “Genome-wide association study identifies variants in PMS1 associated with serum ferritin in a Chinese population.” PLoS One 9(8): e105844.

Lippert, C., J. Listgarten, Y. Liu, C. M. Kadie, R. I. Davidson and D. Heckerman (2011). “FaST linear mixed models for genome-wide association studies.” Nat Methods 8(10): 833–835.

Liu, Y., N. Basty, B. Whitcher, J. D. Bell, E. P. Sorokin, N. van Bruggen, E. L. Thomas and M. Cule (2021). “Genetic architecture of 11 organ traits derived from abdominal MRI using deep learning.” Elife 10.

Love, M. I., W. Huber and S. Anders (2014). “Moderated estimation of fold change and dispersion for RNA-seq data with DESeq2.” Genome Biology 15(12): 550.

Lusis, A. J., M. M. Seldin, H. Allayee, B. J. Bennett, M. Civelek, R. C. Davis, E. Eskin, C. R. Farber, S. Hui, M. Mehrabian, F. Norheim, C. Pan, B. Parks, C. D. Rau, D. J. Smith, T. Vallim, Y. Wang and J. Wang (2016). “The Hybrid Mouse Diversity Panel: a resource for systems genetics analyses of metabolic and cardiovascular traits.” J Lipid Res 57(6): 925–942.

Ma, F., B. K. Fuqua, Y. Hasin, C. Yukhtman, C. D. Vulpe, A. J. Lusis and M. Pellegrini (2019). “A comparison between whole transcript and 3’ RNA sequencing methods using Kapa and Lexogen library preparation methods.” BMC Genomics 20(1): 9.

Martin, S., M. Cule, N. Basty, J. Tyrrell, R. N. Beaumont, A. R. Wood, T. M. Frayling, E. Sorokin, B. Whitcher, Y. Liu, J. D. Bell, E. L. Thomas and H. Yaghootkar (2021). “Genetic Evidence for Different Adiposity Phenotypes and Their Opposing Influences on Ectopic Fat and Risk of Cardiometabolic Disease.” Diabetes 70(8): 1843–1856.

McGarry, M. P., C. A. Protheroe and J. J. Lee (2010). Mouse hematology: A laboratory manual. New York, NY, Cold Spring Harbor Laboratory Press.

McLachlan, S., S. M. Lee, T. M. Steele, P. L. Hawthorne, M. A. Zapala, E. Eskin, N. J. Schork, G. J. Anderson and C. D. Vulpe (2011). “In silico QTL mapping of basal liver iron levels in inbred mouse strains.” Physiol Genomics 43(3): 136–147.

McLachlan, S., K. E. Page, S. M. Lee, A. Loguinov, E. Valore, S. T. Hui, G. Jung, J. Zhou, A. J. Lusis, B. Fuqua, T. Ganz, E. Nemeth and C. D. Vulpe (2017). “Hamp1 mRNA and plasma hepcidin levels are influenced by sex and strain but do not predict tissue iron levels in inbred mice.” Am J Physiol Gastrointest Liver Physiol 313(5): G511–g523.

McLaren, C. E., C. P. Garner, C. C. Constantine, S. McLachlan, C. D. Vulpe, B. M. Snively, V. R. Gordeuk, D. A. Nickerson, J. D. Cook, C. Leiendecker-Foster, K. B. Beckman, J. H. Eckfeldt, L. F. Barcellos, J. A. Murray, P. C. Adams, R. T. Acton, A. A. Killeen and G. D. McLaren (2011). “Genome-wide association study identifies genetic loci associated with iron deficiency.” PLoS One 6(3): e17390.

Meng, Y., S. Heybrock, D. Neculai and P. Saftig (2020). “Cholesterol Handling in Lysosomes and Beyond.” Trends Cell Biol 30(6): 452–466.

Milet, J., V. Dehais, C. Bourgain, A. M. Jouanolle, A. Mosser, M. Perrin, J. Morcet, P. Brissot, V. David, Y. Deugnier and J. Mosser (2007). “Common variants in the BMP2, BMP4, and HJV genes of the hepcidin regulation pathway modulate HFE hemochromatosis penetrance.” Am J Hum Genet 81(4): 799–807.

Miller, C. G. and E. E. Schmidt (2020). “Sulfur Metabolism Under Stress.” Antioxid Redox Signal 33(16): 1158–1173.

Møhlenberg, M., E. Terczynska-Dyla, K. L. Thomsen, J. George, M. Eslam, H. Grønbæk and R. Hartmann (2019). “The role of IFN in the development of NAFLD and NASH.” Cytokine 124: 154519.

Moksnes, M. R., S. E. Graham, K. H. Wu, A. F. Hansen, S. A. Gagliano Taliun, W. Zhou, K. Thorstensen, L. G. Fritsche, D. Gill, A. Mason, F. Cucca, D. Schlessinger, G. R. Abecasis, S. Burgess, B. O. Åsvold, J. B. Nielsen, K. Hveem, C. J. Willer and B. M. Brumpton (2022). “Genome-wide meta-analysis of iron status biomarkers and the effect of iron on all-cause mortality in HUNT.” Commun Biol 5(1): 591.

Muchenditsi, A., C. C. Talbot, Jr., A. Gottlieb, H. Yang, B. Kang, T. Boronina, R. Cole, L. Wang, S. Dev, J. P. Hamilton and S. Lutsenko (2021). “Systemic deletion of Atp7b modifies the hepatocytes’ response to copper overload in the mouse models of Wilson disease.” Sci Rep 11(1): 5659.

Mylonis, I., H. Sembongi, C. Befani, P. Liakos, S. Siniossoglou and G. Simos (2012). “Hypoxia causes triglyceride accumulation by HIF-1-mediated stimulation of lipin 1 expression.” J Cell Sci 125(Pt 14): 3485–3493.

Nishina, T., Y. Deguchi, D. Ohshima, W. Takeda, M. Ohtsuka, S. Shichino, S. Ueha, S. Yamazaki, M. Kawauchi, E. Nakamura, C. Nishiyama, Y. Kojima, S. Adachi-Akahane, M. Hasegawa, M. Nakayama, M. Oshima, H. Yagita, K. Shibuya, T. Mikami, N. Inohara, K. Matsushima, N. Tada and H. Nakano (2021). “Interleukin-11-expressing fibroblasts have a unique gene signature correlated with poor prognosis of colorectal cancer.” Nat Commun 12(1): 2281.

O’Dushlaine, C., M. Germino, N. Verweij, J. B. Nielsen, A. Yadav, C. Benner, J. D. Backman, N. Lin, G. R. Abecasis, A. Baras, M. A. Ferreira, L. A. Lotta, J. R. Walls, P. Parasoglou and J. L. Marchini (2021). “Genome-wide association study of liver fat, iron, and extracellular fluid fraction in the UK Biobank.” medRxiv: 2021.2010.2025.21265127.

Oexle, K., J. S. Ried, A. A. Hicks, T. Tanaka, C. Hayward, M. Bruegel, M. Gögele, P. Lichtner, B. Müller-Myhsok, A. Döring, T. Illig, C. Schwienbacher, C. Minelli, I. Pichler, G. M. Fiedler, J. Thiery, I. Rudan, A. F. Wright, H. Campbell, L. Ferrucci, S. Bandinelli, P. P. Pramstaller, H. E. Wichmann, C. Gieger, J. Winkelmann and T. Meitinger (2011). “Novel association to the proprotein convertase PCSK7 gene locus revealed by analysing soluble transferrin receptor (sTfR) levels.” Hum Mol Genet 20(5): 1042–1047.

Okada, M. and K. Ye (2009). “Nuclear phosphoinositide signaling regulates messenger RNA export.” RNA Biol 6(1): 12–16.

Okuno, Y., A. Fukuhara, E. Hashimoto, H. Kobayashi, S. Kobayashi, M. Otsuki and I. Shimomura (2018). “Oxidative Stress Inhibits Healthy Adipose Expansion Through Suppression of SREBF1-Mediated Lipogenic Pathway.” Diabetes 67(6): 1113–1127.

Parisinos, C. A., H. R. Wilman, E. L. Thomas, M. Kelly, R. C. Nicholls, J. McGonigle, S. Neubauer, A. D. Hingorani, R. S. Patel, H. Hemingway, J. D. Bell, R. Banerjee and H. Yaghootkar (2020). “Genome-wide and Mendelian randomisation studies of liver MRI yield insights into the pathogenesis of steatohepatitis.” J Hepatol 73(2): 241–251.

Pavlovic, Z. and M. Bakovic (2013). “Regulation of Phosphatidylethanolamine Homeostasis&#8212;The Critical Role of CTP:Phosphoethanolamine Cytidylyltransferase (Pcyt2).” Int J Mol Sci 14(2): 2529–2550.

Pelucchi, S., R. Mariani, S. Calza, A. L. Fracanzani, G. L. Modignani, F. Bertola, F. Busti, P. Trombini, M. Fraquelli, G. L. Forni, D. Girelli, S. Fargion, C. Specchia and A. Piperno (2012). “CYBRD1 as a modifier gene that modulates iron phenotype in HFE p.C282Y homozygous patients.” Haematologica 97(12): 1818–1825.

Perng, Y. C. and D. J. Lenschow (2018). “ISG15 in antiviral immunity and beyond.” Nat Rev Microbiol 16(7): 423–439.

Phillips-Krawczak, C. A., A. Singla, P. Starokadomskyy, Z. Deng, D. G. Osborne, H. Li, C. J. Dick, T. S. Gomez, M. Koenecke, J. S. Zhang, H. Dai, L. F. Sifuentes-Dominguez, L. N. Geng, S. H. Kaufmann, M. Y. Hein, M. Wallis, J. McGaughran, J. Gecz, B. Sluis, D. D. Billadeau and E. Burstein (2015). “COMMD1 is linked to the WASH complex and regulates endosomal trafficking of the copper transporter ATP7A.” Mol Biol Cell 26(1): 91–103.

Pierson, H., A. Muchenditsi, B. E. Kim, M. Ralle, N. Zachos, D. Huster and S. Lutsenko (2018). “The Function of ATPase Copper Transporter ATP7B in Intestine.” Gastroenterology 154(1): 168–180.e165.

Pierson, H., H. Yang and S. Lutsenko (2019). “Copper Transport and Disease: What Can We Learn from Organoids?” Annu Rev Nutr 39: 75–94.

Podszun, M. C., A. S. Alawad, S. Lingala, N. Morris, W. A. Huang, S. Yang, M. Schoenfeld, A. Rolt, R. Ouwerkerk, K. Valdez, R. Umarova, Y. Ma, S. Z. Fatima, D. D. Lin, L. S. Mahajan, N. Samala, P. C. Violet, M. Levine, R. Shamburek, A. M. Gharib, D. E. Kleiner, H. M. Garraffo, H. Cai, P. J. Walter and Y. Rotman (2020). “Vitamin E treatment in NAFLD patients demonstrates that oxidative stress drives steatosis through upregulation of de-novo lipogenesis.” Redox Biol 37: 101710.

Prasnicka, A., H. Lastuvkova, F. Alaei Faradonbeh, J. Cermanova, M. Hroch, J. Mokry, E. Dolezelova, P. Pavek, K. Zizalova, L. Vitek, P. Nachtigal and S. Micuda (2019). “Iron overload reduces synthesis and elimination of bile acids in rat liver.” Sci Rep 9(1): 9780.

Pressly, J. D., M. Z. Gurumani, J. T. Varona Santos, A. Fornoni, S. Merscher and H. Al-Ali (2022). “Adaptive and maladaptive roles of lipid droplets in health and disease.” Am J Physiol Cell Physiol 322(3): C468–c481.

Protchenko, O., E. Baratz, S. Jadhav, F. Li, M. Shakoury-Elizeh, O. Gavrilova, M. C. Ghosh, J. E. Cox, J. A. Maschek, V. A. Tyurin, Y. Y. Tyurina, H. Bayir, A. T. Aron, C. J. Chang, V. E. Kagan and C. C. Philpott (2021). “Iron Chaperone Poly rC Binding Protein 1 Protects Mouse Liver From Lipid Peroxidation and Steatosis.” Hepatology 73(3): 1176–1193.

Pruim, R. J., R. P. Welch, S. Sanna, T. M. Teslovich, P. S. Chines, T. P. Gliedt, M. Boehnke, G. R. Abecasis and C. J. Willer (2010). “LocusZoom: regional visualization of genome-wide association scan results.” Bioinformatics 26(18): 2336–2337.

Radio, F. C., S. Majore, C. Aurizi, F. Sorge, G. Biolcati, S. Bernabini, I. Giotti, F. Torricelli, D. Giannarelli, C. De Bernardo and P. Grammatico (2015). “Hereditary hemochromatosis type 1 phenotype modifiers in Italian patients. The controversial role of variants in HAMP, BMP2, FTL and SLC40A1 genes.” Blood Cells Mol Dis 55(1): 71–75.

Raffield, L. M., T. Louie, T. Sofer, D. Jain, E. Ipp, K. D. Taylor, G. J. Papanicolaou, L. Avilés-Santa, L. A. Lange, C. C. Laurie, M. P. Conomos, T. A. Thornton, Y. I. Chen, Q. Qi, S. Cotler, B. Thyagarajan, N. Schneiderman, J. I. Rotter, A. P. Reiner and H. J. Lin (2017). “Genome-wide association study of iron traits and relation to diabetes in the Hispanic Community Health Study/Study of Latinos (HCHS/SOL): potential genomic intersection of iron and glucose regulation?” Hum Mol Genet 26(10): 1966–1978.

Rametta, R., A. L. Fracanzani, S. Fargion and P. Dongiovanni (2020). “Dysmetabolic Hyperferritinemia and Dysmetabolic Iron Overload Syndrome (DIOS): Two Related Conditions or Different Entities?” Curr Pharm Des 26(10): 1025–1035.

Rau, C. D., B. Parks, Y. Wang, E. Eskin, P. Simecek, G. A. Churchill and A. J. Lusis (2015). “High-Density Genotypes of Inbred Mouse Strains: Improved Power and Precision of Association Mapping.” G3 (Bethesda) 5(10): 2021–2026.

Richardson, T. G., E. Sanderson, T. M. Palmer, M. Ala-Korpela, B. A. Ference, G. Davey Smith and M. V. Holmes (2020). “Evaluating the relationship between circulating lipoprotein lipids and apolipoproteins with risk of coronary heart disease: A multivariable Mendelian randomisation analysis.” PLoS Med 17(3): e1003062.

Romero, A. R., A. Mu and J. S. Ayres (2022). “Adipose triglyceride lipase mediates lipolysis and lipid mobilization in response to iron-mediated negative energy balance.” iScience 25(3): 103941.

Sahoo, S., D. Singh, P. Chakraborty and M. K. Jolly (2020). “Emergent Properties of the HNF4α-PPARγ Network May Drive Consequent Phenotypic Plasticity in NAFLD.” J Clin Med 9(3).

Sakaue, S., M. Kanai, Y. Tanigawa, J. Karjalainen, M. Kurki, S. Koshiba, A. Narita, T. Konuma, K. Yamamoto, M. Akiyama, K. Ishigaki, A. Suzuki, K. Suzuki, W. Obara, K. Yamaji, K. Takahashi, S. Asai, Y. Takahashi, T. Suzuki, N. Shinozaki, H. Yamaguchi, S. Minami, S. Murayama, K. Yoshimori, S. Nagayama, D. Obata, M. Higashiyama, A. Masumoto, Y. Koretsune, K. Ito, C. Terao, T. Yamauchi, I. Komuro, T. Kadowaki, G. Tamiya, M. Yamamoto, Y. Nakamura, M. Kubo, Y. Murakami, K. Yamamoto, Y. Kamatani, A. Palotie, M. A. Rivas, M. J. Daly, K. Matsuda and Y. Okada (2021). “A cross-population atlas of genetic associations for 220 human phenotypes.” Nat Genet 53(10): 1415–1424.

Sandling, J. K., P. Pucholt, L. Hultin Rosenberg, F. H. G. Farias, S. V. Kozyrev, M. L. Eloranta, A. Alexsson, M. Bianchi, L. Padyukov, C. Bengtsson, R. Jonsson, R. Omdal, B. A. Lie, L. Massarenti, R. Steffensen, M. A. Jakobsen, S. T. Lillevang, K. Lerang, Ø. Molberg, A. Voss, A. Troldborg, S. Jacobsen, A. C. Syvänen, A. Jönsen, I. Gunnarsson, E. Svenungsson, S. Rantapää-Dahlqvist, A. A. Bengtsson, C. Sjöwall, D. Leonard, K. Lindblad-Toh and L. Rönnblom (2021). “Molecular pathways in patients with systemic lupus erythematosus revealed by gene-centred DNA sequencing.” Ann Rheum Dis 80(1): 109–117.

Santer, D. M., A. E. Wiedeman, T. H. Teal, P. Ghosh and K. B. Elkon (2012). “Plasmacytoid dendritic cells and C1q differentially regulate inflammatory gene induction by lupus immune complexes.” J Immunol 188(2): 902–915.

Seeßle, J., H. Gan-Schreier, M. Kirchner, W. Stremmel, W. Chamulitrat and U. Merle (2020). “Plasma Lipidome, PNPLA3 polymorphism and hepatic steatosis in hereditary hemochromatosis.” BMC Gastroenterol 20(1): 230.

Seldin, M., X. Yang and A. J. Lusis (2019). “Systems genetics applications in metabolism research.” Nat Metab 1(11): 1038–1050.

Sengsuk, C., O. Tangvarasittichai, P. Chantanaskulwong, A. Pimanprom, S. Wantaneeyawong, A. Choowet and S. Tangvarasittichai (2014). “Association of Iron Overload with Oxidative Stress, Hepatic Damage and Dyslipidemia in Transfusion-Dependent β-Thalassemia/HbE Patients.” Indian J Clin Biochem 29(3): 298–305.

Shen, T., Y. Li, L. Yang, X. Xu, F. Liang, S. Liang, G. Ba, F. Xue and Q. Fu (2012). “Upregulation of Polo-like kinase 2 gene expression by GATA-1 acetylation in human osteosarcoma MG-63 cells.” Int J Biochem Cell Biol 44(2): 423–429.

Shu, L., Y. Zhao, Z. Kurt, S. G. Byars, T. Tukiainen, J. Kettunen, L. D. Orozco, M. Pellegrini, A. J. Lusis, S. Ripatti, B. Zhang, M. Inouye, V. P. Mäkinen and X. Yang (2016). “Mergeomics: multidimensional data integration to identify pathogenic perturbations to biological systems.” BMC Genomics 17(1): 874.

Sinnott-Armstrong, N., Y. Tanigawa, D. Amar, N. Mars, C. Benner, M. Aguirre, G. R. Venkataraman, M. Wainberg, H. M. Ollila, T. Kiiskinen, A. S. Havulinna, J. P. Pirruccello, J. Qian, A. Shcherbina, F. Rodriguez, T. L. Assimes, V. Agarwala, R. Tibshirani, T. Hastie, S. Ripatti, J. K. Pritchard, M. J. Daly and M. A. Rivas (2021). “Genetics of 35 blood and urine biomarkers in the UK Biobank.” Nat Genet 53(2): 185–194.

Sirlin, C. B. and S. B. Reeder (2010). “Magnetic resonance imaging quantification of liver iron.” Magn Reson Imaging Clin N Am 18(3): 359–381, ix.

Spracklen, C. N., M. Horikoshi, Y. J. Kim, K. Lin, F. Bragg, S. Moon, K. Suzuki, C. H. T. Tam, Y. Tabara, S. H. Kwak, F. Takeuchi, J. Long, V. J. Y. Lim, J. F. Chai, C. H. Chen, M. Nakatochi, J. Yao, H. S. Choi, A. K. Iyengar, H. J. Perrin, S. M. Brotman, M. van de Bunt, A. L. Gloyn, J. E. Below, M. Boehnke, D. W. Bowden, J. C. Chambers, A. Mahajan, M. I. McCarthy, M. C. Y. Ng, L. E. Petty, W. Zhang, A. P. Morris, L. S. Adair, M. Akiyama, Z. Bian, J. C. N. Chan, L. C. Chang, M. L. Chee, Y. I. Chen, Y. T. Chen, Z. Chen, L. M. Chuang, S. Du, P. Gordon-Larsen, M. Gross, X. Guo, Y. Guo, S. Han, A. G. Howard, W. Huang, Y. J. Hung, M. Y. Hwang, C. M. Hwu, S. Ichihara, M. Isono, H. M. Jang, G. Jiang, J. B. Jonas, Y. Kamatani, T. Katsuya, T. Kawaguchi, C. C. Khor, K. Kohara, M. S. Lee, N. R. Lee, L. Li, J. Liu, A. O. Luk, J. Lv, Y. Okada, M. A. Pereira, C. Sabanayagam, J. Shi, D. M. Shin, W. Y. So, A. Takahashi, B. Tomlinson, F. J. Tsai, R. M. van Dam, Y. B. Xiang, K. Yamamoto, T. Yamauchi, K. Yoon, C. Yu, J. M. Yuan, L. Zhang, W. Zheng, M. Igase, Y. S. Cho, J. I. Rotter, Y. X. Wang, W. H. H. Sheu, M. Yokota, J. Y. Wu, C. Y. Cheng, T. Y. Wong, X. O. Shu, N. Kato, K. S. Park, E. S. Tai, F. Matsuda, W. P. Koh, R. C. W. Ma, S. Maeda, I. Y. Millwood, J. Lee, T. Kadowaki, R. G. Walters, B. J. Kim, K. L. Mohlke and X. Sim (2020). “Identification of type 2 diabetes loci in 433,540 East Asian individuals.” Nature 582(7811): 240–245.

Sylvers-Davie, K. L. and B. S. J. Davies (2021). “Regulation of lipoprotein metabolism by ANGPTL3, ANGPTL4, and ANGPTL8.” Am J Physiol Endocrinol Metab 321(4): E493–e508.

Taher, A. T., D. J. Weatherall and M. D. Cappellini (2018). “Thalassaemia.” The Lancet 391(10116): 155–167.

Thingholm, L. B., M. C. Rühlemann, M. Koch, B. Fuqua, G. Laucke, R. Boehm, C. Bang, E. A. Franzosa, M. Hübenthal, A. Rahnavard, F. Frost, J. Lloyd-Price, M. Schirmer, A. J. Lusis, C. D. Vulpe, M. M. Lerch, G. Homuth, T. Kacprowski, C. O. Schmidt, U. Nöthlings, T. H. Karlsen, W. Lieb, M. Laudes, A. Franke and C. Huttenhower (2019). “Obese Individuals with and without Type 2 Diabetes Show Different Gut Microbial Functional Capacity and Composition.” Cell Host Microbe 26(2): 252–264.e210.

Thoß, M., K. C. Luzynski, M. Ante, I. Miller and D. J. Penn (2015). “Major urinary protein (MUP) profiles show dynamic changes rather than individual ’barcode’ signatures.” Front Ecol Evol 3.

Torti, S. V., D. H. Manz, B. T. Paul, N. Blanchette-Farra and F. M. Torti (2018). “Iron and Cancer.” Annu Rev Nutr 38: 97–125.

Tuominen, I., B. K. Fuqua, C. Pan, N. Renaud, K. Wroblewski, M. Civelek, K. Clerkin, A. Asaryan, S. G. Haroutunian, J. Loureiro, J. Borawski, G. Roma, J. Knehr, W. Carbone, S. French, B. W. Parks, S. T. Hui, M. Mehrabian, C. Magyar, R. M. Cantor, C. Ukomadu, A. J. Lusis and S. W. Beaven (2021). “The Genetic Architecture of Carbon Tetrachloride-Induced Liver Fibrosis in Mice.” Cell Mol Gastroenterol Hepatol 11(1): 199–220.

Turner, S. D. (2018). “qqman: an R package for visualizing GWAS results using Q-Q and manhattan plots.” Journal of Open Source Software 3(25).

van Bokhoven, M. A., C. T. van Deursen and D. W. Swinkels (2011). “Diagnosis and management of hereditary haemochromatosis.” Bmj 342: c7251.

Virbasius, C. M., S. Wagner and M. R. Green (1999). “A human nuclear-localized chaperone that regulates dimerization, DNA binding, and transcriptional activity of bZIP proteins.” Mol Cell 4(2): 219–228.

Vujkovic, M., J. M. Keaton, J. A. Lynch, D. R. Miller, J. Zhou, C. Tcheandjieu, J. E. Huffman, T. L. Assimes, K. Lorenz, X. Zhu, A. T. Hilliard, R. L. Judy, J. Huang, K. M. Lee, D. Klarin, S. Pyarajan, J. Danesh, O. Melander, A. Rasheed, N. H. Mallick, S. Hameed, I. H. Qureshi, M. N. Afzal, U. Malik, A. Jalal, S. Abbas, X. Sheng, L. Gao, K. H. Kaestner, K. Susztak, Y. V. Sun, S. L. DuVall, K. Cho, J. S. Lee, J. M. Gaziano, L. S. Phillips, J. B. Meigs, P. D. Reaven, P. W. Wilson, T. L. Edwards, D. J. Rader, S. M. Damrauer, C. J. O’Donnell, P. S. Tsao, K. M. Chang, B. F. Voight and D. Saleheen (2020). “Discovery of 318 new risk loci for type 2 diabetes and related vascular outcomes among 1.4 million participants in a multi-ancestry meta-analysis.” Nat Genet 52(7): 680–691.

Wang, H., X. Cui, Q. Gu, Y. Chen, J. Zhou, Y. Kuang, Z. Wang and X. Xu (2012). “Retinol dehydrogenase 13 protects the mouse retina from acute light damage.” Mol Vis 18: 1021–1030.

Wang, Q., D. Zhou, M. Wang, M. Zhu, P. Chen, H. Li, M. Lu, X. Zhang, X. Shen, T. Liu and L. Chen (2022). “A Novel Non-Invasive Approach Based on Serum Ceruloplasmin for Identifying Non-Alcoholic Steatohepatitis Patients in the Non-Diabetic Population.” Front Med (Lausanne) 9: 900794.

Wang, T., P. Xiang, J. H. Ha, X. Wang, C. Doguer, S. R. L. Flores, Y. J. Kang and J. F. Collins (2018). “Copper supplementation reverses dietary iron overload-induced pathologies in mice.” J Nutr Biochem 59: 56–63.

Warnick, G. R. (1986). “Enzymatic methods for quantification of lipoprotein lipids.” Methods Enzymol 129: 101–123.

Widjaja, A. A., B. K. Singh, E. Adami, S. Viswanathan, J. Dong, G. A. D’Agostino, B. Ng, W. W. Lim, J. Tan, B. S. Paleja, M. Tripathi, S. Y. Lim, S. G. Shekeran, S. P. Chothani, A. Rabes, M. Sombetzki, E. Bruinstroop, L. P. Min, R. A. Sinha, S. Albani, P. M. Yen, S. Schafer and S. A. Cook (2019). “Inhibiting Interleukin 11 Signaling Reduces Hepatocyte Death and Liver Fibrosis, Inflammation, and Steatosis in Mouse Models of Nonalcoholic Steatohepatitis.” Gastroenterology 157(3): 777–792.e714.

Williams, A. L., V. Khadka, M. Tang, A. Avelar, K. J. Schunke, M. Menor and R. V. Shohet (2018). “HIF1 mediates a switch in pyruvate kinase isoforms after myocardial infarction.” Physiol Genomics 50(7): 479–494.

Wilman, H. R., C. A. Parisinos, N. Atabaki-Pasdar, M. Kelly, E. L. Thomas, S. Neubauer, A. Mahajan, A. D. Hingorani, R. S. Patel, H. Hemingway, P. W. Franks, J. D. Bell, R. Banerjee and H. Yaghootkar (2019). “Genetic studies of abdominal MRI data identify genes regulating hepcidin as major determinants of liver iron concentration.” J Hepatol 71(3): 594–602.

Wu, J., Y. Wang, R. Jiang, R. Xue, X. Yin, M. Wu and Q. Meng (2021). “Ferroptosis in liver disease: new insights into disease mechanisms.” Cell Death Discov 7(1): 276.

Xiao, Q. and V. M. Lauschke (2021). “The prevalence, genetic complexity and population-specific founder effects of human autosomal recessive disorders.” NPJ Genom Med 6(1): 41.

Xu, L., Z. Korade and N. A. Porter (2010). “Oxysterols from free radical chain oxidation of 7-dehydrocholesterol: product and mechanistic studies.” J Am Chem Soc 132(7): 2222–2232.

Yamada, N., T. Karasawa, T. Komada, T. Matsumura, C. Baatarjav, J. Ito, K. Nakagawa, D. Yamamuro, S. Ishibashi, K. Miura, N. Sata and M. Takahashi (2022). “DHCR7 as a novel regulator of ferroptosis in hepatocytes.” bioRxiv: 2022.2006.2015.496212.

Yang, J., S. H. Lee, M. E. Goddard and P. M. Visscher (2011). “GCTA: a tool for genome-wide complex trait analysis.” Am J Hum Genet 88(1): 76–82.

You, M., A. Jogasuria, K. Lee, J. Wu, Y. Zhang, Y. K. Lee and P. Sadana (2017). “Signal Transduction Mechanisms of Alcoholic Fatty Liver Disease: Emer ging Role of Lipin-1.” Curr Mol Pharmacol 10(3): 226–236.

Zein, A. A., R. Kaur, T. O. K. Hussein, G. A. Graf and J. Y. Lee (2019). “ABCG5/G8: a structural view to pathophysiology of the hepatobiliary cholesterol secretion.” Biochem Soc Trans 47(5): 1259–1268.

Zhang, L., X. Huang, Z. Meng, B. Dong, S. Shiah, D. D. Moore and W. Huang (2009). “Significance and mechanism of CYP7a1 gene regulation during the acute phase of liver regeneration.” Mol Endocrinol 23(2): 137–145.

Zhang, P. and K. Reue (2017). “Lipin proteins and glycerolipid metabolism: Roles at the ER membrane and beyond.” Biochim Biophys Acta Biomembr 1859(9 Pt B): 1583–1595.

Zhang, Z. H. and G. L. Song (2021). “Roles of Selenoproteins in Brain Function and the Potential Mechanism of Selenium in Alzheimer’s Disease.” Front Neurosci 15: 646518.

Zhao, M., J. A. Laissue and A. Zimmermann (1997). “Hepatocyte apoptosis in hepatic iron overload diseases.” Histol Histopathol 12(2): 367–374.

Zheng, Y., T. Y. Lin, G. Lee, M. N. Paddock, J. Momb, Z. Cheng, Q. Li, D. L. Fei, B. D. Stein, S. Ramsamooj, G. Zhang, J. Blenis and L. C. Cantley (2018). “Mitochondrial One-Carbon Pathway Supports Cytosolic Folate Integrity in Cancer Cells.” Cell 175(6): 1546–1560.e1517.

Zhou, Y., B. Zhou, L. Pache, M. Chang, A. H. Khodabakhshi, O. Tanaseichuk, C. Benner and S. K. Chanda (2019). “Metascape provides a biologist-oriented resource for the analysis of systems-level datasets.” Nat Commun 10(1): 1523.

Zhou, Z., T. J. Ye, G. Bonavita, M. Daniels, N. Kainrad, A. Jogasuria and M. You (2019). “Adipose-Specific Lipin-1 Overexpression Renders Hepatic Ferroptosis and Exacerbates Alcoholic Steatohepatitis in Mice.” Hepatol Commun 3(5): 656–669.

